# The long isoform of ZAP coordinates multiple enzymes to mediate complete decay of target transcripts

**DOI:** 10.1101/2025.04.28.650959

**Authors:** Clément R. Bouton, Grega Gimpelj Domjanič, María José Lista, Rui Pedro Galão, Thomas Courty, Piotr Kwiatkowski, Harry D. Wilson, Peter W. S. Hill, Hannah E. Mischo, Anob M. Chakrabarti, Mario Poljak, Jernej Ule, Stuart J. D. Neil, Chad M. Swanson

**Author notes:** These authors contributed equally (Clément R. Bouton, Grega Gimpelj Domjanič, María José Lista). Correspondence: Jernej Ule, Stuart Neil, Chad Swanson.

## Abstract

Zinc-finger Antiviral Protein (ZAP)-mediated RNA decay (ZMD) restricts replication of viruses containing CpG dinucleotide clusters. However, why ZAP isoforms differ in antiviral activity and how they recruit cofactors to mediate RNA decay is unclear. Therefore, we determined the ordered events of the ZMD pathway. The long ZAP isoform preferentially binds viral RNA and has distinct binding motifs compared to the short isoform. The endoribonuclease KHNYN then cleaves viral RNA at positions of ZAP binding. The 5’ cleavage fragment undergoes TUT4/TUT7-mediated 3’ uridylation and degradation by DIS3L2. The 3’ cleavage fragment is degraded by XRN1. ZAP and TRIM25 interact with KHNYN, TUT7, DIS3L2 and XRN1 in a RNase-resistant manner. Viral infection promotes the interaction between TRIM25 with these enzymes, leading to viral RNA decay while also decreasing the abundance of cellular transcripts. Overall, the long isoform of ZAP recruits key enzymes to assemble an RNA decay complex on viral RNA.

## INTRODUCTION

A critical component of the innate immune response is the ability to distinguish between self and non-self nucleic acids. The best characterized non-self RNA detection systems include RIG-I, MDA5, PKR, OAS1-3 and ZBP1, which promote signaling that leads to antiviral gene expression, cap-dependent translation inhibition, global RNA degradation and cell death^1^. These pattern recognition receptors (PRRs) bind double-stranded RNA (dsRNA), which acts as the pathogen-associated molecular pattern (PAMP). In contrast, antiviral proteins that bind single-stranded RNA (ssRNA) often directly restrict viral replication by inhibiting viral RNA translation or targeting it for decay^2^. Zinc-finger Antiviral Protein (ZAP, also known as PARP13 or ZC3HAV1) is a paradigmatic example of a ssRNA-binding protein critical for the cell autonomous innate immune response^3,4^.

ZAP is encoded by an interferon-stimulated gene (ISG) that restricts diverse viruses including retroviruses, alphaviruses, Ebola virus and human cytomegalovirus^5^. It has two primary mechanisms for inhibiting viral infection: promoting viral RNA decay^3^ and inhibiting viral mRNA translation^6–8^. Several principles for the composition of ZAP-response elements (ZREs) have been determined from synonymous genome recoding of multiple viruses and structural/biochemical studies. CpG recoding can sensitize diverse viruses to ZAP-mediated restriction^9–15^, though UpA recoding has also been reported to sensitize viral RNA to ZAP^11,13,16^. ZAP-mediated restriction requires the presence of at least 15 CpGs, optimally separated by ∼12-30 nucleotides (nt)^17^, indicating that multivalency is required for substrate recognition. Many mammalian RNA viruses have low CpG abundance, likely due to evolutionary pressure from ZAP^9,18^. The RNA binding domain (RBD) in ZAP contains four zinc finger motifs (ZnF1-4) and directly binds a single CpG dinucleotide in pocket formed by ZnF2^3,10,12,19,20^. In addition to the CpG binding site, RNA base-specific contacts between ZnF3 and ZnF4 have been identified and C(n7)G(n)CG has been proposed as an optimal binding motif^12^. However, some regions of viral RNA appear to be more sensitive to CpG-mediated ZAP restriction for unknown reasons and high CpG abundance sometimes does not sensitize viral RNA to ZAP^14,21–25^. ZAP also regulates cellular transcript expression^26–29^ and this, in addition to CpG DNA methylation^30^, has been proposed to contribute to the selective pressure against CpG dinucleotide abundance in cellular genes^28,31,32^. However, little is known about the composition of ZREs in cellular transcripts.

ZAP is the core of a multicomponent antiviral system. It has no enzymatic activity and acts as a scaffold that recognizes target transcripts to recruit cofactors that inhibit translation or mediate RNA decay^4^. The original model for ZAP-mediated RNA decay (ZMD) was that it recruited enzymes that mediate decapping, deadenylation and exonucleolytic degradation (Figure 1A)^33–36^. Specifically, this model proposed that ZAP directly interacts with DDX17 to recruit DCP1A/DCP2 to mediate decapping and subsequent 5’-3’ decay via XRN1^34^. ZAP was also proposed to directly interact with PARN and RNA exosome subunits to mediate deadenylation and 3’-5’ decay^33,34^. In addition, DHX30 was proposed to promote ZAP activity^37^. Subsequently, ZAP was shown to interact with the E3 ubiquitin ligase TRIM25^38,39^ and the endoribonuclease KHNYN^40^. This led to an endonucleolytic decay model in which ZAP recruits KHNYN to internally cleave viral RNA and this is promoted by TRIM25 through an unclear mechanism^4,40–42^ (Figure 1B). However, how KHNYN cleavage products are degraded is unclear in this model. Neither model has been rigorously tested *in cellulo* and whether the two models can be reconciled is unclear. The KHNYN paralog N4BP1 has also been implicated in ZMD and these proteins have at least partially redundant activity for some transcripts^41,43,44^. Overall, the relative contribution of exonucleolytic versus endonucleolytic RNA decay machinery and how they act in an ordered pathway remain poorly defined.

**Figure 1:**
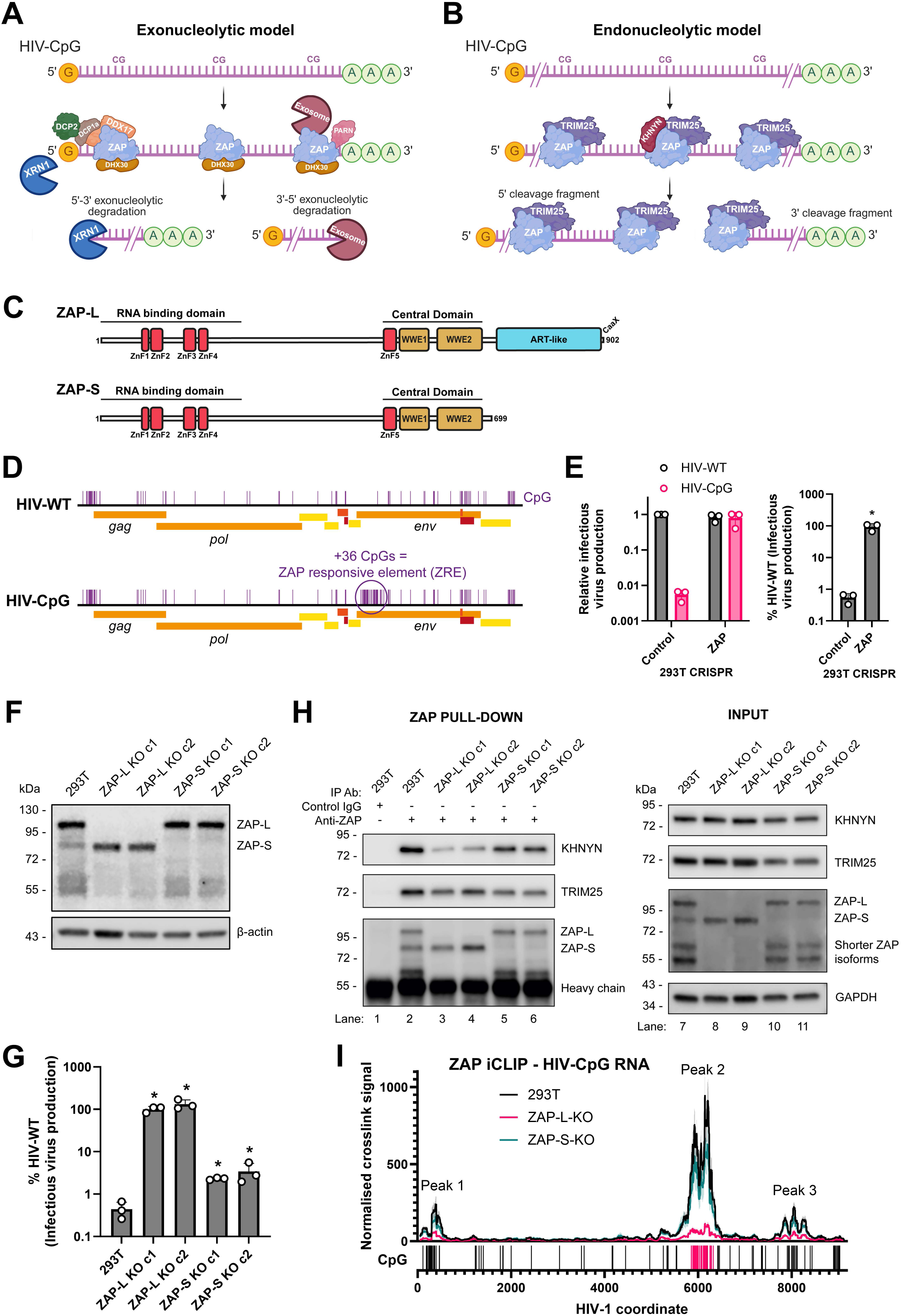
ZAP isoforms differ in their activity to restrict HIV-CpG and interact with KHNYN and viral RNA. **(A)** Schematic representation of the exonucleolytic model for ZMD. Created in BioRender. Bouton, C. (2025) https://BioRender.com/b8wrgls **(B)** Schematic representation of the endonucleolytic model for ZMD. Created in BioRender. Bouton, C. (2025) https://BioRender.com/lpjkyv4 **(C)** Schematic representation of ZAP’s main isoforms, ZAP-L and ZAP-S. ZnF: zinc finger. ART-like: ADP-ribosyltransferase-like domain. CaaX: CaaX box that mediates S-farnesylation. **(D)** Genomic organization and localization of CpG dinucleotides of HIV-WT and HIV-CpG. **(E)** Relative infectious HIV production (left) in HEK293T CRISPR control and ZAP cells, and HIV-CpG production expressed as % infectious HIV-WT production (right). Error bars represent standard deviation. *p <0.05 calculated by one-way ANOVA on log-transformed data, compared to HIV-CpG production in HEK293T CRISPR control cells. **(F)** Immunoblotting of uninfected isoform-specific HEK293T CRISPR ZAP cells. **(G)** Infectious HIV-CpG production, expressed as % infectious HIV-WT production, in isoform-specific HEK293T CRISPR ZAP cells. Error bars represent standard deviation. *p <0.05 calculated by one-way ANOVA on log-transformed data, compared to HIV-CpG production in HEK293T cells. **(H)** Immunoblotting of ZAP coimmunoprecipitations and corresponding input lysates obtained from isoform-specific HEK293T CRISPR ZAP cells at steady state level. The blots are representative of three biological replicates. **(I)** Normalized ZAP iCLIP signal mapped to HIV-CpG gRNA in isoform-specific HEK293T CRISPR ZAP cells. The average signal of three independent experiments and the SEM (shading) are plotted. Localization of CpG dinucleotides on HIV-CpG gRNA are indicated by vertical bars. Pink bars represent the 36 added CpG dinucleotides in HIV-CpG.

There are several ZAP isoforms, with ZAP-long (ZAP-L) and ZAP-short (ZAP-S) being the most abundant^3,45,46^ (Figure 1C). ZAP-S expression is induced by type I interferon while ZAP-L is constitutively expressed^46,47^, though both isoforms are induced by interferon γ^48^. Both ZAP-S and ZAP-L contain the RBD and a central domain that binds ADP-ribose^45,49,50^. ZAP-L also contains a catalytically inactive ADP-ribosyltransferase-like (ART-like) domain and a C-terminal CaaX box that mediates S-farnesylation^45,51^. This post-translational modification localizes ZAP-L to the cytoplasmic endomembrane system^27,51–53^. ZAP isoforms have been proposed to have different functions^47,52^ and, while ZAP-L has been reported to have higher antiviral activity against several viruses than ZAP-S^45,46,52,53^, the mechanism for this is unknown.

There are multiple therapeutic applications arising from understanding the targeting specificity of the ZAP antiviral system and how it mediates ZMD. TRIM25, ZAP, KHNYN and N4BP1 target mRNAs delivered by lipid nanoparticles when they do not contain N^1^-methylpseudouridine modifications, highlighting its importance for therapeutic RNA design^54^. Synonymous CpG-recoding of ZAP-resistant viral genomes has been shown to be a novel way to develop live-attenuated viral vaccines^13–15,17^. Because ZAP expression is downregulated in some types of cancer, ZAP-sensitive viruses have been proposed to be used as oncolytic viruses because they would preferentially replicate in tumors with low ZAP expression compared to healthy tissue^55^.

In this study, we characterized the ZMD pathway to determine how ZAP interacts with multiple enzymes to mediate viral RNA decay. Cells genetically engineered to only express ZAP-L or ZAP-S show that, even though both isoforms contain the RBD, ZAP-L is the predominant isoform that inhibits retroviral replication due to increased binding to viral RNA and KHNYN. We then show that ZAP and TRIM25 interact with multiple RNA decay enzymes including KHNYN, XRN1, TUT7 and DIS3L2 and order these enzymes into a decay pathway. *In cellulo* identification of KHNYN cleavage sites shows that KHNYN mediates endonucleolytic cleavage of the viral RNA in regions with high levels of ZAP binding. The 5’ cleavage fragment is 3’ uridylated by the paralogs TUT4/TUT7 and then degraded by DIS3L2. XRN1 degrades the 3’ cleavage fragment. Viral infection increases binding of TRIM25 to several components of this pathway and promotes regulation of cellular genes by ZAP, TRIM25 and KHNYN. Overall, ZAP acts as a core scaffold for several enzymes to mediate ZMD.

## RESULTS

### Differential antiviral activity between ZAP isoforms correlates with binding viral RNA and KHNYN

ZAP restricts a wide range of viruses^5^. For this study, we used HIV-1 as a molecularly tractable model virus to define the ordered events of the ZMD pathway. Wild-type HIV-1 (HIV-WT) is substantially depleted in CpG dinucleotides (Figure 1D) and the paucity of these is due to strong selective pressure, likely from ZAP^9,32,56,57^. Mutations that introduce a CpG in HIV-1 have a substantial fitness cost and the abundance of CpGs in HIV-1 *env* is linked to disease progression^24,58,59^. Because HIV-1 is substantially depleted in CpGs, these can be introduced through synonymous genome recoding to create HIV-CpG, which contains a potent ZRE in *env* (Figure 1D)^9,40^. This allows isogenic viruses to be analyzed under conditions in which ZAP restricts HIV-CpG but not HIV-WT (Figure 1E). This restriction can be quantified, showing that HIV-CpG infectious virus production is ∼1% of HIV-WT (Figure 1E). Importantly, both viruses produce similar levels of infectious virus when ZAP is knocked out. These paired viruses allow components of the ZAP antiviral system to be characterized under conditions in which the *cis*-acting ZRE is either absent or present in the virus, allowing ZMD substrate specificity to be determined in each experiment.

The relative contribution of the two primary ZAP isoforms (Figure 1C) to ZMD is controversial. While several papers have reported that ZAP-L is more active than ZAP-S, others have proposed that ZAP-S, or shorter fragments of ZAP, are sufficient for antiviral activity^24,41,45,46,51,52,60^. However, many of these experiments were performed by overexpressing ZAP isoforms in wild-type cells or reconstituting ZAP isoforms in CRISPR knockout cells, often by transient transfection, leading to non-physiological protein levels. Furthermore, the mechanism by which ZAP-L may be more active than ZAP-S is unclear. Therefore, we used a set of previously described HEK293T cells that were CRISPR-engineered to express only ZAP-L or ZAP-S by targeting splice sites and poly(A) sites in the endogenous *ZC3HAV1* locus^27^. These cells allow the antiviral activity of ZAP-L or ZAP-S to be characterized without ectopic expression (Figure 1F). ZAP-L knockout eliminated almost all ZAP antiviral activity against HIV-CpG while ZAP-S knockout had only a minor effect (Figure 1G and S1A-B), demonstrating that, in this system, ZAP-L is the essential mediator of viral restriction.

ZAP-L has previously been reported to interact more strongly with TRIM25 and KHNYN than ZAP-S, though one or more of these proteins was often overexpressed in the experimental conditions^38,53,61^. To determine if ZAP-L and ZAP-S preferentially interact with TRIM25 or KHNYN when all the proteins are expressed from their endogenous locus, we performed co-immunoprecipitation experiments in the wild-type, ZAP-L and ZAP-S knockout cell lines. While ZAP-L or ZAP-S knockout did not substantially affect the interaction between ZAP and TRIM25, ZAP-L knockout led to decreased ZAP interaction with KHNYN (Figure 1H and S1C).

Because ZAP directly binds KHNYN through its N-terminal RBD^41,42^, both ZAP isoforms contain the KHNYN interaction domain. A possible mechanism for why depletion of ZAP-L affected the ZAP-KHNYN interaction more than ZAP-S depletion may be that ZAP-L assembles on the ZRE more efficiently than ZAP-S. To test this hypothesis, we used iCLIP^62^ to quantify the binding of each isoform to HIV-WT and HIV-CpG RNA (Figure 1I and S1D-F). Three peaks were identified (Figure S1G), with the first at the 5’ end of the viral genome, the second in the 5’ region of *env* covering the sequence that has been CpG recoded, and the third in the 3’ region of *env*. Peak 1 is ∼250 nt wide in the 5’ leader region and *gag*, which contain a packaging element^63^. Peak 2, encompassing the CpG recoded region in HIV-CpG *env* (Figure 1D, 475 nt long^40^), is ∼800 nt long, indicating that ZAP binding extends beyond the recoded region. Peak 3 is ∼800 nt long and covers splice acceptor 7 (SA7, nt 7904-7915)^64^. As expected, ZAP binding is most pronounced in Peak 2, covering the engineered ZRE. Consistent with ZAP oligomerization and the hypothesis that multivalent binding of ZAP to multiple CpGs is required for its antiviral activity^17,41,65^, the iCLIP data implies assembly of ZAP molecules that cover wide, non-contiguous regions on the viral RNA. Interestingly, the CpG clusters underlying Peak 1 and Peak 3, which are present in both HIV-WT and HIV-CpG, show higher levels of binding in HIV-CpG than HIV-WT RNA (Figure S1D and S1H), indicating that high ZAP binding to the engineered region promotes additional ZAP recruitment at non-contiguous regions in the viral RNA containing clustered CpGs. Of note, the thresholds used for this peak identification were then used for all subsequent ZAP peak calling. In cells depleted for ZAP-S, high levels of ZAP-L binding to HIV-CpG RNA were observed (Figure 1I, S1F and S1I). By contrast, in ZAP-L knockout cells, ZAP-S showed low levels of RNA binding, correlating with its low antiviral activity (Figure 1I, S1E and S1I). In sum, ZAP-L is the primary antiviral isoform against HIV-CpG due to increased binding to HIV-CpG RNA and a stronger interaction with KHNYN.

### ZAP-L and ZAP-S bind different motifs and regulate transcripts from different genes

A potential evolutionary scenario for ZAP-regulated cellular transcripts is that ZAP evolved to restrict viral replication and was then exapted to control cellular transcripts containing similar sequences to the targeted viruses^66^. To determine why ZAP-L may bind HIV-CpG RNA better than ZAP-S, we analyzed the iCLIP data for ZAP binding to cellular transcripts. Motif analysis using PEKA^67^ identified different motifs for ZAP-L and ZAP-S (Figure 2A), with a UpA upstream of the CpG for 12 of the top 14 ZAP-L motifs, often in a UACG or UANCG context (Figure S2A). This may at least partially explain why UpA recoding of viral open reading frames can make a virus ZAP sensitive^11,13,16^.

**Figure 2:**
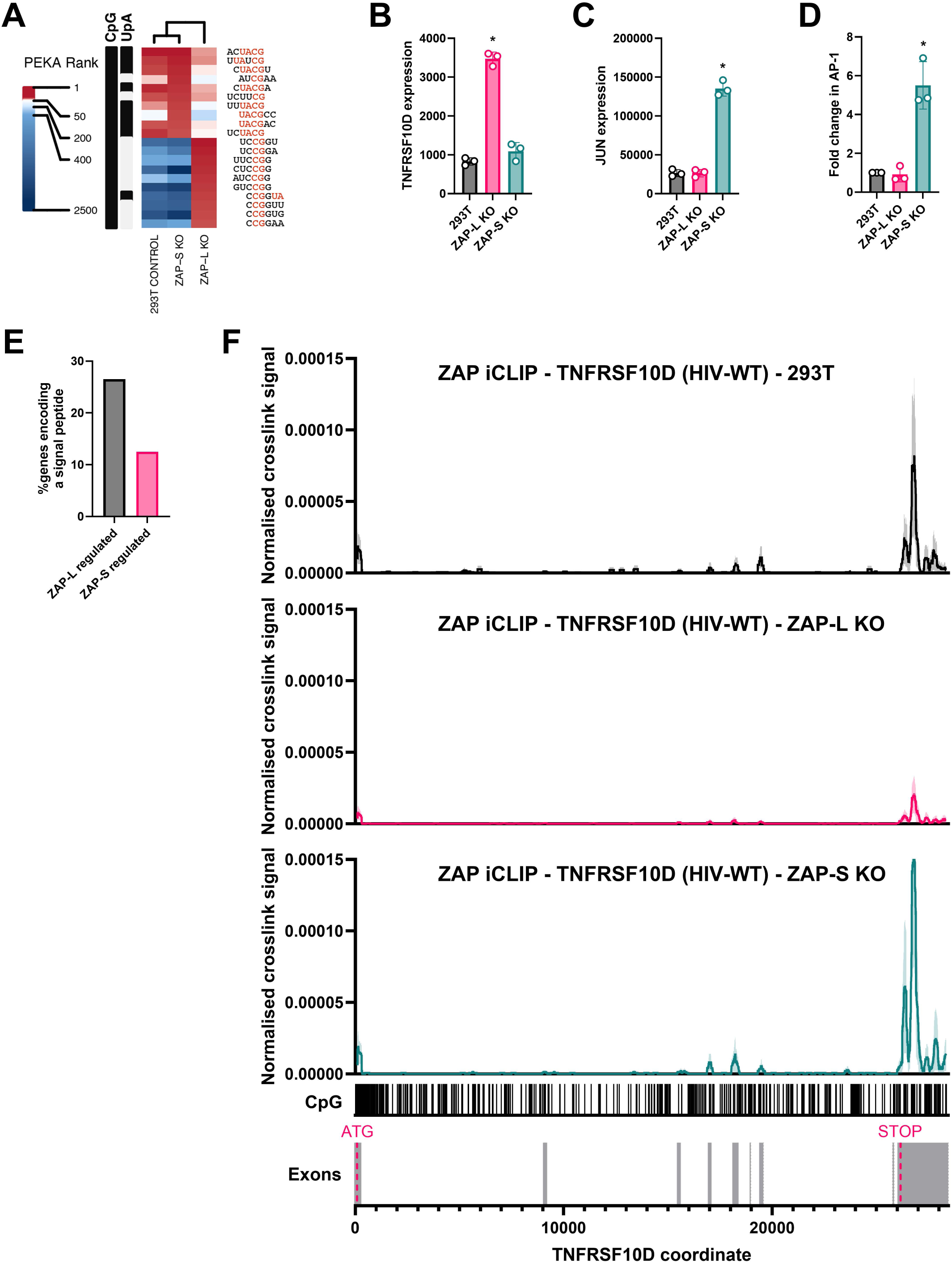
ZAP-L and ZAP-S bind different CpG-rich motifs and target different cellular transcripts. **(A)** Heatmap for the top 10 PEKA motifs from ZAP-S knockout or ZAP-L knockout cell iCLIP experiments shown for HEK293T, ZAP-S knockout and ZAP-L knockout cells. **(B)** Normalized DESeq2 counts for TNFRSF10D from poly(A) RNA-seq in HEK293T, ZAP-L and ZAP-S knockout cells. Error bars represent standard deviation. *adjp <0.05 calculated by DESeq2, compared to HEK293T cells. **(C)** Normalized DESeq2 counts for JUN from poly(A) RNA-seq in HEK293T, ZAP-L and ZAP-S knockout cells. Error bars represent standard deviation. *adjp <0.05 calculated by DESeq2, compared to HEK293T cells. **(D)** Normalized AP-1 reporter activity in HEK293T, ZAP-L and ZAP-S knockout cells. Error bars represent standard deviation. *p <0.05 calculated by one-way ANOVA on log-transformed data, compared to HEK293T cells. **(E)** % of genes specifically regulated by ZAP-L or ZAP-S predicted to contain a signal peptide by SignalP - 6.0^125^. **(F)** Normalized crosslink signal for ZAP iCLIP on TNFRSF10D in HEK293T, ZAP-L KO and ZAP-S KO cells. The average signal of three independent experiments and the SEM (shading) are plotted. The start and stop codon locations are indicated in pink.

We then analyzed whether ZAP-L and ZAP-S regulated different cellular transcripts using poly(A) RNA-seq (Table S1) and identified 156 and 240 genes encoding transcripts specifically regulated by ZAP-L or ZAP-S, respectively, in HIV-WT infected cells. Gene ontology (GO) analysis indicated that these are enriched for different function/localization (Figure S2B-C). Genes encoding ZAP-L regulated transcripts are enriched for plasma membrane, collogen-containing extracellular matrix, basement membrane and cell adhesion (Figure S2B), indicating that they might be associated with the plasma membrane or secreted. An example is the previously identified ZAP-regulated gene TNFRSF10D (also known as TRAILR4), which acts as a decoy receptor to inhibit TRAIL-mediated apoptosis^26,27^. Similar to HIV-CpG, transcript abundance from this gene is preferentially regulated by ZAP-L (Figure 2B). In contrast, genes encoding ZAP-S regulated transcripts are enriched for several transcription terms including transcription factor AP-1 complex (Figure S2C). Several genes encoding components of the AP-1 transcription factor were upregulated specifically in ZAP-S knockout cells, including JUN, JUNB, JUND and FOS (Figure 2C and S2D-F), and AP-1 activity was increased in ZAP-S knockout cells (Figure 2D).

ZAP-L and ZAP-S have differential subcellular localization because ZAP-L has a C-terminal CaaX box mediating S-farnesylation and membrane targeting^26,27,51–53^. ZAP-S is diffuse in the cytoplasm while ZAP-L localizes to the endoplasmic reticulum, endosomes, mitochondria and plasma membrane as well as the cytosol^26,27,51–53^. We determined that HIV-CpG infection does not alter ZAP-L or ZAP-S localization, with both showing the expected localization in uninfected and infected cells (Figure S2G). Because ZAP-L partially localizes to the ER, we determined the proportion of genes encoding transcripts regulated by ZAP-L or ZAP-S that contain a signal peptide sequence. 26% of ZAP-L regulated genes and 13% of ZAP-S regulated genes are predicted to encode a signal peptide (Figure 2E).

ZAP has been reported to regulate cellular transcript abundance through multiple direct or indirect mechanisms including inhibiting microRNA-mediated repression after stress or viral infection^68,69^, modulating stress responses^27^, modulating the type I interferon response^29,31,52^ and directly binding ZREs^26^. Therefore, we used the iCLIP data (Table S2) to determine which genes regulated by ZAP may encode transcripts that are direct targets. Transcripts from 16 of the 156 ZAP-L regulated genes and 42 of the 240 ZAP-S regulated genes had defined ZAP binding peaks. Because ZAP-L is more antiviral than ZAP-S, we focused on ZAP-L peaks. The average peak width for ZAP-L-regulated genes was 246 nt (Table S2), which is long enough for multiple ZAP molecules to bind. Nine of the 16 ZAP-L regulated genes with ZAP binding peaks had a predicted signal peptide or known transmembrane domain, again showing that ZAP-L regulates transcripts that localize to the ER and to other compartments. ZAP-L preferentially bound TNFRSF10D RNA (Figure 2F) and there were two defined peaks in the transcript (Figure S3). Interestingly, while UpAs are selected against in cellular transcripts expressed in the cytoplasm^70^, we observed abundant UpAs interspersed with CpGs in these peaks (Figure S3) and a UACG motif in each peak that matches the core of several ZAP-L binding motifs (Figure 2A and S2A).

### HIV-1 infection induces assembly of the ZAP-TRIM25-KHNYN complex and regulation of cellular transcripts

To determine if the presence of a substantial amount of ZAP substrate during HIV-CpG infection affects the interaction between ZAP, TRIM25 and KHNYN, we performed co-immunoprecipitation experiments in mock infected, HIV-WT and HIV-CpG infected HeLa cells (Figure 3A-B). Interestingly, upon HIV-WT or HIV-CpG infection, there was an increase in the RNase-resistant interaction between TRIM25 and KHNYN (Figure 3B and S4A), even though the abundance of these proteins was unchanged. As expected^41^, TRIM25 knockout did not affect the interaction between ZAP and KHNYN (Figure 3C). Therefore, viral infection appears to promote assembly of the ZAP-TRIM25-KHNYN complex, but this does not require the strong ZRE present in HIV-CpG.

**Figure 3:**
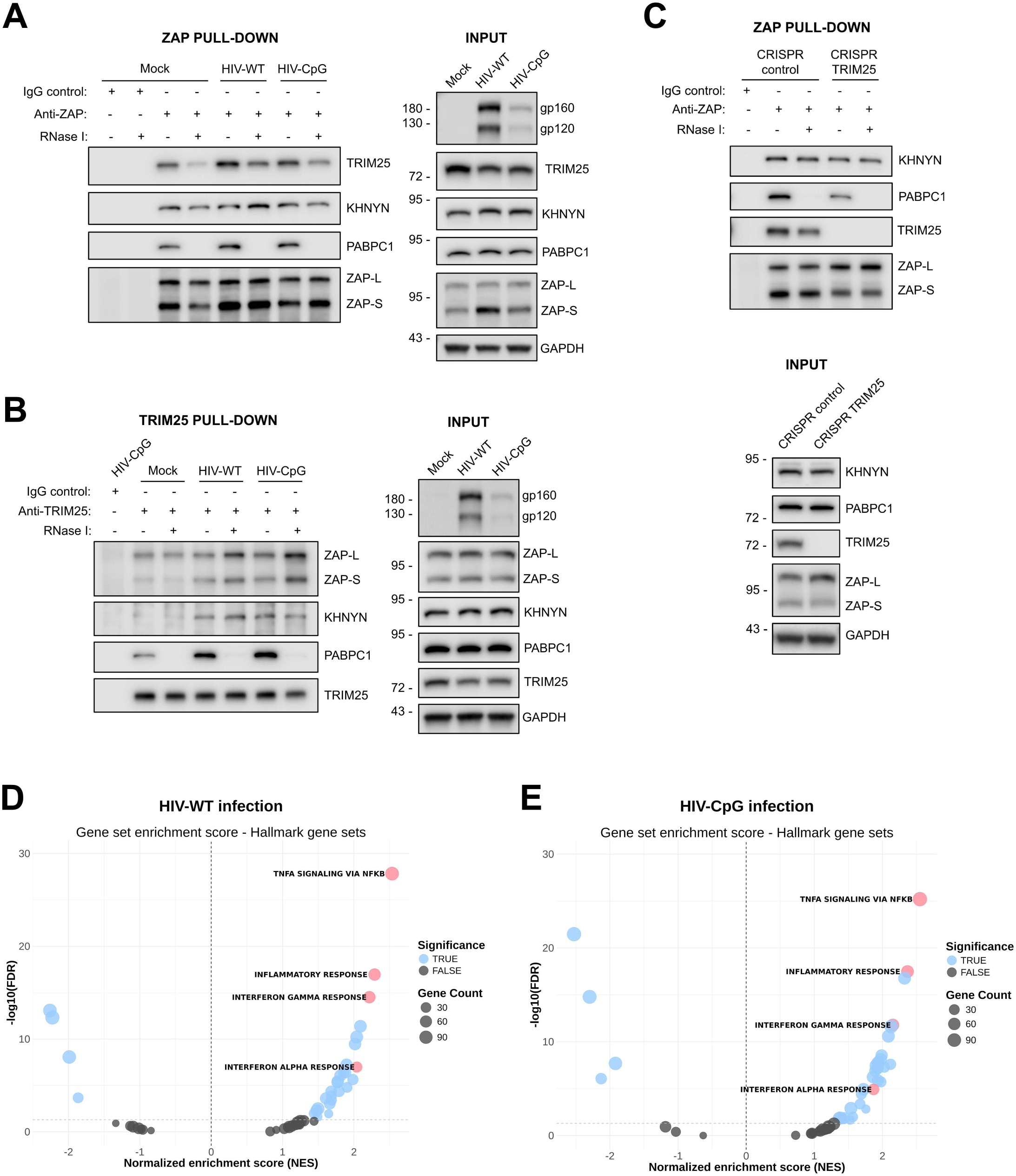
HIV-1 infection promotes the interaction between TRIM25 and KHNYN. **(A)** Immunoblotting of ZAP coimmunoprecipitations and corresponding input lysates obtained from infected HeLa cells. Samples were treated with RNase I after pulldown and PABPC1 is a control to indicate that the RNase I treatment reduced RNA-dependent interactions. The blots are representative of three biological replicates. **(B)** Immunoblotting of TRIM25 coimmunoprecipitations and corresponding input lysates obtained from infected HeLa cells. Samples were treated with RNase I after pulldown. Note that the experiment shown here was performed alongside the experiment shown in Figure 5E, thus the input blots for TRIM25, PABPC1, GAPDH, and gp160/gp120 and pulldown blots for TRIM25 and PABPC1 are identical between the two figures. The blots are representative of three biological replicates. **(C)** Immunoblotting of ZAP coimmunoprecipitations and corresponding input lysates obtained from infected Control and TRIM25 CRISPR HeLa cells. Samples were treated with RNase I after pulldown. Note that the experiment shown here was performed alongside the experiment shown in Figure S9I, thus the input blots for PABPC1, TRIM25, ZAP and GAPDH and pulldown blots for PABPC1, TRIM25 and ZAP are identical between the two panels. The blots are representative of three biological replicates. **(D-E)** Hallmark gene set enrichment plot of differentially expressed genes in HIV-WT **(D)** or HIV-CpG **(E)** infected HeLa CRISPR control cells. Black dots are non-significant, blue and pink dots are significant. Pink dots show functional groups highlighted in the text.

We then analyzed the effect of viral infection for ZMD of cellular transcripts using poly(A) RNA-seq data from mock, HIV-WT or HIV-CpG infected Control, ZAP, TRIM25 or KHNYN CRISPR cells (Table S3). Gene set enrichment analysis of differentially expressed genes between HIV-WT or HIV-CpG infected cells compared to mock infected cells showed a substantial change in transcripts associated with pathogen signaling including TNFα signaling, inflammatory response, interferon-α and interferon-γ responses (Figure 3D-E). This suggests that the increase in ZMD complex assembly correlates with an overall antiviral response and we posited that HIV-1 infection may stimulate ZMD activity.

To determine how HIV-WT and HIV-CpG infection affect transcripts regulated by ZMD, we focused on transcripts upregulated when ZAP, TRIM25 or KHNYN were knocked out. HIV-WT or HIV-CpG infection increased the number of genes expressing transcripts upregulated by ZAP or TRIM25 by ∼2-3-fold and KHNYN-upregulated transcripts by ∼5-fold (Figure S4B-D). Analysis of genes encoding transcripts co-upregulated by ZAP, TRIM25 and KHNYN showed an ∼10-fold increase in both HIV-WT and HIV-CpG infected cells compared to mock infected cells (Figure 4A-B). As expected from ZAP-L localization and preferential regulation of transcripts encoding membrane proteins^26,27^, the GO analysis for ZAP-TRIM25-KHNYN co-upregulated genes showed that these were enriched for encoding proteins associated with the endoplasmic reticulum (Figure 4C, Table S3). Because ZAP has been reported to regulate a subset of interferon-repressed genes^31^, we also analyzed whether ZAP-TRIM25-KHNYN were required for repression of some genes whose expression is downregulated after HIV-1 infection (Figure S5A). Exemplar genes that encode transcripts regulated by ZAP, TRIM25 and KHNYN include the previously reported ZAP-regulated gene ZMAT3^71^ as well as HSPB8 and DIO2, which is downregulated upon viral infection (Figure 4D-G and S5B-C).

**Figure 4:**
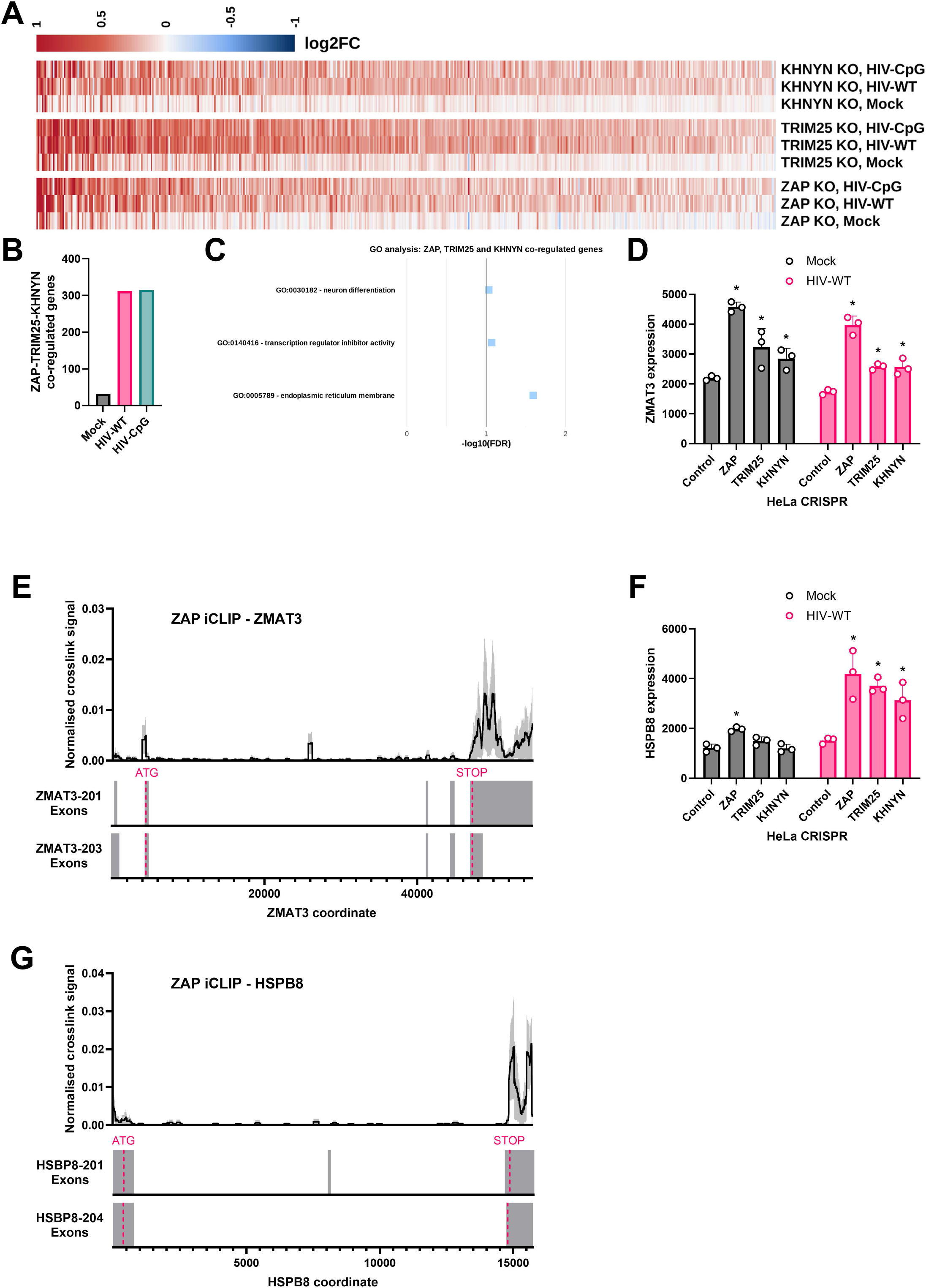
HIV-1 infection increases the number of cellular genes that are co-upregulated by ZAP, TRIM25, and KHNYN. **(A)** Heat map indicating differentially expressed genes that are co-upregulated in ZAP, TRIM25 and KHNYN CRISPR cells compared to CRISPR Control cells in mock, HIV-WT or HIV-CpG infected cells. **(B)** Number of genes co-upregulated by ZAP, TRIM25 and KHNYN in mock, HIV-WT and HIV-CpG infected cells. **(C)** GO analysis for ZAP, TRIM25 and KHNYN co-regulated genes with FDR <0.1 in HIV-WT infected cells. **(D)** Normalized DESeq2 counts for ZMAT3 from poly(A) RNA-seq in mock or HIV-WT infected Control, ZAP, TRIM25 or KHNYN CRISPR cells. Error bars represent standard deviation. *adjp <0.05 calculated by DESeq2, compared to Control CRISPR cells. **(E)** Normalized ZAP iCLIP signal mapped to ZMAT3 in HIV-WT infected Control CRISPR cells. The average signal of three independent experiments and the SEM (shading) are plotted. Exon tracks represent the isoforms expressed >1% of total normalized DESeq2 counts. The start and stop codon locations are indicated in pink. **(F)** Normalized DESeq2 counts for HSPB8 from poly(A) RNA-seq in mock or HIV-WT infected Control, ZAP, TRIM25 or KHNYN CRISPR cells. Error bars represent standard deviation. *adjp <0.05 calculated by DESeq2, compared to Control CRISPR cells. **(G)** Normalized ZAP iCLIP signal mapped to HSPB8 in HIV-WT infected Control CRISPR cells. The average signal of three independent experiments and the SEM (shading) are plotted. Exon tracks represent the isoforms expressed >1% of total normalized DESeq2 counts. The start and stop codon locations are indicated in pink.

ZAP, TRIM25 and KHNYN may also autoregulate ZAP-L abundance. ZAP-L expression has previously been shown to increase in TRIM25 knockout cells after human cytomegalovirus infection^72,73^, though this was initially proposed to be due to changes in alternative splicing of ZAP mRNA^72^. However, ZAP has recently been shown to bind a region in the 3’ UTR of ZAP-L^29^, which may indicate that ZAP-L abundance is controlled by TRIM25 and KHNYN through this ZRE. We confirmed that there is a strong ZAP binding peak encompassing the ZAP-L mRNA 3’ UTR that is not present in the ZAP-S mRNA (Figure S5D-E). We also observed that ZAP-L expression increased in TRIM25 and KHNYN knockout cells (Figure S5F-G). An isoform level analysis of poly(A) RNA-seq showed that this correlates with increased ZAP-L mRNA abundance in HIV-infected TRIM25 and KHNYN knockout cells (Figure S5H) This highlights that ZAP, TRIM25 and KHNYN autoregulate ZAP-L expression. Overall, HIV-1 infection increases the interaction between TRIM25 and KHNYN and this correlates with an increase in the number of genes that are co-regulated by ZAP, TRIM25 and KHNYN, implying that an infectious stimulus modulates the activity of ZMD on host transcripts.

Previous reports have shown that TRIM25 depletion decreases the interaction between ZAP and Ebola virus RNA or a reporter transcript that is regulated by ZAP at the level of translation^39,74^. Consistent with these reports, RNA-immunoprecipitation experiments showed that ZAP binding to HIV-CpG RNA was moderately decreased in TRIM25 knockout cells (Figure S6A). Analyzing ZAP iCLIP data for HIV-CpG infected TRIM25 knockout cells also showed a moderate decrease in ZAP bound to viral RNA (Figure S6B-C). TRIM25 could directly regulate ZAP binding to viral RNA or depleting TRIM25 could sequester ZAP on cellular transcripts that are unable to undergo decay, which would limit the pool of ZAP available to bind exogenous transcripts such as viral RNA. We analyzed the effect of TRIM25 depletion on ZAP binding to cellular transcripts in our iCLIP datasets and found that 127 genes had increased normalized read abundance in TRIM25 CRISPR cells compared to Control CRISPR cells (Figure S6D). Most of the 52 genes downregulated for ZAP binding had low read abundance. ZAP (ZC3HAV1) was one of the top genes upregulated for ZAP binding in TRIM25 knockout cells and a substantial increase in signal was observed in the ZAP binding peak within the ZAP-L 3’ UTR (Figure S6E), which could reflect both the increased ZAP-L abundance in TRIM25 depleted cells and ZAP accumulation on the transcript due to reduced decay. Taken together, the decreased ZAP binding we and others have observed on exogenous transcripts when TRIM25 is depleted could be due to increased ZAP binding on cellular transcripts and therefore a smaller pool of ZAP available to target viral RNA.

While TRIM25 has been proposed to regulate the specificity of ZMD in addition to its potency^41^, we did not observe any substantial change in ZAP antiviral activity against HIV-WT in TRIM25 knockout cells (Figure S5G). Furthermore, the relative position of ZAP binding to HIV-CpG RNA did not change in TRIM25 knockout cells (Figure S6B, Spearman correlation of relative crosslink signal at each nucleotide in CRISPR control and TRIM25 knockout cells = 0.92). PEKA analysis of iCLIP data for cellular transcripts in CRISPR control and TRIM25 knockout cells showed that the UpA + CpG motifs identified for ZAP-L were enriched in both conditions and there was no substantial change in the number of cellular transcripts with ZAP binding peaks (Figure S6F-G, Table S4). Therefore, TRIM25 appears to moderately regulate ZAP binding to viral RNA, possibly due to ZAP being sequestered on cellular RNA when TRIM25 is depleted, but does not change the ZAP targeting specificity.

### XRN1 degrades the 3’ fragment after endonucleolytic cleavage

We then sought to test the exonucleolytic and endonucleolytic decay models for ZMD (Figure 1A-B). One prediction of the ZMD endonucleolytic model (Figure 1B) is that KHNYN binding would be present in regions with clustered CpGs. Therefore, we used iCLIP to map KHNYN binding sites on HIV-CpG RNA. Because we have not found an antibody that efficiently immunoprecipitates KHNYN, we identified KHNYN binding sites on viral RNA using a previously characterized stable cell line expressing an active KHNYN-GFP fusion protein^61^. KHNYN had substantial binding within areas of high CpG abundance, including the CpG recoded region of HIV-CpG (Figure 5A). It also bound motifs containing a CpG (Figure S7A). As KHNYN has no known RNA binding specificity^42,44^, this supports the hypothesis that ZAP recruits KHNYN to target RNA. Interestingly, motifs enriched around KHNYN crosslink sites also contain UpA dinucleotides, including UACG motifs (Figure S7A).

**Figure 5:**
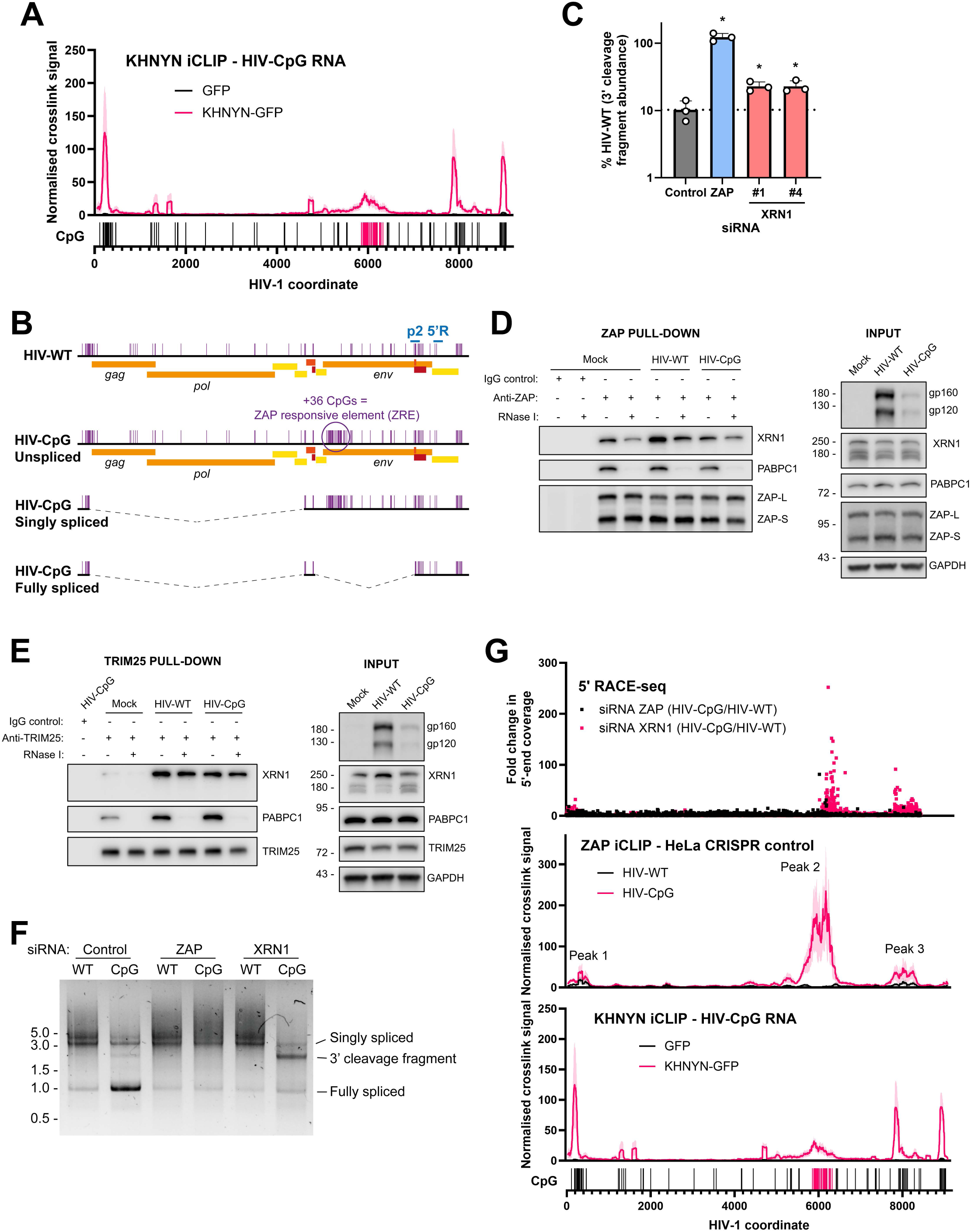
Degradation of HIV-CpG 3’ cleavage fragment by XRN1. **(A)** KHNYN iCLIP signal mapped to HIV-CpG gRNA in HeLa CRISPR KHNYN + GFP and HeLa CRISPR KHNYN + CRISPR-resistant KHNYN-GFP. The average signal of three independent experiments and the SEM (shading) are plotted. **(B)** Genomic organization of HIV-WT and HIV-CpG and localization of CpG dinucleotides. The qRT-PCR probe used to quantify the 3’ cleavage fragment (probe 2 (p2)), and the HIV specific primer used for the 5’ RACE (5’R) are shown. **(C)** Level of HIV-CpG 3’ cleavage fragment measured by qRT-PCR in cells transfected with siRNA targeting XRN1. Results are expressed as % 3’ cleavage fragment in HIV-WT infected cells. The dotted line represents the level of 3’ cleavage fragment in cells treated with a non-targeting siRNA control. Error bars represent standard deviation. *p <0.05 calculated by one-way ANOVA on log-transformed data, compared to the level of 3’ cleavage fragment in cells treated with a non-targeting siRNA control. **(D)** Immunoblotting of ZAP coimmunoprecipitations and corresponding input lysates obtained from infected HeLa cells. Samples were treated with RNase I after pulldown. The blots are representative of three biological replicates. **(E)** Immunoblotting of TRIM25 coimmunoprecipitations and corresponding input lysates obtained from infected HeLa cells. Samples were treated with RNase I after pulldown. Note that the experiment shown here was performed alongside the experiment shown in Figure 3B, thus the input blots for gp160/gp120, PABPC1, TRIM25, and GAPDH and the pulldown blots for PABPC1 and TRIM25 are identical between the two figures. The blots are representative of three biological replicates. **(F)** Agarose gel electrophoresis of 5’ RACE reactions. Sizes in kilobases are indicated on the left. **(G)** 5’ RACE DNA was sequenced by long-read sequencing and the 5’ end of each molecule was identified and mapped onto HIV gRNA. The number of 5’ end at every position was determined and compared between experimental conditions. iCLIP signals for ZAP (from Figure S6B) and KHNYN (from panel A) are shown for reference.

For the ZMD exonucleolytic decay model (Figure 1A), ZAP has been reported to interact with PARN, the exosome complex (EXOSC5, EXOSC7), DCP2, XRN1, DDX17 and DDX30^33–35,37,75^. In addition, ZAP has been reported to interact with components of the CCR4-NOT deadenylation complex^76,77^. The endonucleolytic and exonucleolytic models could be reconciled if these are parallel pathways or if some of the exonucleolytic model components are required for RNA decay after endonucleolytic cleavage. To identify whether any of these proteins are required for ZAP antiviral activity on HIV-CpG, we depleted them using RNAi. While siRNA-mediated depletion of ZAP, TRIM25 and KHNYN increased HIV-CpG infectious virus production, none of the ZAP cofactors implicated in the exonucleolytic model had an effect (Figure S7B-E), consistent with a previous study^9^. This suggests that the endonucleolytic pathway is dominant for restricting HIV-CpG, leading to a cleaved viral genome that can’t be translated or packaged into virions. However, exonucleases could be required to degrade viral RNA fragments after KHNYN cleavage. Therefore, we designed a probe to potentially identify the 3’ cleavage fragment of HIV-1 RNA with the presumption that many KHNYN cleavage sites will be near the major ZAP binding site in the CpG-recoded region (probe 2 (p2) in Figure 5B). Of note, due to the complex HIV-1 alternative splicing program^64^, only the unspliced and singly spliced HIV-1 transcripts will contain the ZAP-sensitive CpG-recoded region and the fully spliced RNAs are not subject to ZMD^9,40^ (Figure 5B). To quantify only ZAP-sensitive transcripts, p2 was designed to span SA7, which is the last sequence that is specific to the singly and unspliced transcripts. Notably, this sequence is within ZAP binding peak 3 (Figure 1I). RNAi-mediated depletion of CNOT1, PARN, EXOSC3, EXOSC5, EXOSC7, DDX17 or DHX30 did not affect HIV-CpG RNA abundance using the probe for the 3’ fragment (Figure S7F-G), but depletion of the 5’-3’ exonuclease XRN1 led to a significant increase in abundance (Figure 5C).

To determine if ZAP and TRIM25 interact with XRN1, we performed co-immunoprecipitation experiments in the absence and presence of HIV-1 infection (Figure 5D-E and Figure S7H). Both proteins interact with XRN1 in a RNase-resistant manner and the interaction between TRIM25 and XRN1 is stimulated by either HIV-WT or HIV-CpG infection. Similar to the interaction between ZAP and KHNYN, TRIM25 is not required for ZAP to interact with XRN1 (Figure S7I). Overall, we hypothesize that a ZAP-TRIM25 complex recruits XRN1 to degrade the 3’ cleavage product.

An important prediction of the endonucleolytic model for ZMD is that internal cleavage sites should be identified when degradation of the cleavage products is inhibited. Therefore, we used 5’ RACE to identify potential cleavage sites in XRN1 depleted cells using a HIV-1 specific primer at the 3’ end of the viral genome (Figure 5B). Visualizing the RACE products on an agarose gel showed that a novel band appeared for the control siRNA HIV-CpG sample (3’ cleavage fragment in Figure 5F). When ZAP was depleted, this band was not present and the abundance of the band increased when XRN1 was knocked down. Therefore, we hypothesized that this band could be a KHNYN cleavage product that is further degraded by XRN1. To map the cleavage sites, we used Oxford Nanopore sequencing for the total 5’ RACE reactions and identified the 5’ end of each read. In XRN1 knockdown cells, HIV-CpG 5’ ends clustered in ZAP binding Peaks 2 and 3 (Figure 5G). This indicates that ZAP likely recruits KHNYN to these sites to mediate cleavage and that the ZAP binding in Peak 3 is sufficient for KHNYN cleavage. This also suggests that the increase in 3’ cleavage fragment abundance when XRN1 is depleted (Figure 5C) may be underestimated because probe 2 (Figure 5B) covers the site of endonucleolytic cleavage in ZAP binding Peak 3. However, as described above, this is the most distal 3’ sequence that the probe can cover and still be specific for the ZRE-containing RNA. Importantly, the 5’ RACE assay has a 3’ bias in that if KHNYN cuts multiple times in a transcript, the 5’ end closest to the RACE primer will be identified, so the relative frequency of cleavage sites in Peaks 2 and 3 should not be directly compared. The sequence underlying Peak 3 is identical between HIV-WT and HIV-CpG, suggesting that ZAP binding in Peak 2 not only promotes ZAP binding in Peak 3 (Figure S1D and H) but that this binding is functional, leading to endonucleolytic cleavage.

We identified 153 high confidence cleavage sites spanning nt 6007-8299. Interestingly, 68 of these were within a 92 nt region from nt 6261-6352 in which almost every nucleotide was a high confidence cleavage site (Figure S8A-B). Within this region, both CpG and UpA are enriched (observed/expected CpG = 1.72, UpA = 1.48). There are nine UACG motifs in the HIV-CpG genome, seven of which have been introduced by CpG-recoding, and three of these are found within the 92 nt region (Figure S8B). There was strong ZAP iCLIP signal and moderate KHNYN iCLIP signal (Figure S8A), which may be indicative of active cleavage and a short KHNYN residence time on the RNA. Therefore, frequent endonucleolytic cleavage correlated with high levels of ZAP binding in a region containing UACG motifs, consistent with the sequence within several of the top ZAP-L iCLIP motifs (Figure 2A and S2A).

### The 5’ cleavage fragment is 3’ uridylated by TUT4/7 and then degraded by DIS3L2

To determine how the 5’ cleavage fragment was degraded, we set up a qRT-PCR assay with a primer-probe set upstream of Peak 2 (probe 1 (p1) in Figure S9A) and then screened the RNA degradation proteins linked to ZAP to determine if they regulate the abundance of this fragment. Surprisingly, depletion of three exosome subunits, EXOSC3, EXOSC5 or EXOSC7, had no effect on the abundance of the 5’ cleavage fragment (Figure S9B). Depletion of CNOT1, PARN, XRN1, DDX17 or DHX30 also had no effect, indicating that degradation of the 5’ fragment was not regulated by known ZAP-interacting proteins linked to RNA decay. Therefore, we performed 3’ RACE to identify the location and identity of the 3’ ends of the 5’ cleavage fragment (Figure 6A and S9A). A potential 5’ cleavage fragment was observed specifically in cells infected with HIV-CpG and was not present when ZAP was depleted. We cloned this band and sequenced seven clones using Sanger sequencing. Interestingly, reads terminated within ZAP binding Peak 2 and several had non-templated uridines at the 3’ end (Figure 6B). This result raised the possibility that the 5’ fragment was modified by 3’ uridylation.

**Figure 6:**
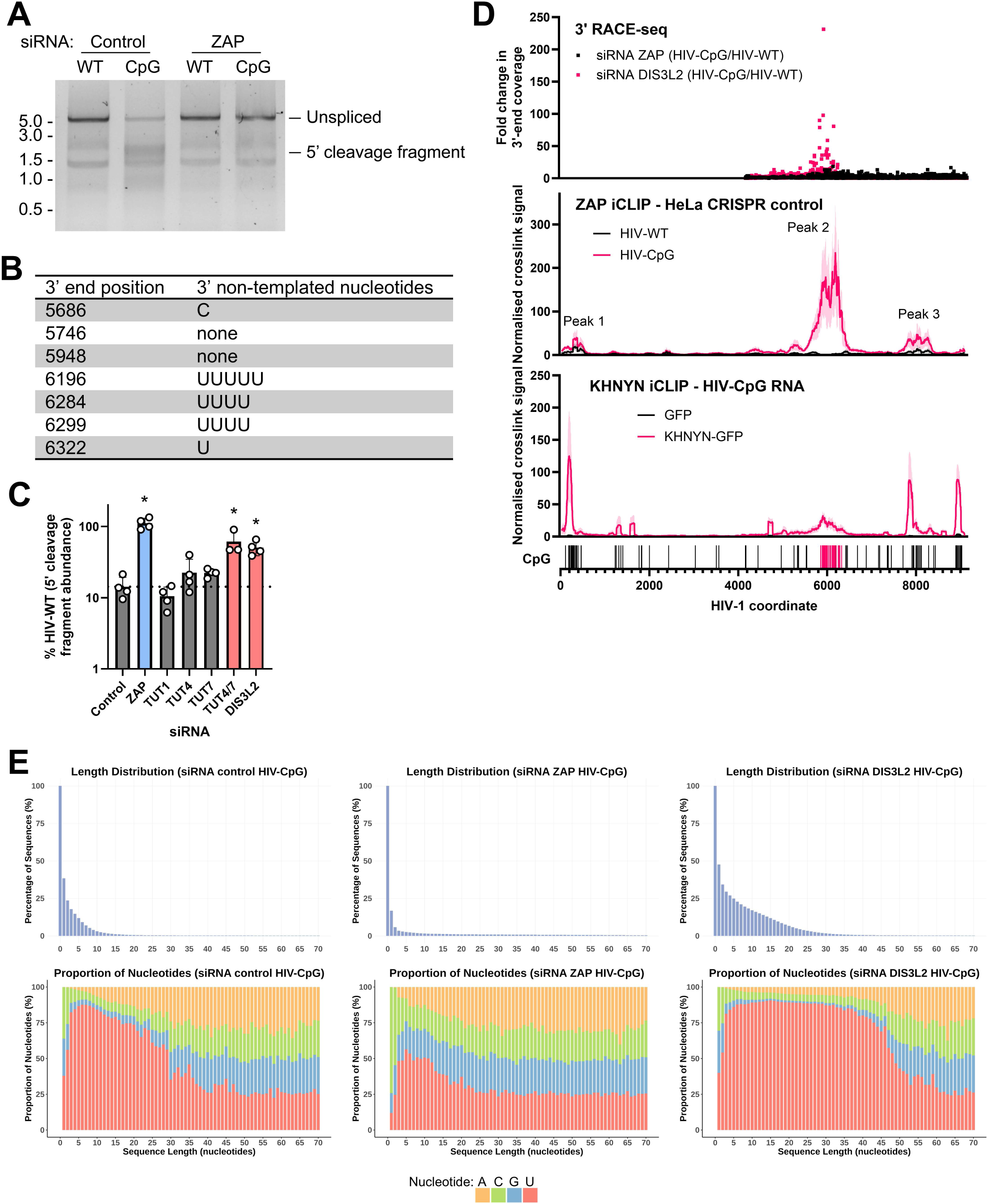
HIV-CpG 5’ cleavage fragment abundance is regulated by TUT4/TUT7 and DIS3L2. **(A)** Agarose gel electrophoresis of 3’ RACE reactions. Sizes in kilobases are indicated on the left. **(B)** Non-templated nucleotides are found in the 3’ end of HIV-CpG 5’ cleavage fragments amplified by 3’ RACE and determined by Sanger sequencing. The 3’ end positions shown here all fall within the CpG-enriched region of HIV-CpG. **(C)** Level of HIV-CpG 5’ cleavage fragment measured by qRT-PCR in cells transfected with siRNA targeting TUTs and DIS3L2. Results are expressed as % 5’ cleavage fragment in HIV-WT infected cells. The dotted line represents the level of 5’ cleavage fragment in cells treated with a non-targeting siRNA control. Error bars represent standard deviation. *p <0.05 calculated by one-way ANOVA on log-transformed data, compared to the level of 5’ cleavage fragment in cells treated with a non-targeting siRNA control. **(D)** 3’ RACE DNA was sequenced by long-read sequencing and the 3’ end of each molecule was identified and mapped onto HIV gRNA. The number of 3’ end at every position was determined and compared between experimental conditions. iCLIP signals for ZAP (from Figure S6B) and KHNYN (from Figure 5A) are shown for reference. **(E)** The sequence and length of non-templated nucleotides was extracted for every sequenced 3’ end in HIV-CpG infected cells transfected with siRNA control or siRNA targeting ZAP or DIS3L2.

Several members of the TENT family of polymerases add non-templated nucleotides at the 3’ end of transcripts and TUT1, TUT4 and TUT7 are three proteins that specifically add non-templated uridines^78^. TUT1 is a nuclear protein that had no effect when it was depleted (Figure 6C). In contrast, depletion of two cytoplasmic paralogs that are often functionally redundant, TUT4 and TUT7, promoted an increase in the abundance of the 5’ cleavage fragment (Figure 6C and S9C). As expected, TUT4/7 depletion had no effect on infectious virus production, indicating that it acts downstream of endonucleolytic cleavage (Figure S9D), or the abundance of the 3’ cleavage fragment (Figure S9E). We then analyzed whether ZAP and TRIM25 interacted with TUT7. Both proteins interacted with it in an RNase-resistant manner and this interaction was increased in the presence of HIV-WT or HIV-CpG infection (Figure S9F-H). The interaction between ZAP and TUT7 was not altered in TRIM25 knockout cells (Figure S9I).

DIS3L2 is an exosome-independent, 3’-5’ exoribonuclease that degrades transcripts with 3’ terminal uridylation^79–81^. Depleting its expression increased expression of transcripts using the 5’ probe but not the 3’ probe (Figure 6C and S10A-B), indicating that it is required to degrade the 5’ KHNYN cleavage fragment. DIS3L2 depletion does not affect HIV-CpG infectious virus production (Figure S10C), indicating that it also acts after endonucleolytic cleavage. ZAP and TRIM25 interact with DIS3L2 in an RNase-resistant manner, and the interaction between TRIM25 and DIS3L2 is increased by HIV infection (Figure S10D-F). Knocking out TRIM25 did not affect the interaction between ZAP and DIS3L2 (Figure S10G). We then used 3’ RACE-seq to map internal cleavage sites in HIV-CpG after DIS3L2 depletion and found that they are concentrated in ZAP binding Peak 2 (Figure 6D). We also determined proportion of 3’ uridylation and length. When DIS3L2 is depleted, ∼50% of the reads have at least one non-templated nucleotide and a substantial portion have at least 10 non-templated nucleotides (Figure 6E). These non-templated nucleotides are highly enriched in uridine, indicating a poly-uridine tail. In contrast, when ZAP is depleted, few reads contain non-templated tails and the strong enrichment for uridine is not present. These data indicate that after KHNYN cleavage, the 5’ cleavage products of the viral RNA become 3’-polyuridylated by TUT4/7 and are then degraded by DIS3L2.

Overall, these experiments lead to a comprehensive model for ZMD (Figure 7). ZAP-L forms a complex with TRIM25 and KHNYN to direct KHNYN-mediated endonucleolytic cleavage in regions of high ZAP binding. After cleavage, ZAP and TRIM25 interact with additional RNA decay machinery to mediate degradation of the resultant fragments. XRN1 degrades the 3’ cleavage fragment. The 5’ cleavage fragment is 3’ uridylated by TUT4/7 and then degraded by DIS3L2. This allows the entire viral RNA to be degraded after its recognition by ZAP.

**Figure 7:**
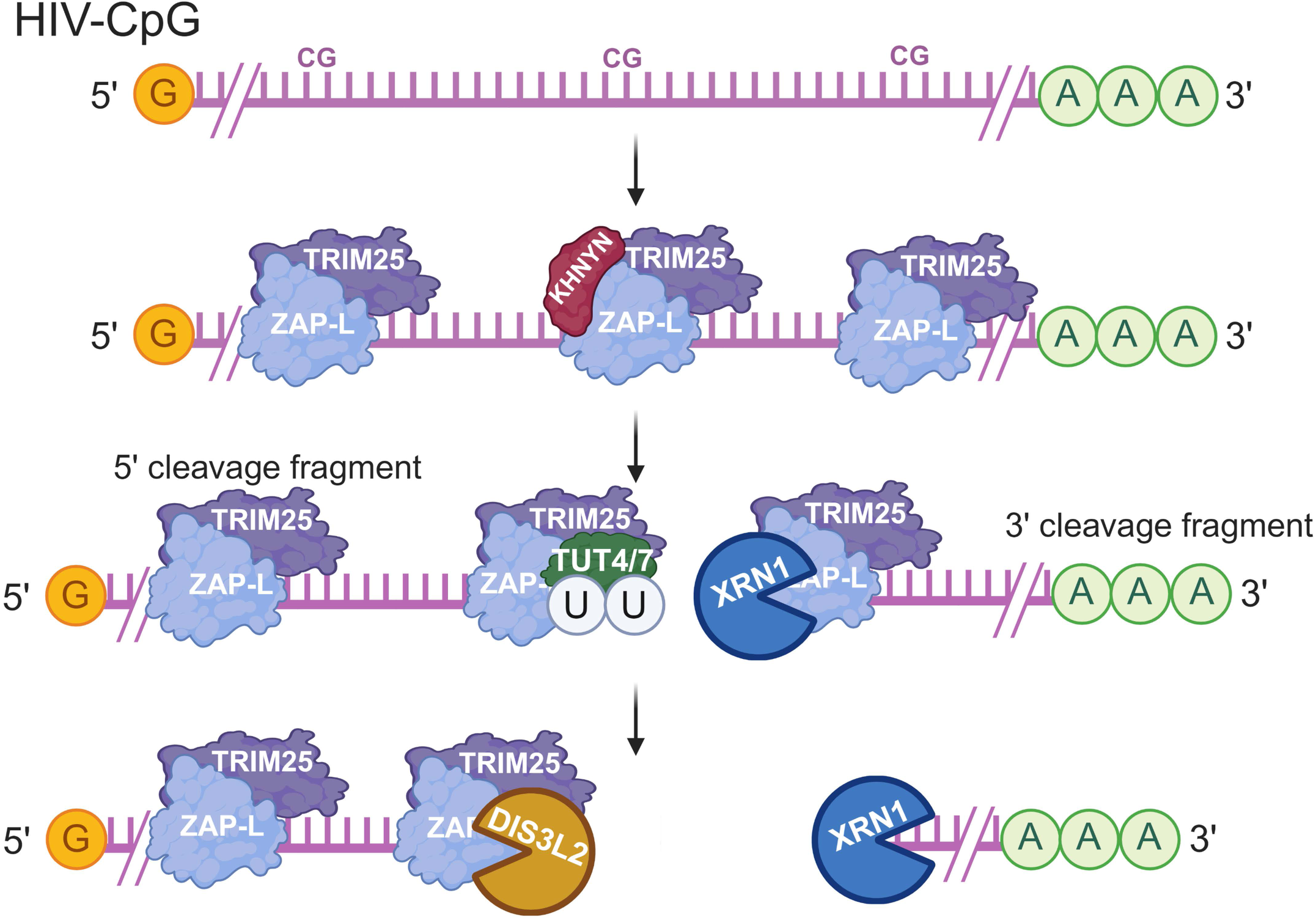
Model of the ZAP-mediated RNA decay. A ZAP-L-TRIM25-KHNYN complex assembles on clustered CpG dinucleotides. This leads to RNA cleavage by KHNYN. The 3’ cleavage fragment is then degraded by the 5’-3’ exonuclease XRN1, while the 5’ cleavage fragment is uridylated by TUT4/7 and then degraded by the 3’-5’ exonuclease DIS3L2. ZAP interacts with each component of the complex. Created in BioRender. Bouton, C. (2025) https://BioRender.com/f8zm1lm

## DISCUSSION

Determining how ZAP selects specific transcripts for decay is essential to utilize it for novel vaccine and oncolytic virus development and to prevent it from targeting therapeutic RNAs. A simple model for ZMD specificity is that ZAP binds CpG dinucleotides through ZnF2 in its RBD and regions containing clustered CpGs are sufficient to mediate specificity^9,10,12,17^. Supporting this model, ZAP has high affinity for RNAs containing a CpG dinucleotide and CpG recoding can sensitize diverse viruses to ZAP-mediated restriction^9–14^. However, some regions of viral RNA appear to be more sensitive to CpG-mediated ZAP restriction and high levels of CpG abundance sometimes does not sensitize viral RNA to ZAP^14,21–25^. In particular, this has been shown for synonymous recoding of influenza virus to develop novel live attenuated virus vaccines.

Some CpG recoding strategies have led to ZAP-mediated attenuation and in other cases substantial increases in CpG abundance have not sensitized the virus to ZAP^14,22,82,83^. In addition, UpA recoding has been reported to sensitize viral RNA to ZAP^11,13,16^. Our data indicate that while both ZAP-L and ZAP-S bind motifs containing a CpG, ZAP-L binds motifs that also contain an upstream UpA. This may explain why UpA recoding can sensitize a virus to ZAP-mediated restriction. Many viral genomes are depleted in both CpG and UpA dinucleotides^70^ and introducing both into such viruses may increase their sensitivity to ZAP-L. Supporting this hypothesis, a study published while this manuscript was under revision compared CpG to CpG + UpA recoding for West Nile virus and found that CpG + UpA recoding had superior pre-clinical vaccine characteristics^15^.

Why ZAP-L and ZAP-S bind different RNA sequence motifs given they have the same RBD remains unclear, but it is likely directly or indirectly regulated by the ART-like domain and C-terminal S-farnesylation. Adding a CaaX box at the C-terminus of ZAP-S partially increases its antiviral activity, and this correlates with its localization to the endomembrane system^52,53^. However, transferring the CaaX box to the ZAP-S does not restore full antiviral activity and the ART-like domain likely has a role as well, though its function is not known. While ZAP-S is predominately cytosolic, ZAP-L localizes to multiple organelles in the endomembrane system as well as the cytosol^26,27,51–53^ and may target transcripts in several cytoplasmic regions. Recent studies have highlighted that cellular transcripts associate with multiple compartments in the endomembrane system including the ER, outer mitochondrial membrane, endosomes and TIS granules^84–86^. High resolution studies will be required to determine how ZAP-L and ZAP-S subcellular localization controls transcripts that localize in specific cytoplasmic compartments.

ZAP-L and ZAP-S could bind different motifs through interacting with different cofactors or due to conformational changes, possibly regulated by membrane binding. The ZAP RBD has four zinc fingers and RNA binding has been observed in regions outside of the ZnF2 CpG binding pocket^12,20,87^. In addition, it has been reported to interact with many other RNA binding proteins^4^. How ZnF1, ZnF3 and ZnF4 in the ZAP RBD and other ZAP-interacting RNA binding proteins contribute to specific RNA targeting and ZMD activity will require future studies.

While we focused on ZAP-L-mediated RNA decay in this study, ZAP-S has several reported roles and how RNA binding differences between ZAP-L and ZAP-S regulate different aspects of cellular function remain an important area for further investigation. ZAP-S has been shown to regulate RIG-I signalling^47^, interferon mRNA abundance^52^ and the unfolded protein response during cellular stress^27^ as well as restrict SARS-CoV-2 by inhibiting programmed ribosomal frameshifting^8^. An interesting implication for how ZAP-L and ZAP-S regulate different transcripts is how this contributes to diseases in which ZAP expression is dysregulated, such as some types of cancer^26,55,88^. While several studies have analyzed ZAP-regulated transcripts in the context of knocking out both ZAP-S and ZAP-L^26,28,29^, here we have extended a previous study^27^ showing differential transcript targeting by the two isoforms. ZAP-L regulation of TNFRSF10D and ZAP-S regulation of AP-1 activity are two examples of how ZAP isoforms may impact tumorigenesis. While the functional consequences of alterations in ZAP-L or ZAP-S expression for cellular transcripts requires further study, future characterization of the role of ZAP in tumors should consider isoform specific changes as well as overall changes in ZAP expression.

In this study, we have focused on the role of KHNYN for mediating the endonucleolytic cleavage event but N4BP1 may also cleave ZAP-targeted RNA. N4BP1 is a core vertebrate ISG^89^ and is functionally redundant with KHNYN for some transcripts^41,43,44^. N4BP1 targets specific mRNAs^90^ and whether N4BP1 and KHNYN have differential transcript specificity is not clear. While the ZAP RBD binds the KHNYN CUE-like domain^41^, several of the residues in KHNYN that mediate this interaction are not conserved in the N4BP1 CUE-like domain^61^. This implies that there may be further interactions between ZAP and KHNYN/N4BP1 or that they interact with other components of the ZAP antiviral system^54^. How KHNYN is regulated so that it only cleaves at sites directed by ZAP is unclear, though it shows specificity for targeting ZAP-sensitive viruses even when overexpressed^21,40^. KHNYN abundance on a per molecule basis is substantially lower than ZAP^61^ and it is likely a limiting factor for antiviral activity^40^. Structural studies will be required to understand how multiple ZAP molecules assemble on a ZRE and interact with KHNYN to provide targeted decay. Other endonucleases in the same gene family as KHNYN, such as ZC3H12A^91^, are controlled through a combination of post-translational modifications that regulate their stability and interaction with other proteins and this may be important for KHNYN activity as well. Furthermore, KHNYN is highly active in the presence of manganese^92^, which can be released into the cytosol after viral infection^93^, and local manganese abundance may also regulate its activity.

One of the major unresolved questions for ZMD is how TRIM25 regulates ZAP activity. TRIM25 has ubiquitylation/ISGylation activity^38,39,74,94^, multimerizes through its N-terminal RING domain^41,95^ and also binds RNA^96–98^. All of these functions have been proposed to regulate its antiviral activity and an interaction between TRIM25 and TUT4 has also been reported to control let-7 microRNA decay^99^. Consistent with previous reports^39,74^, we found that TRIM25 depletion decreased ZAP binding to viral RNA. However, this effect is moderate and may be due to an increased occupancy of ZAP on cellular transcripts, which could decrease the pool of ZAP available to target exogenous RNA. Because TRIM25 binds both cellular and viral RNA^96–102^, it could act as a co-reader with ZAP to ensure target specificity^29^. Alternatively, it could promote multimeric ZAP complex formation through RING domain dimerisation^41^ or modify a component of the ZAP antiviral system with ubiquitin or ISG15^38,39,74,94^. Specific mutations in TRIM25 that isolate different activities will be required to determine how it regulates ZAP antiviral activity, though this is challenging since TRIM25 binds ssRNA and dsRNA through multiple sites and RNA binding, multimerization and E3 ubiquitin ligase activity are linked^96–98^. HIV infection increased TRIM25 interaction with KHNYN, XRN1, TUT7 and DIS3L2 and this correlates with increased repression of cellular transcripts by ZAP, TRIM25 and KHNYN. Importantly, this occurred for both HIV-WT and HIV-CpG infection, indicating that viral RNA substrate is not required for increased TRIM25 interaction with RNA decay enzymes. While TRIM25 RNA binding activity is regulated by pH^54^ and local pH changes caused by viral infection could modulate its interaction with RNA, possibly through virion fusion with endosomes or alterations in mitochondria, we note that the interaction between TRIM25 and the RNA decay enzymes was RNase-resistant. Therefore, additional changes in TRIM25 in the context of viral infection, such as post-translational modifications, may control the ZAP antiviral system by altering the TRIM25 interactome.

We have shown that depletion of the 5’ RNA decay fragment requires TUT4/7, indicating that uridylation after an internal cleavage event is required for complete viral RNA decay. While 3’ uridylation has previously been linked to restriction of viruses^103,104^ and LINE-1 retrotransposition^105^, this has predominately been proposed to occur at the 3’ end of the transcript. One reason why 3’ uridylation might be critical for degradation of viral RNA is that it can recruit DIS3L2, which can degrade structured RNA^106^. Higher order RNA structures are common in viral RNA, especially positive strand RNA virus genomes. For example, the HIV-1 genome has several regions of RNA structure^107^ and the ability of DIS3L2 to degrade structured RNA may be required to completely destroy the viral RNA. It should be noted that many ZAP-sensitive viruses produce viral RNA that comprise a substantial percentage of total poly(A) RNA^108–110^. Complete degradation of the RNA may be important to prevent partially degraded RNA species from sequestering cellular RNA binding proteins, stimulating the innate immune response or otherwise dysregulating cellular RNA metabolism.

## LIMITATIONS OF THE STUDY

While the iCLIP experiments demonstrate ZAP binding over three broad regions in HIV-CpG RNA, this technique does not determine the number and location of ZAP molecules bound to HIV-CpG at individual transcript resolution. Therefore, while we hypothesize that ZAP multivalent binding is important for target specificity, which is supported by previous genetic studies^9,17^, the iCLIP experiments do not show this definitively. Furthermore, while the RACE-seq experiments show that ZAP-specific internal cleavage sites correlate with regions of ZAP binding on HIV-CpG, we cannot correlate cleavage sites with specific ZAP binding sites on an individual transcript basis.

## METHODS

### Cell lines, viruses and infection

HEK293T CRISPR cells^27^, HeLa, HeLa CRISPR cells^40,61^ and TZM-bI cells^111–113^ were maintained in high-glucose DMEM supplemented with GlutaMAX (Thermo Fisher Scientific), 10% fetal bovine serum, 100 U/mL penicillin and 100 µg/mL streptomycin and grown with 5% CO2 at 37°C.

HIV-WT and HIV-CpG stocks were produced by co-transfecting pHIV-WT^114^ or pHIV-CpG^40^ with pVSV-G^115^ into HEK293T CRISPR ZAP cells, harvesting at 48h post-transfection and titering on TZM-bl cells as described below. The sequences of HIV-WT and HIV-CpG are available in the National Center for Biotechnology Information Nucleotide Database with Accession MN685337 and MN685350, respectively.

Unless otherwise stated, cells were infected at MOI=3. The medium was replaced 18 hours (h) post-infection. Cell lysates and supernatants were collected at 48 hours post-infection. Cell-free supernatants were produced by centrifugation at 300×*g* for five minutes.

### siRNA transfection and infection

siRNA were purchased from Horizon Discovery and transfected using Lipofectamine RNAiMAX (Invitrogen). siRNAs were reverse-transfected with 1 µL lipofectamine at 25 nM final concentration for 90,000 HeLa cells in a 24-well plate. 24 hours later, the medium was replaced, and cells were transfected again with siRNA at 25 nM final concentration with 1µL lipofectamine. 6-8 hours later, cells were infected with HIV-1 at MOI=3. The medium was replaced 18 hours post-infection. Supernatant and cell lysates were collected 48 hours post-infection as described above.

### TZM-bl infectivity assay

The TZM-bl cell line was used to quantify the amount of infectious virus^111–113^. 1×10^4^ cells per well were seeded in 96-well plates and infected by incubation with virus stocks or supernatants of infected cells. At 40-48 hours post-infection, cells were lysed and infectivity was measured by β-galactosidase expression using the Galacto-Star assay system as per the manufacturer instructions. β-galactosidase activity was quantified as relative light units per second using a PerkinElmer luminometer.

### Immunoblotting

Cell lysates for immunoblotting were prepared in Laemmli buffer. Virions were pelleted from culture supernatant by centrifugation at 20,000 x *g* for 2 hours at 4°C through a 20% sucrose cushion in phosphate-buffered saline (PBS). Viral pellets were resuspended in 2x Laemmli buffer. Samples were boiled for 10 min at 95°C, separated on 8-16% gradient precast gels (BioRad; Millipore) and then transferred to nitrocellulose or PVDF membranes (Amersham). Membranes were briefly rinsed in PBS then blocked in blocking buffer (PBS, 5% non-fat milk, 0.1% Tween 20) for 30 minutes at RT. Membranes were incubated with primary antibodies diluted in blocking buffer at 4°C overnight, then washed 3 times for 5 min in PBS-T (PBS, 0.1% Tween 20), then incubated with secondary antibodies diluted in blocking buffer for 1h at RT, then washed for 3 times for 5 min in PBS-T. Membranes were imaged using an Odyssey FC (LI-COR) measuring secondary antibody fluorescence, or using Amersham ECL Prime Western Blotting Detection reagent (Cytiva) for HRP-linked antibodies with an Amersham ImageQuant 800 (Cytiva). Antibodies used in this study are reported in the STAR methods table.

### Coimmunoprecipitation

1.5 mg Dynabeads protein A (Invitrogen) per sample were incubated in lysis buffer (0.5% Nonidet P 40 (NP40) substitute, 10mM HEPES pH=7.5, 150mM KCl, 3mM MgCL2, protease inhibitor) supplemented with 5% BSA for 30 min at RT and under gentle agitation. 4 µg of antibody per sample was bound to the beads for 30 min at RT and under gentle agitation. The unbound antibody was then washed off and the beads were resuspended in 50 µL lysis buffer per sample. Cells cultivated in a 10-cm dish were lysed on ice in 1mL lysis buffer. In case of infected cells (MOI=3), cells were lysed 48 hours post-infection. Lysates were gently mixed on a rotator for 30 min at 4°C, then centrifuged at 20,000 ×*g* for 10 min to pellet the nuclear fraction. 75 µL of the post-nuclear supernatant was saved as the input lysate and mixed with 1 volume of Laemmli buffer. 900 µL of the remaining post-nuclear supernatant was incubated with the prepared beads under gentle agitation over night at 4°C. The samples were then washed four times in wash buffer (0.05% Nonidet P 40 substitute, 10mM HEPES pH=7.5, 150mM KCl, 3mM MgCL2, protease inhibitor) before being resuspended in 90 µL Laemmli buffer. For experiments including an RNase I treatment, the samples were washed twice in wash buffer after the overnight incubation with the prepared beads, then resuspended in 100 µL wash buffer. The samples were incubated with 300 U RNase I for 30 min at 37°C and washed three times in wash buffer before being resuspended in 90 µL Laemmli buffer. Input lysates and samples were analyzed by immunoblotting as described above. Coimmunoprecipitation and pull-down immunoblots were quantified using Image Studio (LI-COR Biotech). Relative coimmunoprecipitation efficiency was calculated as the ratio of coimmunoprecipitation to pulldown then normalized to the same ratio calculated in control conditions.

### qRT-PCR

Total RNA was extracted using the RNeasy Mini Kit (Qiagen). cDNA was synthesized using the High-Capacity cDNA RT kit (Applied Biosystems) following the manufacturer’s instructions. cDNA was diluted 20 times in RNase-free water and quantified by qPCR using Applied Biosystems TaqMan Universal PCR Master Mix (Applied Biosystems) in a QuantStudio 5 qPCR System (Applied Biosystems). Relative cDNA quantities were determined using the ΔΔCt method. The primer-probe set for these experiments are in the 3’ end of *gag* and are reported in the STAR methods table.

### RNA Immunoprecipitation

HeLa CRISPR Control/TRIM25/KHNYN cells^40^ were plated in 6-well plates and transfected using TransIT-LT1 transfection reagent with 500 ng of HIV-WT or HIV-CpG proviral DNA, and medium was replaced 4-6 hours later. 48 hours post-transfection cells were washed with PBS prior to UVC crosslinking (254nm) with 400mJ/cm2 using a UV Stratalinker 2400 (Stratagene). Cells were then pelleted, lysed with RIPA buffer (50mM Tris-HCl pH 7.4; 150mM NaCl; 0.1% SDS; 0.5% sodium deoxycholate, 1% Nonidet P 40 substitute and protease inhibitors) and then sonicated. Cleared lysates were immunoprecipitated overnight at 4°C with anti-ZAP antibody (Abcam) and Dynabeads protein G (Invitrogen). Following 3 washes with RIPA buffer, beads were resuspended in 100 µl of RIPA and boiled for 10min to decouple protein/RNA complexes from the beads. Finally, input and pulldown samples were incubated with proteinase K (Thermo Fisher) for 1 hr at 37°C, and then boiled for 10 minutes to inactivate the enzyme. Samples were stored at -20°C for downstream RNA extraction and qRT-PCR analysis.

RNA from the input and pulldown samples was extracted using QIAzol (QIAGEN). cDNA synthesis was performed using the High-Capacity cDNA RT kit as described above. Of the final reaction, 5 µl were subjected to qPCR using the primer/probe sets for human GAPDH and HIV-1 genomic RNA reported in the STAR methods table. Serial dilutions of HIV-WT proviral DNA were used in standard curves as calibrator for quantification of HIV-1 RNA copies in the pull-down fractions.

### AP-1 reporter assay

One day prior to transfection, HEK293T cells were seeded in a 24-well plate at a density of 2×10^5^ cells per well. Approximately 8 h later, cells were treated with 50 ng/mL phorbol 12-myristate 13-acetate (PMA; Peprotech) and incubated overnight. The following day, cells were transfected with a DNA mix containing 85 ng of the AP-1 firefly luciferase reporter plasmid^116^ and 15 ng of the Renilla luciferase control plasmid (pGL4.74 hRluc/TK; Promega cat. #E6921) using 1 µL Lipofectamine 2000 (Invitrogen). Cells were incubated for 24 h and then lysed in 1x Passive Lysis Buffer (Promega). Dual-Luciferase Reporter Assay (Promega) was performed according to the manufacturer’s instructions, and luminescence was measured using a VICTOR X3 microplate reader (Perkin Elmer). Data was normalized as fold-change (ZAP knockout over control) by calculating the ratio of firefly to Renilla luminescence, with values from the control cell line set to 1.

### Immunofluorescence and RNA FISH

Cells were seeded on poly-L-lysine coated plates and infected with HIV-WT or HIV-CpG (MOI=1) then fixed in 3.7% (vol./vol.) formaldehyde at 24hpi. Cells were first immunostained for ZAP and the ER marker RTN4, then hybridized with Stellaris probes targeting unspliced HIV RNA (add ref https://doi.org/10.7554/eLife.62470) using Stellaris FISH buffers following the Stellaris protocol for sequential immunofluorescence (IF) + Stellaris RNA FISH in adherent cells (Biosearch Technologies). Images were acquired using a Nikon AXR NSPARC microscope at the Nikon Imaging Centre, King’s College London.

### ZAP and KHNYN iCLIP

Cells were infected at MOI=3 and media was replaced 18 h post-infection with DMEM supplemented with 2% fetal bovine serum. At 48 hours post-infection, cells were washed with ice-cold PBS, irradiated once with 150 mJ/cm2 in a Stratalinker 2400 at 254 nm, harvested by scraping in PBS, pelleted by centrifugation then resuspended in lysis buffer (50 mM Tris-HCl, pH 7.4; 100 mM NaCl; 1% Igepal CA-630; 0.1% SDS; 0.5% sodium deoxycholate; protease inhibitor) and stored at -80°C until use.

Individual cross-linking and immunoprecipitation (iCLIP) was performed according to the previously published protocol^117^, with the following details. For ZAP iCLIP, 4 µg of anti-ZAP antibody (GeneTex, GTX120134) was used for immunoprecipitation per sample. For the KHNYN KO, TRIM25 KO and ZAP-L KO samples, the membrane was cut above 110 kDa. In the ZAP-S KO experiments, the membrane was cut from 130 kDa upwards. For KHNYN-GFP iCLIP, 4 µg anti-GFP antibody (Roche, ref. 11814460001) was used for immunoprecipitation per sample. The membrane was cut from 130 kDa upwards. RNA-seq libraries were sequenced by Novogene as paired-end 150 bp reads on Illumina Novaseq 6000. Each iCLIP experiment was performed with three biological replicates.

### iCLIP data processing

Reads were mapped to the human GRCh38.p13 genome build with GENCODE v38 annotation. The HIV-WT or HIV-CpG, genome and annotation were merged with the human genome build and annotation files. Biological replicates were first analyzed individually to assess variability (standard deviation), after which the reads were pooled for all subsequent analyses. Pair-end reads were pre-processed using a custom script written in Bash. Briefly, the unique molecular identifiers (UMIs) were extracted using *UMI-tools extract*^118^ (https://github.com/CGATOxford/UMI-tools,--extract-method=regex) and the paired-end reads merged with *BBTools’ BBMerge*^119^ (https://sourceforge.net/projects/bbmap/, *mininsert=0, minoverlap=5*). The processed reads were analyzed according to the standardized iCLIP workflow with the *nf-core/clipseq v1.0.0* pipeline^120,121^ (https://doi.org/10.5281/zenodo.4723017), which was run with *Nextflow v22.10.0*^122^ and the following parameters: *--deduplicate true, --smrna_org human, --peakcaller paraclu*^123^. In addition, the crosslink BED files generated during the iCLIP analysis were used to call peaks on the HIV-1 genome with *Clippy v1.5.0* tool (https://github.com/ulelab/clippy). The following parameters were used: *--window_size 100, --min_prom_adjust 1, --min_height_adjust 1.5, --min_gene_counts 20, --min_peak_counts 5*.

To account for the variability in library size and RNA abundance across experiments, the crosslink signal was normalized in two steps. First, crosslink counts were scaled to counts per million (CPM) using the *clipplotr v1.0.0* command-line tool^124^ (https://github.com/ulelab/clipplotr) to adjust for differences in library size. Second, to control for variations in HIV-1 or cellular RNA abundance between samples, CPM-normalized crosslink counts were further adjusted using mean HIV-1 or specific cellular RNA abundance values, calculated from the DESeq2 normalized RNA-seq data across all three biological replicates. The standard error of the mean, shown as the error band on the iCLIP plots, was calculated from the standard deviation of three biological replicates. The data was smoothed using a rolling mean of 100.

Clippy (https://github.com/ulelab/clippy) was used for peak calling with rollmean100, minHeightAdjust1.5, minPromAdjust1, minGeneCount20, peakWidth0.9, and ≥ 0.25 crosslinks/nucleotide. For quantification of signal with specified peaks for HIV-WT or HIV-CpG in Figure S1H-I, the normalized crosslink data was summed for the peaks defined by Clippy for HIV-CpG in HEK293T cells (Peak 1 = nt 304-557, Peak 2 = nt 5650-6438 and Peak 3 = nt 7631-8443). For Figure S6C, the Clippy peaks for HIV-CpG in HeLa CRISPR control cells (Peak 1 = nt 258-535, Peak 2 = nt 5531-6497, Peak 3 = nt 7615-8454) were used to quantify the normalized crosslink data.

### RNA-seq

RNA was extracted using the Direct-zol RNA Miniprep Plus Kit (Zymo Pure) from cells grown and infected in the same conditions as the iCLIP experiments. Poly(A)+ RNA library preparation and sequencing were performed by GENEWIZ. Sequencing reads were processed using the *nf-core/rnaseq v3.17.0* pipeline^120^ (https://doi.org/10.5281/zenodo.13986791) run on *Nextflow v24.04.2*^122^ with default parameters. Differentially expressed genes were identified with DESeq2 using the *nf-core/differentialabundance v1.2.0* pipeline (https://doi.org/10.5281/zenodo.7846648). To account for transcript length bias, the analysis utilized the raw gene count matrix (salmon.merged.gene_counts.tsv) and transcript length matrix (salmon.merged.gene_lengths.tsv) generated by the *nf-core/rnaseq* pipeline. The pipeline was run with *Nextflow v24.04.2*^122^ using default settings. Differentially expressed genes were defined as adjusted p <0.05 and a basemean >500. In addition, ZAP-L regulated genes were defined by log2FC ZAP-L KO / HEK293T cells >0.5 and log2FC ZAP-L KO / ZAP-S KO cells >0.5. ZAP-S regulated genes were defined by log2FC ZAP-S KO / HEK293T cells >0.5 and log2FC ZAP-S KO / ZAP-L KO cells >0.5. ZAP, TRIM25 and KHNYN co-upregulated genes were defined by basemean >500, adjusted p <0.05 and log2FC > 0. SignalP 6.0^125^ was used to determine if any transcript encoded by ZAP-L or ZAP-S regulated genes in the GENCODE human basic gene annotation CHR was predicted to encode a signal peptide. The presence of transmembrane domains in specific genes was determined using information in the Localization section of GeneCards^126^.

Gene set enrichment analysis (GSEA) was performed using the R package clusterProfiler v4.12.6^127^ and human gene sets from the 50 Hallmark pathways obtained using the package msigdbr v7.5.1 (https://CRAN.R-project.org/package=msigdbr). GO enrichment analysis was performed using the DAVID Functional Annotation Tool with Gene Ontology Biological Process, Cellular Component, and Molecular Function Direct terms. The background included all genes with baseMean >500 in HEK293T CRISPR or HeLa CRISPR RNA-seq datasets. Only annotation terms with more than five genes were considered, and significant terms were defined as those with FDR <0.05. Gene expression relationships between different sample sets were analyzed and visualized using the R packages VennDiagram v1.7.3^128^ and eulerr v7.0.2 (https://CRAN.R-project.org/package=eulerr). The packages ggplot2 v3.5.1 (https://ggplot2.tidyverse.org) and pheatmap v1.0.12 (https://github.com/raivokolde/pheatmap) were used to create heat maps of gene expression.

### 5’ and 3’ RACE-seq

Total RNA was extracted using the Direct-zol RNA Miniprep Plus Kit. The 5’ RACE was performed from 1 µg total RNA using the SMARTer RACE 5’/3’ Kit according to the manufacturer’s instructions. To make cleaved RNA compatible with the 3’ RACE cDNA synthesis step, the total RNA was poly(A) tailed using *E. coli* Poly(A) polymerase (NEB) and then purified with the Monarch Spin RNA Cleanup Kit (NEB). The 3’ RACE was performed from 500 ng purified poly(A) tailed RNA using the SMARTer RACE 5’/3’ Kit according to the manufacturer’s instructions. The HIV specific primers used for 5’ and 3’ RACE are reported in the STAR methods table. RACE sequencing libraries were prepared using the Native Barcoding Kit 24 V14 (Oxford Nanopore Technologies). Sequencing of 5’ RACE product libraries was performed on R10.4.1 flowcells (FLO-MIN114, Oxford Nanopore, UK). 3’ RACE sequencing libraries were sequenced on PromethION with R10.4.1 flowcells.

For the 5’ RACE-seq, basecalling of *FAST5* files for the first replicate was completed using *guppy_basecaller* (v6.4.6). Dorado (v0.5.1) was used to basecall *POD5* files generated from 5’ RACE-seq replicates 2 and 3 as well as the 3’ RACE-seq replicates. Reads were basecalled and demultiplexed using the Super High Accuracy (SUP) model (v4.3.0). Additional command-line arguments were implemented to ensure that only reads with identical nanopore barcodes at both ends were considered, as well as excluding any reads containing barcodes in the middle. Bash scripts submitted to KCL’s CREATE HPC for basecalling are available on Github (5’ RACE-seq: https://github.com/Tycour/HIV1-ZAP_RACEseq; 3’ RACE-seq: https://github.com/GregaGD/ZAP_HIV-1_3-prime_RACE-seq), along with the bash scripts and custom Python and R scripts used in the downstream analysis.

Following basecalling, the FASTQ files were filtered with *NanoFilt v2.8.0*^129^ to eliminate low-quality reads and retain reads with an average quality score > Q20. The quality metrics of the filtered reads were evaluated using *NanoPlot v1.42.0*^129^. A custom Python script was implemented to identify and retain reads containing primer sequences at both ends that were used in the RACE reaction. For the 5’ RACE reaction, the 5’ primer was CTCACTATAGGGCAAGCAGTGGTATCAACGCAGAGTACATGGG and the 3’ primer was CCAGCAGCAGATGGGGTGGGAGCAGTAT. For the 3’ RACE reaction, the 5’ primer was TGGTGGGCGGGGATCAAGCAGGAATTTGGC and the 3’ primer was GTACTCTGCGTTGATACCACTGCTTGCCCTATAGTGAG. Reads without internal primer sequences were filtered out to ensure specificity in the downstream analysis.

Based on the infection conditions, the processed reads were aligned to either the HIV-WT or the HIV-CpG reference genome using *Minimap2*^130,131^ *v2.17-r941*, with the splice-aware argument (*--splice*). The alignment files in SAM format were converted to a sorted and indexed BAM format using *samtools (v1.10 or* v1.19). Each read was oriented to the leading or lagging strand using a custom Python script to facilitate downstream processing.

*Bedtools* (v2.31.1) converted the *BAM* file containing deduplicated reads to *BED* format. For the 5’ RACE-seq, an *awk* (v20200816) command was employed to extract the 5’ start positions for reads. These positions were then analyzed using a custom script that maps and plots the distribution of read starts along the reference genome. To examine the cleavage sites and detect non-templated nucleotides for the 3’ RACE-seq, the alignment files in BAM format were analyzed using a custom R script. Using the *GenomicAlignments* (Lawrence et al., 2013) and *Biostrings* (https://bioconductor.org/packages/Biostrings) packages, the final mapped position of the read was determined by parsing the CIGAR string. Specifically, the last mapped segment was extracted from the CIGAR string and the corresponding 3′ end nucleotide position was recorded. This was used in later steps to determine if any nucleotides beyond the mapped part of the read were non-templated.

Similarly, the *GenomicAlignments*^132^ and *Biostrings* (https://bioconductor.org/packages/Biostrings) packages were used to identify stretches of poly(A) nucleotides added during the 3’ RACE protocol. The 3′ end soft-clipped segment of the read was extracted based on the CIGAR string and analyzed to detect poly(A) segments of at least 15 consecutive adenosines, allowing up to two mismatches. The position of the poly(A) start in the read sequence was recorded and only reads with recognized poly(A) segments were selected for downstream analysis. The non-templated nucleotides were defined as the sequence between the final mapped position and the start of the poly(A) sequence in each read. The length distribution and nucleotide composition of these regions were analyzed, and the length distribution was visualized with bar charts, while the nucleotide composition per sequence length was visualized with stacked bar charts. The package *ggplot2 v3.5.1* (https://ggplot2.tidyverse.org) was used to generate all charts.

### Statistics

Unless otherwise stated, data are represented as mean ± standard deviation and statistical significance was determined by one-way ANOVA or unpaired t test from log-transformed data using GraphPad Prism10 (GraphPad Software Inc.). Significance was ascribed to p values < 0.05 compared to control conditions.

## RESOURCE AVAILABILITY

### Lead Contact

Additional information and resource requests will be fulfilled by the lead contact, Chad M. Swanson.

### Materials availability

No new reagents were generated in this study.

### Data and code availability

- The poly(A) RNA-seq, iCLIP-seq and RACE-seq data reported in this paper are available at ArrayExpress (E-MTAB-15071, E-MTAB-15072, E-MTAB-15077, E-MTAB-15065, E-MTAB-15056, E-MTAB-15061, E-MTAB-15060, E-MTAB-15095, E-MTAB-15064, E-MTAB-15097). These data are publicly available as of the date of publication.
- All original code has been deposited at Zenodo and is publicly available at https://doi.org/10.5281/zenodo.18832411 and https://zenodo.org/records/18839080 as of the date of publication. Other custom analysis scripts are available upon request.
- Any additional information required to reanalyse the data reported in this paper is available from the lead contact upon request.

## AUTHOR CONTRIBUTIONS

Conceptualization: C.M.S, S.J.D.N, J.U; Software: A.M.C.; Formal analysis: C.B., G.G.D., M.J.L, R.P.G, T.C., P.K., H.W., P.H., H.M., A.M.C., Investigation: C.B., G.G.D., M.J.L., R.P.G, P.K., H.W.; Writing - Original Draft: C.M.S, C.B., G.G.D., Writing - Review & Editing: all authors; Visualization: C.B., G.G.D, M.J.L, R.P.G., P.K., T.C.; Supervision: J.U., S.J.D.N, C.M.S.; Funding acquisition: M.P., J.U., S.J.D.N., C.M.S.

## ACKNOWLEDGEMENTS

We thank members of the Ule, Neil and Swanson laboratories for helpful discussions. The following reagents were obtained through the NIH AIDS Research and Reference Reagent Program, Division of AIDS, NIAID, NIH: TZM-bl from Dr. John C. Kappes, Dr. Xiaoyun Wu and Tranzyme Inc; HIV-1 p24 Hybridoma (183-H12-5C) from Dr. Bruce Chesebro. The Antiserum to HIV-1 gp120 #20 (ARP421) was obtained from the NIBSC Centre for AIDS Reagents. We thank Xavier Roca for the ZAP-L and ZAP-S knockout cells. We also thank Rebecca Sumner for the AP-1 reporter plasmid. We gratefully acknowledge the support of King’s Genomics, Social Genetic & Development Psychiatry Centre, Institute of Psychiatry, Psychology & Neuroscience, King’s College London for the Oxford Nanopore Technologies PromethION sequencing and the HPC CREATE at King’s College London. We also thank the Nikon Imaging Centre at Kings College London for help with light microscopy. These studies were funded by Medical Research Council grants MR/S000844/1 and MR/W018519/1 to SJDN and CMS, Wellcome Early Career Award (225081/Z/22/Z) to AMC, Sir Henry Dale Fellowship from the Wellcome Trust and the Royal Society (218537/Z/19/Z) to HEM, Wellcome Trust Senior Research Fellowship (WT098049AIA) to SJDN, the European Research Council (ERC) under the European Union’s Horizon 2020 research and innovation programme (grant agreement No 835300) to JU, and Slovenian Research and Innovation Agency ARIS grant P3-00083 to MP and GGD. The Slovenian Research Agency financially supported GGD through the Young Researcher Training Program for four years (2021–2025; contract number 802-7/2021-1).

## SUPPLEMENTAL FIGURE LEGENDS

**Figure S1: ZAP isoforms differ in their activity and their interaction with viral RNA, related to Figure 1**.

**(A)** Relative infectious virus production in isoform-specific HEK293T CRISPR ZAP cells. The dotted line represents HIV-CpG production in HEK293T cells. Error bars represent standard deviation.

**(B)** Immunoblotting of cell lysates and virions from HIV-WT- or HIV-CpG-infected isoform-specific HEK293T CRISPR ZAP cells.

**(C)** Quantification of KHNYN and TRIM25 abundance in the ZAP PULL-DOWN immunoblot from panel 1H. The signal was normalized ZAP abundance in the ZAP PULL-DOWN samples. Error bars represent standard deviation. *p <0.05 calculated by one-way ANOVA on log-transformed data, compared to HEK293T cells.

**(D-F)** Normalized ZAP iCLIP signal mapped to HIV gRNA in HEK293T cells **(D)**, HEK293T ZAP-L KO **(E)** and HEK293T ZAP-S KO **(F)**. The average of three independent experiments and the SEM (shading) are plotted.

**(G)** ZAP binding peaks on HIV-CpG RNA in 293T cells. Peaks were called by Clippy and their width are indicated.

**(H)** Quantification of normalized crosslink signal in Peak1 and Peak 3 for Figure S1D. Error bars represent standard deviation. *p <0.05 calculated by unpaired t test on log-transformed data, compared to HIV-WT.

**(I)** Quantification of normalized crosslink signal in Peak1, Peak 2 and Peak 3 for Figure 1I. Error bars represent standard deviation. *p <0.05 calculated by unpaired t test on log-transformed data, compared to 293T cells.

**Figure S2: ZAP-L and ZAP-S bind different motifs, target different cellular transcripts and have differential subcellular localization that is not affected by HIV-CpG infection, related to Figure 2.**

**(A)** UpA + CpG motifs ranked in the top 15 from the ZAP iCLIP experiments in ZAP-S knockout cells and their corresponding rank in HEK293T and ZAP-L KO cells.

**(B)** GO analysis for ZAP-L regulated genes with FDR <0.1 in HIV-WT infected cells. The FDR for the same terms for ZAP-S regulated genes are shown.

**(C)** GO analysis for ZAP-S regulated genes with FDR <0.1 in HIV-WT infected cells. The FDR for the same terms for ZAP-L regulated genes are shown.

**(D-F)** Normalized DESeq2 counts for JUNB **(D)**, JUND **(C)** and FOS **(F)** from poly(A) RNA-seq in HEK293T, ZAP-L and ZAP-S knockout cells. Error bars represent standard deviation. *adjp <0.05 calculated by DESeq2, compared to HEK293T cells.

**(G)** ZAP, ER (RTN4) and HIV-1 genomic RNA localization in mock or HIV-CpG infected HEK293T, ZAP-L KO or ZAP-S KO cells detected using indirect immunofluorescence for ZAP and RTN4 (an ER resident protein) plus RNA FISH for HIV-1 genomic RNA.

**Figure S3: ZAP binding peaks in TNFRSF10D have interspersed UpA and CpG dinucleotides, related to Figure 2.** ZAP binding peaks in TNFRSF10D identified in ZAP-S knockout cells using clippy. UpA dinucleotides are highlighted in yellow, CpG dinucleotides are highlighted in blue and UACG motifs are highlighted in green.

**Figure S4: HIV-1 infection increases the number of ZAP, TRIM25 or KHNYN-regulated genes, related to Figure 3 and 4**.

**(A)** Quantification of KHNYN abundance in the TRIM25 PULL-DOWN immunoblot from panel 3B. The signal was normalized to TRIM25 abundance in the TRIM25 PULL-DOWN samples. Error bars represent standard deviation.

**(B)** Heat map indicating differentially expressed genes that are upregulated in ZAP, TRIM25 or KHNYN CRISPR cells compared to CRISPR control cells in mock, HIV-WT or HIV-CpG infected cells.

**(C)** Number of upregulated genes in ZAP, KHNYN or TRIM25 knockout cells compared to CRISPR control cells in mock, HIV-WT or HIV-CpG infected cells.

**(D)** Euler diagrams showing differentially expressed genes that are upregulated in ZAP, TRIM25 or KHNYN CRISPR cells compared to CRISPR control cells in mock, HIV-WT or HIV-CpG infected cells.

**Figure S5: ZAP, TRIM25 and KHNYN co-regulate a subset of genes downregulated by HIV-1 infection and ZAP-L mRNA, related to Figure 4.**

**(A)** Euler diagram showing differentially expressed genes that are upregulated in ZAP, TRIM25 or KHNYN CRISPR cells compared to CRISPR control cells in mock, HIV-WT or HIV-CpG infected cells and downregulated by HIV-WT infection.

**(B)** Normalized DESeq2 counts for DIO2 from poly(A) RNA-seq in mock or HIV-WT infected Control, ZAP, TRIM25 or KHNYN knockout cells. Error bars represent standard deviation. *adjp <0.05 calculated by DESeq2, comparing HIV-WT infected Control CRISPR cells to mock infected Control CRISPR cells. *adjp <0.05 calculated by DESeq2, comparing HIV-WT infected ZAP, TRIM25 or KHNYN CRISPR cells to HIV-WT infected Control CRISPR cells.

**(C)** Normalized ZAP iCLIP signal mapped to DIO2 in HIV-WT infected Control CRISPR cells. The average signal of three independent experiments and the SEM (shading) are plotted. Exon tracks represent the isoforms expressed >1% of total normalized DESeq2 counts. The start and stop codon locations are indicated in pink.

**(D-E)** Normalized ZAP iCLIP signal mapped to ZAP-L **(D)** or ZAP-S **(E)** mRNA in HIV-WT infected Control CRISPR cells. The average signal of three independent experiments and the SEM (shading) are plotted. The normalized ZAP crosslink signal was distributed to ZAP-L or ZAP-S based on the isoform’s normalized DESeq2 count. Exons are indicated in grey and start and stop codon locations are in pink.

**(F)** Immunoblotting of mock, HIV-WT and HIV-CpG infected Control, ZAP, TRIM25 or KHNYN CRISPR cells.

**(G)** Relative infectious HIV-1 production (left) in Control, ZAP, TRIM25 and KHNYN CRISPR cells, and HIV-CpG production expressed as % infectious HIV-WT production (right). Error bars represent standard deviation. *p <0.05 calculated by one-way ANOVA on log-transformed data, compared to HIV-CpG production in Control CRISPR control cells.

**(H)** Normalized DESeq2 counts for ZAP-L mRNA from poly(A) RNA-seq in mock or HIV-WT infected Control, ZAP, TRIM25 or KHNYN knockout cells. Error bars represent standard deviation. *adjp <0.05 calculated by DESeq2, compared to Control CRISPR cells.

**Figure S6: TRIM25 modulates ZAP binding to HIV-CpG RNA but does not alter its target specificity, related to Figure 4.**

**(A)** Relative levels of ZAP-associated HIV-1 gRNA measured by qRT-PCR in HeLa Control CRISPR (left) and TRIM25 CRISPR (right) cells transfected with proviral DNA. Error bars represent standard deviation. *p <0.05 calculated by unpaired two-tailed t test, compared to ZAP-associated HIV-WT gRNA.

**(B)** ZAP iCLIP signal mapped to HIV-CpG gRNA in HeLa CRISPR TRIM25 and control. The average signal of three independent experiments and the SEM (shading) are plotted.

**(C)** Quantification of normalized crosslink signal in Peak1, Peak 2 and Peak 3 for Figure S6B. Error bars represent standard deviation. *p <0.05 calculated by unpaired t test on log-transformed data, compared to HIV-WT.

**(D)** MA plot showing genes with differential iCLIP read abundance in HIV-WT infected TRIM25 CRISPR cells compared to Control CRISPR cells.

**(E)** Normalized ZAP iCLIP signal mapped to ZAP in HIV-WT infected Control CRISPR cells. The average signal of three independent experiments and the SEM (shading) are plotted. Exon for ZAP-L and ZAP-S mRNAs are represented in grey and the start and stop codon locations are in pink.

**(F)** Heatmap showing the top 15 PEKA motifs from ZAP iCLIP experiments in CRISPR control cells and their relative rank in TRIM25 CRISPR cells.

**(G)** Euler diagram of genes encoding transcripts with ZAP peaks in HIV-WT infected CRISPR control and TRIM25 knockout cells.

**Figure S7: Depletion of CNOT1, PARN, EXOSC3, EXOSC5, EXOSC7, DDX17, DHX30 or DCP2 does not affect HIV-CpG infectious virus production, related to Figure 5**.

**(A)** Top 25 PEKA motifs for KHNYN iCLIP. CpG and UpA motifs are highlighted in red.

**(B)** Infectious HIV-CpG production, expressed as % infectious HIV-WT production in HeLa cells transfected with siRNA targeting ZAP, TRIM25 or KHNYN. The dotted line represents HIV-CpG production in cells transfected with a non-targeting control siRNA. Error bars represent standard deviation. *p <0.05 calculated by one-way ANOVA on log-transformed data, compared to HIV-CpG production in cells treated with a non-targeting siRNA control.

**(C-D)** Infectious HIV-CpG production, expressed as % infectious HIV-WT production in HeLa cells transfected with siRNA targeting potential ZAP cofactors. The dotted line represents HIV-CpG production in cells transfected with a non-targeting control siRNA. Error bars represent standard deviation. *p <0.05 calculated by one-way ANOVA on log-transformed data, compared to HIV-CpG production in cells treated with a non-targeting siRNA control.

**(E)** Immunoblotting of HeLa cells transfected with siRNA targeting the genes in (B) and (C), then infected with HIV-WT.

**(F-G)** Level of HIV-CpG 3’ cleavage fragment measured by qRT-PCR in cells transfected with siRNA targeting potential ZAP cofactors. Results are expressed as % 3’ cleavage fragment in HIV-WT infected cells. The dotted line represents the level of 3’ cleavage fragment in cells treated with a non-targeting siRNA control. Error bars represent standard deviation. *p <0.05 calculated by one-way ANOVA on log-transformed data, compared to the level of 3’ cleavage fragment in cells treated with a non-targeting siRNA control.

**(H)** Quantification of XRN1 abundance in the TRIM25 PULL-DOWN immunoblot from panel 5E. The signal was normalized TRIM25 abundance in the TRIM25 PULL-DOWN samples. Error bars represent standard deviation.

**(I)** Immunoblotting of ZAP coimmunoprecipitations and corresponding input lysates obtained from infected Control and TRIM25 CRISPR HeLa cells. Samples were treated with RNase I after pulldown. Note that the experiment shown here was performed alongside the experiment shown in Figure S10G, thus the input blots for PABPC1, TRIM25, ZAP and GAPDH are identical between the two panels. The blots are representative of three biological replicates.

**Figures S8: Clustered endonuclease cleavage sites correlate with high levels of ZAP binding, related to Figure 5**

**(A)** Normalized ZAP crosslink signal (top panel), KHNYN crosslink signal (middle panel) and HIV-CpG / HIV-WT fold change in 5’ end coverage in XRN1 knockout cells (bottom panel).

**(B)** HIV-CpG sequence nt 6000-6700. CpG is highlighted in blue, UpA is highlighted in yellow and UACG is highlighted in green. Cleavage sites are indicated by a red slash.

**Figure S9: Depletion of CNOT1, PARN, EXOSC3, EXOSC5, EXOSC7, XRN1, DDX17, or DHX30 does not affect the abundance of the 5’ HIV-CpG RNA cleavage fragment, related to Figure 6**.

**(A)** Genomic organization of HIV-WT and HIV-CpG and localization of CpG dinucleotides. The qRT-PCR primer/probe set used to quantify the 5’ cleavage fragment (probe 1 (p1)) and the HIV-specific primer used for the 3’ RACE (3’R) are shown.

**(B)** Level of HIV-CpG 5’ cleavage fragment measured by qRT-PCR in cells transfected with siRNA targeting potential ZAP cofactors. Results are expressed as % 5’ cleavage fragment in HIV-WT infected cells. The dotted line represents the level of 5’ cleavage fragment in cells treated with a non-targeting siRNA control. Error bars represent standard deviation. *p <0.05 calculated by one-way ANOVA on log-transformed data, compared to the level of 3’ cleavage fragment in cells treated with a non-targeting siRNA control.

**(C)** Immunoblotting of HeLa cells transfected with siRNA targeting TUT4 and TUT7 then infected with HIV-WT.

**(D)** Infectious HIV-CpG production, expressed as % infectious HIV-WT production in HeLa cells transfected with siRNA targeting TUTs. The dotted line represents HIV-CpG production in cells transfected with a non-targeting control siRNA. Error bars represent standard deviation. *p <0.05 calculated by one-way ANOVA on log-transformed data, compared to HIV-CpG production in cells treated with a non-targeting siRNA control.

**(E)** Level of HIV-CpG 3’ cleavage fragment measured by qRT-PCR in cells transfected with siRNA targeting TUTs. Results are expressed as % 3’ cleavage fragment in HIV-WT infected cells. The dotted line represents the level of 3’ cleavage fragment in cells treated with a non-targeting siRNA control. Error bars represent standard deviation. *p <0.05 calculated by one-way ANOVA on log-transformed data, compared to the level of 3’ cleavage fragment in cells treated with a non-targeting siRNA control.

**(F)** Immunoblotting of ZAP coimmunoprecipitations and corresponding input lysates obtained from infected HeLa cells. Samples were treated with RNase I after pulldown. The blots are representative of three biological replicates.

**(G)** Immunoblotting of TRIM25 coimmunoprecipitations and corresponding input lysates obtained from infected HeLa cells. Samples were treated with RNase I after pulldown. The blots are representative of three biological replicates.

**(H)** Quantification of TUT7 abundance in the TRIM25 PULL-DOWN immunoblot from panel S6G. The signal was normalized to TRIM25 abundance in the TRIM25 PULL-DOWN samples. Error bars represent standard deviation. *p <0.05 calculated by one-way ANOVA on log-transformed data.

**(I)** Immunoblotting of ZAP coimmunoprecipitations and corresponding input lysates obtained from infected Control and TRIM25 CRISPR HeLa cells. Samples were treated with RNase I after pulldown. Note that the experiment shown here was performed alongside the experiment shown in Figure 3C, thus the input blots for PABPC1, TRIM25, ZAP and GAPDH are identical between the two panels. The blots are representative of three biological replicates.

**Figure S10: Depletion of DIS3L2 does not affect HIV-CpG infectious virus production or abundance of the 3’ cleavage fragment, related to Figure 6**.

**(A)** Immunoblotting of HeLa cells transfected with siRNA targeting DIS3L2 then infected with HIV-WT.

**(B)** Level of HIV-CpG 3’ cleavage fragment measured by qRT-PCR in cells transfected with siRNA targeting DIS3L2. Results are expressed as % 3’ cleavage fragment in HIV-WT infected cells. The dotted line represents the level of 3’ cleavage fragment in cells treated with a non-targeting siRNA control. Error bars represent standard deviation. *p <0.05 calculated by one-way ANOVA on log-transformed data, compared to the level of 5’ cleavage fragment in cells treated with a non-targeting siRNA control. This experiment was performed alongside the experiment shown in figure S9E and therefore shared the same control and ZAP values.

**(C)** Infectious HIV-CpG production, expressed as % infectious HIV-WT production in HeLa cells transfected with siRNA targeting DIS3L2. The dotted line represents HIV-CpG production in cells transfected with a non-targeting control siRNA. Error bars represent standard deviation. *p <0.05 calculated by one-way ANOVA on log-transformed data, compared to HIV-CpG production in cells treated with a non-targeting siRNA control. This experiment was performed alongside the experiment shown in figure 9E and therefore shared the same control and ZAP values.

**(D)** Immunoblotting of ZAP coimmunoprecipitations and corresponding input lysates obtained from infected HeLa cells. Samples were treated with RNase I after pulldown. The blots are representative of three biological replicates.

**(E)** Immunoblotting of TRIM25 coimmunoprecipitations and corresponding input lysates obtained from infected HeLa cells. Samples were treated with RNase I after pulldown. The blots are representative of three biological replicates.

**(F)** Quantification of DIS3L2 abundance in the TRIM25 PULL-DOWN immunoblot from panel S10E. The signal was normalized to TRIM25 abundance in the TRIM25 PULL-DOWN samples. Error bars represent standard deviation.

**(G)** Immunoblotting of ZAP coimmunoprecipitations and corresponding input lysates obtained from infected Control and TRIM25 CRISPR HeLa cells. Samples were treated with RNase I after pulldown. Note that the experiment shown here was performed alongside the experiment shown in Figure 7I, thus the input blots for PABPC1, TRIM25, ZAP and GAPDH are identical between the two panels. The blots are representative of three biological replicates.

## SUPPLEMENTARY DATASETS

**Table S1: poly(A)RNA-seq in mock, HIV-WT and HIV-CpG infected HEK293T, ZAP-L and ZAP-S knockout cells, related to Figure 1 and 2**.

**Table S2: ZAP binding peaks in HIV-WT and HIV-CpG infected HEK293T, ZAP-L and ZAP-S knockout cells, related to Figure 1 and 2**.

**Table S3: poly(A)RNA-seq in mock, HIV-WT and HIV-CpG infected Control, ZAP, TRIM25 and KHNYN CRISPR cells, related to Figure 3 and 4**.

**Table S4: ZAP binding peaks in HIV-WT and HIV-CpG infected Control and TRIM25 CRISPR cells, related to Figure S6.**

## REFERENCES

1. Chan, C.P., and Jin, D.Y. (2022). Cytoplasmic RNA sensors and their interplay with RNA-binding partners in innate antiviral response: theme and variations. RNA 28, 449–477. 10.1261/rna.079016.121.

2. Yang, E., and Li, M.M.H. (2020). All About the RNA: Interferon-Stimulated Genes That Interfere With Viral RNA Processes. Front Immunol 11, 605024. 10.3389/fimmu.2020.605024.

3. Gao, G., Guo, X., and Goff, S.P. (2002). Inhibition of retroviral RNA production by ZAP, a CCCH-type zinc finger protein. Science 297, 1703–1706. 10.1126/science.1074276.

4. Ficarelli, M., Neil, S.J.D., and Swanson, C.M. (2021). Targeted Restriction of Viral Gene Expression and Replication by the ZAP Antiviral System. Annu Rev Virol 8, 265–283. 10.1146/annurev-virology-091919-104213.

5. Shao, R., Visser, I., Fros, J.J., and Yin, X. (2024). Versatility of the Zinc-Finger Antiviral Protein (ZAP) As a Modulator of Viral Infections. Int J Biol Sci 20, 4585–4600. 10.7150/ijbs.98029.

6. Bick, M.J., Carroll, J.W., Gao, G., Goff, S.P., Rice, C.M., and MacDonald, M.R. (2003). Expression of the zinc-finger antiviral protein inhibits alphavirus replication. J Virol 77, 11555–11562. 10.1128/jvi.77.21.11555-11562.2003.

7. Zhu, Y., Wang, X., Goff, S.P., and Gao, G. (2012). Translational repression precedes and is required for ZAP-mediated mRNA decay. EMBO J 31, 4236–4246. 10.1038/emboj.2012.271.

8. Zimmer, M.M., Kibe, A., Rand, U., Pekarek, L., Ye, L., Buck, S., Smyth, R.P., Cicin-Sain, L., and Caliskan, N. (2021). The short isoform of the host antiviral protein ZAP acts as an inhibitor of SARS-CoV-2 programmed ribosomal frameshifting. Nature communications 12, 7193. 10.1038/s41467-021-27431-0.

9. Takata, M.A., Goncalves-Carneiro, D., Zang, T.M., Soll, S.J., York, A., Blanco-Melo, D., and Bieniasz, P.D. (2017). CG dinucleotide suppression enables antiviral defence targeting non-self RNA. Nature 550, 124–127. 10.1038/nature24039.

10. Meagher, J.L., Takata, M., Goncalves-Carneiro, D., Keane, S.C., Rebendenne, A., Ong, H., Orr, V.K., MacDonald, M.R., Stuckey, J.A., Bieniasz, P.D., and Smith, J.L. (2019). Structure of the zinc-finger antiviral protein in complex with RNA reveals a mechanism for selective targeting of CG-rich viral sequences. Proc Natl Acad Sci U S A 116, 24303–24309. 10.1073/pnas.1913232116.

11. Odon, V., Fros, J.J., Goonawardane, N., Dietrich, I., Ibrahim, A., Alshaikhahmed, K., Nguyen, D., and Simmonds, P. (2019). The role of ZAP and OAS3/RNAseL pathways in the attenuation of an RNA virus with elevated frequencies of CpG and UpA dinucleotides. Nucleic Acids Res 47, 8061–8083. 10.1093/nar/gkz581.

12. Luo, X., Wang, X., Gao, Y., Zhu, J., Liu, S., Gao, G., and Gao, P. (2020). Molecular Mechanism of RNA Recognition by Zinc-Finger Antiviral Protein. Cell reports 30, 46–52 e44. 10.1016/j.celrep.2019.11.116.

13. Fros, J.J., Visser, I., Tang, B., Yan, K., Nakayama, E., Visser, T.M., Koenraadt, C.J.M., van Oers, M.M., Pijlman, G.P., Suhrbier, A., and Simmonds, P. (2021). The dinucleotide composition of the Zika virus genome is shaped by conflicting evolutionary pressures in mammalian hosts and mosquito vectors. PLoS Biol 19, e3001201. 10.1371/journal.pbio.3001201.

14. Sharp, C.P., Thompson, B.H., Nash, T.J., Diebold, O., Pinto, R.M., Thorley, L., Lin, Y.T., Sives, S., Wise, H., Clohisey Hendry, S., et al. (2023). CpG dinucleotide enrichment in the influenza A virus genome as a live attenuated vaccine development strategy. PLoS Pathog 19, e1011357. 10.1371/journal.ppat.1011357.

15. Le, N.P.K., Singh, P.P., Trus, I., and Karniychuk, U. (2025). West Nile virus vaccine candidates attenuated by dinucleotide enrichment are immunogenic and protective against lethal infection. PLoS Pathog 21, e1013560. 10.1371/journal.ppat.1013560.

16. Goonawardane, N., Nguyen, D., and Simmonds, P. (2021). Association of Zinc Finger Antiviral Protein Binding to Viral Genomic RNA with Attenuation of Replication of Echovirus 7. mSphere 6, e01138–01120. 10.1128/mSphere.01138-20.

17. Goncalves-Carneiro, D., Mastrocola, E., Lei, X., DaSilva, J., Chan, Y.F., and Bieniasz, P.D. (2022). Rational attenuation of RNA viruses with zinc finger antiviral protein. Nat Microbiol 7, 1558–1567. 10.1038/s41564-022-01223-8.

18. Karlin, S., Doerfler, W., and Cardon, L.R. (1994). Why is CpG suppressed in the genomes of virtually all small eukaryotic viruses but not in those of large eukaryotic viruses? J Virol 68, 2889–2897. 10.1128/JVI.68.5.2889-2897.1994.

19. Guo, X., Carroll, J.W., Macdonald, M.R., Goff, S.P., and Gao, G. (2004). The zinc finger antiviral protein directly binds to specific viral mRNAs through the CCCH zinc finger motifs. J Virol 78, 12781–12787. 10.1128/JVI.78.23.12781-12787.2004.

20. Chen, S., Xu, Y., Zhang, K., Wang, X., Sun, J., Gao, G., and Liu, Y. (2012). Structure of N-terminal domain of ZAP indicates how a zinc-finger protein recognizes complex RNA. Nat Struct Mol Biol 19, 430–435. 10.1038/nsmb.2243.

21. Ficarelli, M., Antzin-Anduetza, I., Hugh-White, R., Firth, A.E., Sertkaya, H., Wilson, H., Neil, S.J.D., Schulz, R., and Swanson, C.M. (2020). CpG Dinucleotides Inhibit HIV-1 Replication through Zinc Finger Antiviral Protein (ZAP)-Dependent and -Independent Mechanisms. J Virol 94, e01337–01319. 10.1128/JVI.01337-19.

22. Groenke, N., Trimpert, J., Merz, S., Conradie, A.M., Wyler, E., Zhang, H., Hazapis, O.G., Rausch, S., Landthaler, M., Osterrieder, N., and Kunec, D. (2020). Mechanism of Virus Attenuation by Codon Pair Deoptimization. Cell reports 31, 107586. 10.1016/j.celrep.2020.107586.

23. Nguyen, L.P., Aldana, K.S., Yang, E., Yao, Z., and Li, M.M.H. (2023). Alphavirus Evasion of Zinc Finger Antiviral Protein (ZAP) Correlates with CpG Suppression in a Specific Viral nsP2 Gene Sequence. Viruses 15. 10.3390/v15040830.

24. Kmiec, D., Nchioua, R., Sherrill-Mix, S., Sturzel, C.M., Heusinger, E., Braun, E., Gondim, M.V.P., Hotter, D., Sparrer, K.M.J., Hahn, B.H., et al. (2020). CpG Frequency in the 5’ Third of the env Gene Determines Sensitivity of Primary HIV-1 Strains to the Zinc-Finger Antiviral Protein. mBio 11, e02903–02919. 10.1128/mBio.02903-19.

25. Sertkaya, H., Hidalgo, L., Ficarelli, M., Kmiec, D., Signell, A.W., Ali, S., Parker, H., Wilson, H., Neil, S.J.D., Malim, M.H., et al. (2021). Minimal impact of ZAP on lentiviral vector production and transduction efficiency. Mol Ther Methods Clin Dev 23, 147–157. 10.1016/j.omtm.2021.08.008.

26. Todorova, T., Bock, F.J., and Chang, P. (2014). PARP13 regulates cellular mRNA post-transcriptionally and functions as a pro-apoptotic factor by destabilizing TRAILR4 transcript. Nature communications 5, 5362. 10.1038/ncomms6362.

27. Ly, P.T., Xu, S., Wirawan, M., Luo, D., and Roca, X. (2022). ZAP isoforms regulate unfolded protein response and epithelial-mesenchymal transition. Proc Natl Acad Sci U S A 119, e2121453119. 10.1073/pnas.2121453119.

28. Goncalves-Carneiro, D., Mastrocola, E., Lei, X., and Bieniasz, P.D. (2025). Modulation of host gene expression by the zinc finger antiviral protein. Proc Natl Acad Sci U S A 122, e2420819122. 10.1073/pnas.2420819122.

29. Busa, V.F., Ando, Y., Aigner, S., Yee, B.A., Yeo, G.W., and Leung, A.K.L. (2024). Transcriptome regulation by PARP13 in basal and antiviral states in human cells. iScience 27, 109251. 10.1016/j.isci.2024.109251.

30. Cooper, D.N., and Gerber-Huber, S. (1985). DNA methylation and CpG suppression. Cell Differ 17, 199–205. 10.1016/0045-6039(85)90488-9.

31. Shaw, A.E., Rihn, S.J., Mollentze, N., Wickenhagen, A., Stewart, D.G., Orton, R.J., Kuchi, S., Bakshi, S., Collados, M.R., Turnbull, M.L., et al. (2021). The antiviral state has shaped the CpG composition of the vertebrate interferome to avoid self-targeting. PLoS Biol 19, e3001352. 10.1371/journal.pbio.3001352.

32. Nchioua, R., Kmiec, D., Krchlikova, V., Mattes, S., Noettger, S., Bibollet-Ruche, F., Russell, R.M., Sparrer, K.M.J., Charpentier, T., Tardy, F., et al. (2025). Host ZAP activity correlates with the levels of CpG suppression in primate lentiviruses. Proc Natl Acad Sci U S A 122, e2419489122. 10.1073/pnas.2419489122.

33. Guo, X., Ma, J., Sun, J., and Gao, G. (2007). The zinc-finger antiviral protein recruits the RNA processing exosome to degrade the target mRNA. Proc Natl Acad Sci U S A 104, 151–156. 10.1073/pnas.0607063104.

34. Zhu, Y., Chen, G., Lv, F., Wang, X., Ji, X., Xu, Y., Sun, J., Wu, L., Zheng, Y.T., and Gao, G. (2011). Zinc-finger antiviral protein inhibits HIV-1 infection by selectively targeting multiply spliced viral mRNAs for degradation. Proc Natl Acad Sci U S A 108, 15834–15839. 10.1073/pnas.1101676108.

35. Chen, G., Guo, X., Lv, F., Xu, Y., and Gao, G. (2008). p72 DEAD box RNA helicase is required for optimal function of the zinc-finger antiviral protein. Proc Natl Acad Sci U S A 105, 4352–4357. 10.1073/pnas.0712276105.

36. Lee, H., Komano, J., Saitoh, Y., Yamaoka, S., Kozaki, T., Misawa, T., Takahama, M., Satoh, T., Takeuchi, O., Yamamoto, N., et al. (2013). Zinc-finger antiviral protein mediates retinoic acid inducible gene I-like receptor-independent antiviral response to murine leukemia virus. Proc Natl Acad Sci U S A 110, 12379–12384. 10.1073/pnas.1310604110.

37. Ye, P., Liu, S., Zhu, Y., Chen, G., and Gao, G. (2010). DEXH-Box protein DHX30 is required for optimal function of the zinc-finger antiviral protein. Protein Cell 1, 956–964. 10.1007/s13238-010-0117-8.

38. Li, M.M., Lau, Z., Cheung, P., Aguilar, E.G., Schneider, W.M., Bozzacco, L., Molina, H., Buehler, E., Takaoka, A., Rice, C.M., et al. (2017). TRIM25 Enhances the Antiviral Action of Zinc-Finger Antiviral Protein (ZAP). PLoS Pathog 13, e1006145. 10.1371/journal.ppat.1006145.

39. Zheng, X., Wang, X., Tu, F., Wang, Q., Fan, Z., and Gao, G. (2017). TRIM25 Is Required for the Antiviral Activity of Zinc Finger Antiviral Protein. J Virol 91, e00088–00017. 10.1128/JVI.00088-17.

40. Ficarelli, M., Wilson, H., Pedro Galao, R., Mazzon, M., Antzin-Anduetza, I., Marsh, M., Neil, S.J., and Swanson, C.M. (2019). KHNYN is essential for the zinc finger antiviral protein (ZAP) to restrict HIV-1 containing clustered CpG dinucleotides. Elife 8, e46767. 10.7554/eLife.46767.

41. Bohn, J.A., Meagher, J.L., Takata, M.A., Goncalves-Carneiro, D., Yeoh, Z.C., Ohi, M.D., Smith, J.L., and Bieniasz, P.D. (2024). Functional anatomy of zinc finger antiviral protein complexes. Nature communications 15, 10834. 10.1038/s41467-024-55192-z.

42. Yeoh, Z.C., Meagher, J.L., Kang, C.Y., Bieniasz, P.D., Smith, J.L., and Ohi, M.D. (2024). A minimal complex of KHNYN and zinc-finger antiviral protein binds and degrades single-stranded RNA. Proc Natl Acad Sci U S A 121, e2415048121. 10.1073/pnas.2415048121.

43. OhAinle, M., Helms, L., Vermeire, J., Roesch, F., Humes, D., Basom, R., Delrow, J.J., Overbaugh, J., and Emerman, M. (2018). A virus-packageable CRISPR screen identifies host factors mediating interferon inhibition of HIV. Elife 7, e39823. 10.7554/eLife.39823.

44. Youle, R.L., Lista, M.J., Bouton, C., Kunzelmann, S., Wilson, H., Cottee, M.A., Purkiss, A.G., Morris, E.R., Neil, S.J.D., Taylor, I.A., and Swanson, C.M. (2025). Structural and functional characterization of the extended-diKH domain from the antiviral endoribonuclease KHNYN. J Biol Chem, 108336. 10.1016/j.jbc.2025.108336.

45. Kerns, J.A., Emerman, M., and Malik, H.S. (2008). Positive selection and increased antiviral activity associated with the PARP-containing isoform of human zinc-finger antiviral protein. PLoS genetics 4, e21. 10.1371/journal.pgen.0040021.

46. Li, M.M.H., Aguilar, E.G., Michailidis, E., Pabon, J., Park, P., Wu, X., de Jong, Y.P., Schneider, W.M., Molina, H., Rice, C.M., and MacDonald, M.R. (2019). Characterization of Novel Splice Variants of Zinc Finger Antiviral Protein (ZAP). J Virol 93, e00715–00719. 10.1128/JVI.00715-19.

47. Hayakawa, S., Shiratori, S., Yamato, H., Kameyama, T., Kitatsuji, C., Kashigi, F., Goto, S., Kameoka, S., Fujikura, D., Yamada, T., et al. (2011). ZAPS is a potent stimulator of signaling mediated by the RNA helicase RIG-I during antiviral responses. Nat Immunol 12, 37–44. 10.1038/ni.1963.

48. Nchioua, R., Kmiec, D., Muller, J.A., Conzelmann, C., Gross, R., Swanson, C.M., Neil, S.J.D., Stenger, S., Sauter, D., Munch, J., et al. (2020). SARS-CoV-2 Is Restricted by Zinc Finger Antiviral Protein despite Preadaptation to the Low-CpG Environment in Humans. mBio 11, e01930–01920. 10.1128/mBio.01930-20.

49. Xue, G., Braczyk, K., Goncalves-Carneiro, D., Dawidziak, D.M., Sanchez, K., Ong, H., Wan, Y., Zadrozny, K.K., Ganser-Pornillos, B.K., Bieniasz, P.D., and Pornillos, O. (2022). Poly(ADP-ribose) potentiates ZAP antiviral activity. PLoS Pathog 18, e1009202. 10.1371/journal.ppat.1009202.

50. Kuttiyatveetil, J.R.A., Soufari, H., Dasovich, M., Uribe, I.R., Mirhasan, M., Cheng, S.J., Leung, A.K.L., and Pascal, J.M. (2022). Crystal structures and functional analysis of the ZnF5-WWE1-WWE2 region of PARP13/ZAP define a distinctive mode of engaging poly(ADP-ribose). Cell reports 41, 111529. 10.1016/j.celrep.2022.111529.

51. Charron, G., Li, M.M., MacDonald, M.R., and Hang, H.C. (2013). Prenylome profiling reveals S-farnesylation is crucial for membrane targeting and antiviral activity of ZAP long-isoform. Proc Natl Acad Sci U S A 110, 11085–11090. 10.1073/pnas.1302564110.

52. Schwerk, J., Soveg, F.W., Ryan, A.P., Thomas, K.R., Hatfield, L.D., Ozarkar, S., Forero, A., Kell, A.M., Roby, J.A., So, L., et al. (2019). RNA-binding protein isoforms ZAP-S and ZAP-L have distinct antiviral and immune resolution functions. Nat Immunol 20, 1610–1620. 10.1038/s41590-019-0527-6.

53. Kmiec, D., Lista, M.J., Ficarelli, M., Swanson, C.M., and Neil, S.J.D. (2021). S-farnesylation is essential for antiviral activity of the long ZAP isoform against RNA viruses with diverse replication strategies. PLoS Pathog 17, e1009726. 10.1371/journal.ppat.1009726.

54. Kim, M., Pyo, Y., Hyun, S.I., Jeong, M., Choi, Y., and Kim, V.N. (2025). Exogenous RNA surveillance by proton-sensing TRIM25. Science 388, eads4539. 10.1126/science.ads4539.

55. Lin, Y., Zhang, H., Liang, J., Li, K., Zhu, W., Fu, L., Wang, F., Zheng, X., Shi, H., Wu, S., et al. (2014). Identification and characterization of alphavirus M1 as a selective oncolytic virus targeting ZAP-defective human cancers. Proc Natl Acad Sci U S A 111, E4504–4512. 10.1073/pnas.1408759111.

56. Kypr, J., Mrazek, J., and Reich, J. (1989). Nucleotide composition bias and CpG dinucleotide content in the genomes of HIV and HTLV 1/2. Biochim Biophys Acta 1009, 280–282. 10.1016/0167-4781(89)90114-0.

57. Shpaer, E.G., and Mullins, J.I. (1990). Selection against CpG dinucleotides in lentiviral genes: a possible role of methylation in regulation of viral expression. Nucleic Acids Res 18, 5793–5797. 10.1093/nar/18.19.5793.

58. Wasson, M.K., Borkakoti, J., Kumar, A., Biswas, B., and Vivekanandan, P. (2017). The CpG dinucleotide content of the HIV-1 envelope gene may predict disease progression. Sci Rep 7, 8162. 10.1038/s41598-017-08716-1.

59. Theys, K., Feder, A.F., Gelbart, M., Hartl, M., Stern, A., and Pennings, P.S. (2018). Within-patient mutation frequencies reveal fitness costs of CpG dinucleotides and drastic amino acid changes in HIV. PLoS genetics 14, e1007420. 10.1371/journal.pgen.1007420.

60. Goncalves-Carneiro, D., Takata, M.A., Ong, H., Shilton, A., and Bieniasz, P.D. (2021). Origin and evolution of the zinc finger antiviral protein. PLoS Pathog 17, e1009545. 10.1371/journal.ppat.1009545.

61. Lista, M.J., Ficarelli, M., Wilson, H., Kmiec, D., Youle, R.L., Wanford, J., Winstone, H., Odendall, C., Taylor, I.A., Neil, S.J.D., and Swanson, C.M. (2023). A Nuclear Export Signal in KHNYN Required for Its Antiviral Activity Evolved as ZAP Emerged in Tetrapods. J Virol 97, e0087222. 10.1128/jvi.00872-22.

62. Konig, J., Zarnack, K., Rot, G., Curk, T., Kayikci, M., Zupan, B., Turner, D.J., Luscombe, N.M., and Ule, J. (2010). iCLIP reveals the function of hnRNP particles in splicing at individual nucleotide resolution. Nat Struct Mol Biol 17, 909–915. 10.1038/nsmb.1838.

63. Keane, S.C., and Summers, M.F. (2016). NMR Studies of the Structure and Function of the HIV-1 5’-Leader. Viruses 8, 338. 10.3390/v8120338.

64. Sertznig, H., Hillebrand, F., Erkelenz, S., Schaal, H., and Widera, M. (2018). Behind the scenes of HIV-1 replication: Alternative splicing as the dependency factor on the quiet. Virology 516, 176–188. 10.1016/j.virol.2018.01.011.

65. Law, L.M., Albin, O.R., Carroll, J.W., Jones, C.T., Rice, C.M., and Macdonald, M.R. (2010). Identification of a dominant negative inhibitor of human zinc finger antiviral protein reveals a functional endogenous pool and critical homotypic interactions. J Virol 84, 4504–4512. 10.1128/JVI.02018-09.

66. Hamid, F.M., and Makeyev, E.V. (2016). Exaptive origins of regulated mRNA decay in eukaryotes. Bioessays 38, 830–838. 10.1002/bies.201600100.

67. Kuret, K., Amalietti, A.G., Jones, D.M., Capitanchik, C., and Ule, J. (2022). Positional motif analysis reveals the extent of specificity of protein-RNA interactions observed by CLIP. Genome Biol 23, 191. 10.1186/s13059-022-02755-2.

68. Leung, A.K., Vyas, S., Rood, J.E., Bhutkar, A., Sharp, P.A., and Chang, P. (2011). Poly(ADP-ribose) regulates stress responses and microRNA activity in the cytoplasm. Mol Cell 42, 489–499. 10.1016/j.molcel.2011.04.015.

69. Seo, G.J., Kincaid, R.P., Phanaksri, T., Burke, J.M., Pare, J.M., Cox, J.E., Hsiang, T.Y., Krug, R.M., and Sullivan, C.S. (2013). Reciprocal inhibition between intracellular antiviral signaling and the RNAi machinery in mammalian cells. Cell Host Microbe 14, 435–445. 10.1016/j.chom.2013.09.002.

70. Simmonds, P., Xia, W., Baillie, J.K., and McKinnon, K. (2013). Modelling mutational and selection pressures on dinucleotides in eukaryotic phyla--selection against CpG and UpA in cytoplasmically expressed RNA and in RNA viruses. BMC genomics 14, 610. 10.1186/1471-2164-14-610.

71. Gonzalez-Perez, A.C., Stempel, M., Wyler, E., Urban, C., Piras, A., Hennig, T., Ganskih, S., Wei, Y., Heim, A., Landthaler, M., et al. (2021). The Zinc Finger Antiviral Protein ZAP Restricts Human Cytomegalovirus and Selectively Binds and Destabilizes Viral UL4/UL5 Transcripts. mBio 12. 10.1128/mBio.02683-20.

72. Lin, Y.T., Chiweshe, S., McCormick, D., Raper, A., Wickenhagen, A., DeFillipis, V., Gaunt, E., Simmonds, P., Wilson, S.J., and Grey, F. (2020). Human cytomegalovirus evades ZAP detection by suppressing CpG dinucleotides in the major immediate early 1 gene. PLoS Pathog 16, e1008844. 10.1371/journal.ppat.1008844.

73. Lista, M.J., Witney, A.A., Nichols, J., Davison, A.J., Wilson, H., Latham, K.A., Ravenhill, B.J., Nightingale, K., Stanton, R.J., Weekes, M.P., et al. (2023). Strain-Dependent Restriction of Human Cytomegalovirus by Zinc Finger Antiviral Proteins. J Virol, e0184622. 10.1128/jvi.01846-22.

74. Galao, R.P., Wilson, H., Schierhorn, K.L., Debeljak, F., Bodmer, B.S., Goldhill, D., Hoenen, T., Wilson, S.J., Swanson, C.M., and Neil, S.J.D. (2022). TRIM25 and ZAP target the Ebola virus ribonucleoprotein complex to mediate interferon-induced restriction. PLoS Pathog 18, e1010530. 10.1371/journal.ppat.1010530.

75. Goodier, J.L., Pereira, G.C., Cheung, L.E., Rose, R.J., and Kazazian, H.H., Jr. (2015). The Broad-Spectrum Antiviral Protein ZAP Restricts Human Retrotransposition. PLoS genetics 11, e1005252. 10.1371/journal.pgen.1005252.

76. Youn, J.Y., Dunham, W.H., Hong, S.J., Knight, J.D.R., Bashkurov, M., Chen, G.I., Bagci, H., Rathod, B., MacLeod, G., Eng, S.W.M., et al. (2018). High-Density Proximity Mapping Reveals the Subcellular Organization of mRNA-Associated Granules and Bodies. Mol Cell 69, 517–532 e511. 10.1016/j.molcel.2017.12.020.

77. Lau, N.C., Kolkman, A., van Schaik, F.M., Mulder, K.W., Pijnappel, W.W., Heck, A.J., and Timmers, H.T. (2009). Human Ccr4-Not complexes contain variable deadenylase subunits. Biochem J 422, 443–453. 10.1042/BJ20090500.

78. Yu, S., and Kim, V.N. (2020). A tale of non-canonical tails: gene regulation by post-transcriptional RNA tailing. Nat Rev Mol Cell Biol 21, 542–556. 10.1038/s41580-020-0246-8.

79. Chang, H.M., Triboulet, R., Thornton, J.E., and Gregory, R.I. (2013). A role for the Perlman syndrome exonuclease Dis3l2 in the Lin28-let-7 pathway. Nature 497, 244–248. 10.1038/nature12119.

80. Lubas, M., Damgaard, C.K., Tomecki, R., Cysewski, D., Jensen, T.H., and Dziembowski, A. (2013). Exonuclease hDIS3L2 specifies an exosome-independent 3’-5’ degradation pathway of human cytoplasmic mRNA. EMBO J 32, 1855–1868. 10.1038/emboj.2013.135.

81. Malecki, M., Viegas, S.C., Carneiro, T., Golik, P., Dressaire, C., Ferreira, M.G., and Arraiano, C.M. (2013). The exoribonuclease Dis3L2 defines a novel eukaryotic RNA degradation pathway. EMBO J 32, 1842–1854. 10.1038/emboj.2013.63.

82. Sharp, C.P., Thompson, B.H., Hoque, A.F., Diebold, O., Tesla, B., Kurian, D., Simmonds, P., Digard, P., and Gaunt, E. (2025). Understanding off-target growth defects introduced to influenza A virus by synonymous recoding. RNA 31, 1557–1574. 10.1261/rna.080675.125.

83. Wang, Y., Ma, T., He, Y., Li, Q., Mai, K., Mo, M., Cao, C., Li, J., Feng, P., Peng, J., et al. (2025). Genome-Wide Codon Reprogramming Enables a Multifactorially Attenuated Influenza Vaccine with Broad Cross-Protection. Adv Sci (Weinh), e16448. 10.1002/advs.202516448.

84. Fazal, F.M., Han, S., Parker, K.R., Kaewsapsak, P., Xu, J., Boettiger, A.N., Chang, H.Y., and Ting, A.Y. (2019). Atlas of Subcellular RNA Localization Revealed by APEX-Seq. Cell 178, 473–490 e426. 10.1016/j.cell.2019.05.027.

85. Schuhmacher, J.S., Tom Dieck, S., Christoforidis, S., Landerer, C., Davila Gallesio, J., Hersemann, L., Seifert, S., Schafer, R., Giner, A., Toth-Petroczy, A., et al. (2023). The Rab5 effector FERRY links early endosomes with mRNA localization. Mol Cell 83, 1839–1855 e1813. 10.1016/j.molcel.2023.05.012.

86. Horste, E.L., Fansler, M.M., Cai, T., Chen, X., Mitschka, S., Zhen, G., Lee, F.C.Y., Ule, J., and Mayr, C. (2023). Subcytoplasmic location of translation controls protein output. Mol Cell 83, 4509–4523 e4511. 10.1016/j.molcel.2023.11.025.

87. Bae, J.W., Kwon, S.C., Na, Y., Kim, V.N., and Kim, J.S. (2020). Chemical RNA digestion enables robust RNA-binding site mapping at single amino acid resolution. Nat Struct Mol Biol 27, 678–682. 10.1038/s41594-020-0436-2.

88. Todorova, T., Bock, F.J., and Chang, P. (2015). Poly(ADP-ribose) polymerase-13 and RNA regulation in immunity and cancer. Trends in molecular medicine 21, 373–384. 10.1016/j.molmed.2015.03.002.

89. Shaw, A.E., Hughes, J., Gu, Q., Behdenna, A., Singer, J.B., Dennis, T., Orton, R.J., Varela, M., Gifford, R.J., Wilson, S.J., and Palmarini, M. (2017). Fundamental properties of the mammalian innate immune system revealed by multispecies comparison of type I interferon responses. PLoS Biol 15, e2004086. 10.1371/journal.pbio.2004086.

90. Zheng, W., Guo, J., Ma, S., Sun, R., Song, Y., Chen, Y., Mao, R., and Fan, Y. (2024). The NEDD4-binding protein N4BP1 degrades mRNA substrates through the coding sequence independent of nonsense-mediated decay. J Biol Chem 300, 107954. 10.1016/j.jbc.2024.107954.

91. Tse, K.M., and Takeuchi, O. (2023). Innate immune sensing of pathogens and its post-transcriptional regulations by RNA-binding proteins. Arch Pharm Res 46, 65–77. 10.1007/s12272-023-01429-2.

92. Youle, R.L., Lista, M.J., Brudenell, E.L., Thompson, B., Bouton, C., Morris, E.R., Neil, S.J.D., Swanson, C.M., and Taylor, I.A. (2025). KHNYN is a manganese-dependent endoribonuclease required for ZAP-mediated antiviral restriction. Nucleic Acids Res 53. 10.1093/nar/gkaf1360.

93. Wang, C., Guan, Y., Lv, M., Zhang, R., Guo, Z., Wei, X., Du, X., Yang, J., Li, T., Wan, Y., et al. (2018). Manganese Increases the Sensitivity of the cGAS-STING Pathway for Double-Stranded DNA and Is Required for the Host Defense against DNA Viruses. Immunity 48, 675–687 e677. 10.1016/j.immuni.2018.03.017.

94. Yang, E., Huang, S., Jami-Alahmadi, Y., McInerney, G.M., Wohlschlegel, J.A., and Li, M.M.H. (2022). Elucidation of TRIM25 ubiquitination targets involved in diverse cellular and antiviral processes. PLoS Pathog 18, e1010743. 10.1371/journal.ppat.1010743.

95. Sanchez, J.G., Chiang, J.J., Sparrer, K.M.J., Alam, S.L., Chi, M., Roganowicz, M.D., Sankaran, B., Gack, M.U., and Pornillos, O. (2016). Mechanism of TRIM25 Catalytic Activation in the Antiviral RIG-I Pathway. Cell reports 16, 1315–1325. 10.1016/j.celrep.2016.06.070.

96. Choudhury, N.R., Heikel, G., Trubitsyna, M., Kubik, P., Nowak, J.S., Webb, S., Granneman, S., Spanos, C., Rappsilber, J., Castello, A., and Michlewski, G. (2017). RNA-binding activity of TRIM25 is mediated by its PRY/SPRY domain and is required for ubiquitination. BMC Biol 15, 105. 10.1186/s12915-017-0444-9.

97. Sanchez, J.G., Sparrer, K.M.J., Chiang, C., Reis, R.A., Chiang, J.J., Zurenski, M.A., Wan, Y., Gack, M.U., and Pornillos, O. (2018). TRIM25 Binds RNA to Modulate Cellular Anti-viral Defense. J Mol Biol 430, 5280–5293. 10.1016/j.jmb.2018.10.003.

98. Alvarez, L., Haubrich, K., Iselin, L., Gillioz, L., Ruscica, V., Lapouge, K., Augsten, S., Huppertz, I., Choudhury, N.R., Simon, B., et al. (2024). The molecular dissection of TRIM25’s RNA-binding mechanism provides key insights into its antiviral activity. Nature communications 15, 8485. 10.1038/s41467-024-52918-x.

99. Choudhury, N.R., Nowak, J.S., Zuo, J., Rappsilber, J., Spoel, S.H., and Michlewski, G. (2014). Trim25 Is an RNA-Specific Activator of Lin28a/TuT4-Mediated Uridylation. Cell reports 9, 1265–1272. 10.1016/j.celrep.2014.10.017.

100. Kwon, S.C., Yi, H., Eichelbaum, K., Fohr, S., Fischer, B., You, K.T., Castello, A., Krijgsveld, J., Hentze, M.W., and Kim, V.N. (2013). The RNA-binding protein repertoire of embryonic stem cells. Nat Struct Mol Biol 20, 1122–1130. 10.1038/nsmb.2638.

101. Manokaran, G., Finol, E., Wang, C., Gunaratne, J., Bahl, J., Ong, E.Z., Tan, H.C., Sessions, O.M., Ward, A.M., Gubler, D.J., et al. (2015). Dengue subgenomic RNA binds TRIM25 to inhibit interferon expression for epidemiological fitness. Science 350, 217–221. 10.1126/science.aab3369.

102. Choudhury, N.R., Trus, I., Heikel, G., Wolczyk, M., Szymanski, J., Bolembach, A., Dos Santos Pinto, R.M., Smith, N., Trubitsyna, M., Gaunt, E., et al. (2022). TRIM25 inhibits influenza A virus infection, destabilizes viral mRNA, but is redundant for activating the RIG-I pathway. Nucleic Acids Res 50, 7097–7114. 10.1093/nar/gkac512.

103. Le Pen, J., Jiang, H., Di Domenico, T., Kneuss, E., Kosalka, J., Leung, C., Morgan, M., Much, C., Rudolph, K.L.M., Enright, A.J., et al. (2018). Terminal uridylyltransferases target RNA viruses as part of the innate immune system. Nat Struct Mol Biol 25, 778–786. 10.1038/s41594-018-0106-9.

104. Huo, Y., Shen, J., Wu, H., Zhang, C., Guo, L., Yang, J., and Li, W. (2016). Widespread 3’-end uridylation in eukaryotic RNA viruses. Sci Rep 6, 25454. 10.1038/srep25454.

105. Warkocki, Z., Krawczyk, P.S., Adamska, D., Bijata, K., Garcia-Perez, J.L., and Dziembowski, A. (2018). Uridylation by TUT4/7 Restricts Retrotransposition of Human LINE-1s. Cell 174, 1537–1548 e1529. 10.1016/j.cell.2018.07.022.

106. Meze, K., Axhemi, A., Thomas, D.R., Doymaz, A., and Joshua-Tor, L. (2023). A shape-shifting nuclease unravels structured RNA. Nat Struct Mol Biol 30, 339–347. 10.1038/s41594-023-00923-x.

107. Watts, J.M., Dang, K.K., Gorelick, R.J., Leonard, C.W., Bess, J.W., Jr., Swanstrom, R., Burch, C.L., and Weeks, K.M. (2009). Architecture and secondary structure of an entire HIV-1 RNA genome. Nature 460, 711–716. 10.1038/nature08237.

108. Garcia-Moreno, M., Noerenberg, M., Ni, S., Jarvelin, A.I., Gonzalez-Almela, E., Lenz, C.E., Bach-Pages, M., Cox, V., Avolio, R., Davis, T., et al. (2019). System-wide Profiling of RNA-Binding Proteins Uncovers Key Regulators of Virus Infection. Mol Cell 74, 196–211 e111. 10.1016/j.molcel.2019.01.017.

109. Rozman, B., Nachshon, A., Levi Samia, R., Lavi, M., Schwartz, M., and Stern-Ginossar, N. (2022). Temporal dynamics of HCMV gene expression in lytic and latent infections. Cell reports 39, 110653. 10.1016/j.celrep.2022.110653.

110. Burgess, H.M., Grande, R., Riccio, S., Dinesh, I., Winkler, G.S., Depledge, D.P., and Mohr, I. (2023). CCR4-NOT differentially controls host versus virus poly(a)-tail length and regulates HCMV infection. EMBO Rep 24, e56327. 10.15252/embr.202256327.

111. Derdeyn, C.A., Decker, J.M., Sfakianos, J.N., Wu, X., O’Brien, W.A., Ratner, L., Kappes, J.C., Shaw, G.M., and Hunter, E. (2000). Sensitivity of human immunodeficiency virus type 1 to the fusion inhibitor T-20 is modulated by coreceptor specificity defined by the V3 loop of gp120. J Virol 74, 8358–8367. 10.1128/jvi.74.18.8358-8367.2000.

112. Wei, X., Decker, J.M., Liu, H., Zhang, Z., Arani, R.B., Kilby, J.M., Saag, M.S., Wu, X., Shaw, G.M., and Kappes, J.C. (2002). Emergence of resistant human immunodeficiency virus type 1 in patients receiving fusion inhibitor (T-20) monotherapy. Antimicrob Agents Chemother 46, 1896–1905. 10.1128/aac.46.6.1896-1905.2002.

113. Platt, E.J., Wehrly, K., Kuhmann, S.E., Chesebro, B., and Kabat, D. (1998). Effects of CCR5 and CD4 cell surface concentrations on infections by macrophagetropic isolates of human immunodeficiency virus type 1. J Virol 72, 2855–2864. 10.1128/JVI.72.4.2855-2864.1998.

114. Antzin-Anduetza, I., Mahiet, C., Granger, L.A., Odendall, C., and Swanson, C.M. (2017). Increasing the CpG dinucleotide abundance in the HIV-1 genomic RNA inhibits viral replication. Retrovirology 14, 49. 10.1186/s12977-017-0374-1.

115. Fouchier, R.A., Meyer, B.E., Simon, J.H., Fischer, U., and Malim, M.H. (1997). HIV-1 infection of non-dividing cells: evidence that the amino-terminal basic region of the viral matrix protein is important for Gag processing but not for post-entry nuclear import. EMBO J 16, 4531–4539. 10.1093/emboj/16.15.4531.

116. Chen, R.A., Ryzhakov, G., Cooray, S., Randow, F., and Smith, G.L. (2008). Inhibition of IkappaB kinase by vaccinia virus virulence factor B14. PLoS Pathog 4, e22. 10.1371/journal.ppat.0040022.

117. Lee, F.C.Y., Chakrabarti, A.M., Hänel, H., Monzón-Casanova, E., Hallegger, M., Militti, C., Capraro, F., Sadée, C., Toolan-Kerr, P., Wilkins, O., et al. (2021). An improved iCLIP protocol. bioRxiv, 2021.2008.2027.457890. 10.1101/2021.08.27.457890.

118. Smith, T., Heger, A., and Sudbery, I. (2017). UMI-tools: modeling sequencing errors in Unique Molecular Identifiers to improve quantification accuracy. Genome Res 27, 491–499. 10.1101/gr.209601.116.

119. Bushnell, B., Rood, J., and Singer, E. (2017). BBMerge - Accurate paired shotgun read merging via overlap. PLoS One 12, e0185056. 10.1371/journal.pone.0185056.

120. Ewels, P.A., Peltzer, A., Fillinger, S., Patel, H., Alneberg, J., Wilm, A., Garcia, M.U., Di Tommaso, P., and Nahnsen, S. (2020). The nf-core framework for community-curated bioinformatics pipelines. Nat Biotechnol 38, 276–278. 10.1038/s41587-020-0439-x.

121. West, C., Capitanchik, C., Cheshire, C., Luscombe, N.M., Chakrabarti, A., and Ule, J. (2023). nf-core/clipseq - a robust Nextflow pipeline for comprehensive CLIP data analysis. Wellcome Open Res 8, 286. 10.12688/wellcomeopenres.19453.1.

122. Di Tommaso, P., Chatzou, M., Floden, E.W., Barja, P.P., Palumbo, E., and Notredame, C. (2017). Nextflow enables reproducible computational workflows. Nat Biotechnol 35, 316–319. 10.1038/nbt.3820.

123. Frith, M.C., Valen, E., Krogh, A., Hayashizaki, Y., Carninci, P., and Sandelin, A. (2008). A code for transcription initiation in mammalian genomes. Genome Res 18, 1–12. 10.1101/gr.6831208.

124. Chakrabarti, A.M., Capitanchik, C., Ule, J., and Luscombe, N.M. (2023). clipplotr-a comparative visualization and analysis tool for CLIP data. RNA 29, 715–723. 10.1261/rna.079326.122.

125. Teufel, F., Almagro Armenteros, J.J., Johansen, A.R., Gislason, M.H., Pihl, S.I., Tsirigos, K.D., Winther, O., Brunak, S., von Heijne, G., and Nielsen, H. (2022). SignalP 6.0 predicts all five types of signal peptides using protein language models. Nat Biotechnol 40, 1023–1025. 10.1038/s41587-021-01156-3.

126. Stelzer, G., Rosen, N., Plaschkes, I., Zimmerman, S., Twik, M., Fishilevich, S., Stein, T.I., Nudel, R., Lieder, I., Mazor, Y., et al. (2016). The GeneCards Suite: From Gene Data Mining to Disease Genome Sequence Analyses. Current protocols in bioinformatics / editoral board, Andreas D. Baxevanis … [et al.] 54, 1 30 31–31 30 33. 10.1002/cpbi.5.

127. Yu, G., Wang, L.G., Han, Y., and He, Q.Y. (2012). clusterProfiler: an R package for comparing biological themes among gene clusters. Omics : a journal of integrative biology 16, 284–287. 10.1089/omi.2011.0118.

128. Chen, H., and Boutros, P.C. (2011). VennDiagram: a package for the generation of highly-customizable Venn and Euler diagrams in R. BMC bioinformatics 12, 35. 10.1186/1471-2105-12-35.

129. De Coster, W., D’Hert, S., Schultz, D.T., Cruts, M., and Van Broeckhoven, C. (2018). NanoPack: visualizing and processing long-read sequencing data. Bioinformatics 34, 2666–2669. 10.1093/bioinformatics/bty149.

130. Li, H. (2018). Minimap2: pairwise alignment for nucleotide sequences. Bioinformatics 34, 3094–3100. 10.1093/bioinformatics/bty191.

131. Li, H. (2021). New strategies to improve minimap2 alignment accuracy. Bioinformatics 37, 4572–4574. 10.1093/bioinformatics/btab705.

132. Lawrence, M., Huber, W., Pages, H., Aboyoun, P., Carlson, M., Gentleman, R., Morgan, M.T., and Carey, V.J. (2013). Software for computing and annotating genomic ranges. PLoS computational biology 9, e1003118. 10.1371/journal.pcbi.1003118.

