## Supplementary Figures for "The long isoform of ZAP coordinates multiple enzymes to mediate complete decay of target transcripts"

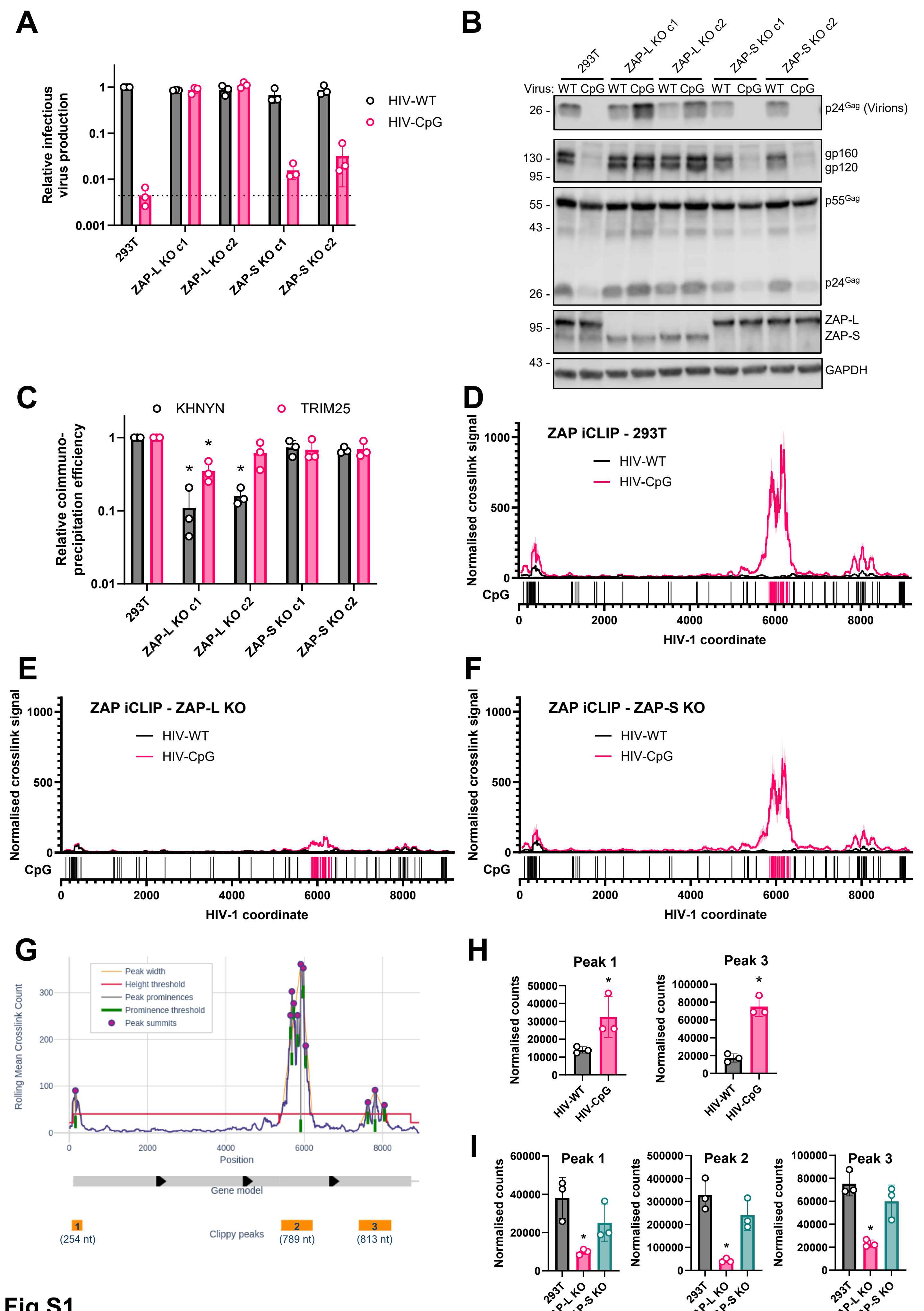
A
Relative infectious virus production
HIV-WT
HIV-CpG
293T
ZAP-L KO c1
ZAP-L KO c2
ZAP-S KO c1
ZAP-S KO c2
B
Virus: WT CpG WT CpG WT CpG WT CpG WT CpG
p24Gag (Virions)
gp160
gp120
p55Gag
p24Gag
ZAP-L
ZAP-S
GAPDH
C
KHNYN
TRIM25
Relative coimmuno-precipitation efficiency
D
ZAP iCLIP - 293T
Normalised crosslink signal
CpG
HIV-1 coordinate
E
ZAP iCLIP - ZAP-L KO
F
ZAP iCLIP - ZAP-S KO
G
Peak width
Height threshold
Peak prominences
Prominence threshold
Peak summits
Rolling Mean Crosslink Count
Position
Gene model
Clippy peaks
(254 nt)
(789 nt)
(813 nt)
H
Peak 1
Peak 3
Normalised counts
I
Peak 1
Peak 2
Peak 3
293T
ZAP-L KO
ZAP-S KO

Fig S1

**A**

| Motif | 293T rank | ZAP-S KO rank | ZAP-L KO rank |
|---|---|---|---|
| ACUACG | 1 | 1 | 27 |
| UUAUCG | 6 | 2 | 24 |
| CUACGU | 8 | 3 | 45 |
| CUACGA | 3 | 5 | 32 |
| UUUACG | 25 | 7 | 92 |
| UACGCC | 85 | 8 | 236 |
| UACGAC | 11 | 9 | 44 |
| UCUACG | 5 | 10 | 48 |
| UAACGU | 9 | 11 | 59 |
| UACGUG | 13 | 12 | 91 |
| UAUCGU | 17 | 13 | 34 |
| UUAACG | 102 | 14 | 197 |

**B**

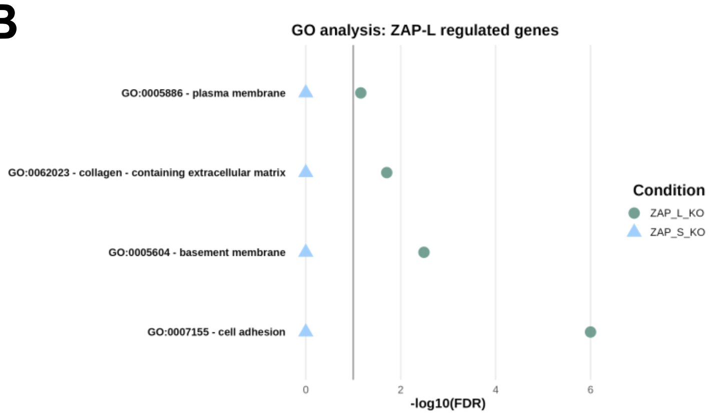


**C**

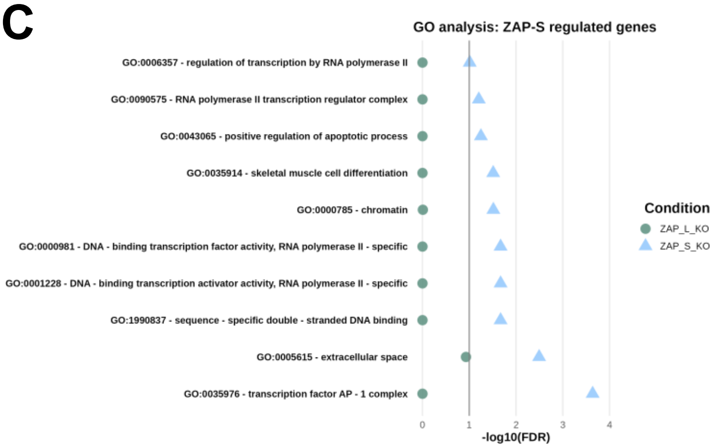


**D** **E** **F**

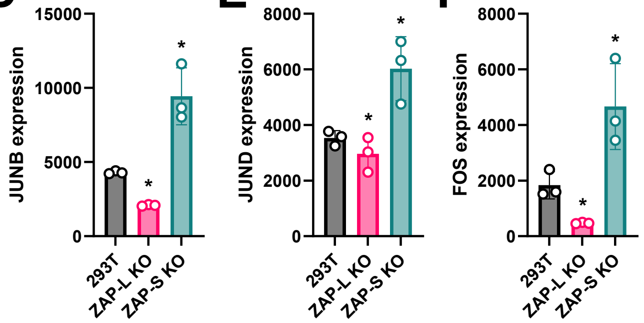


**G**

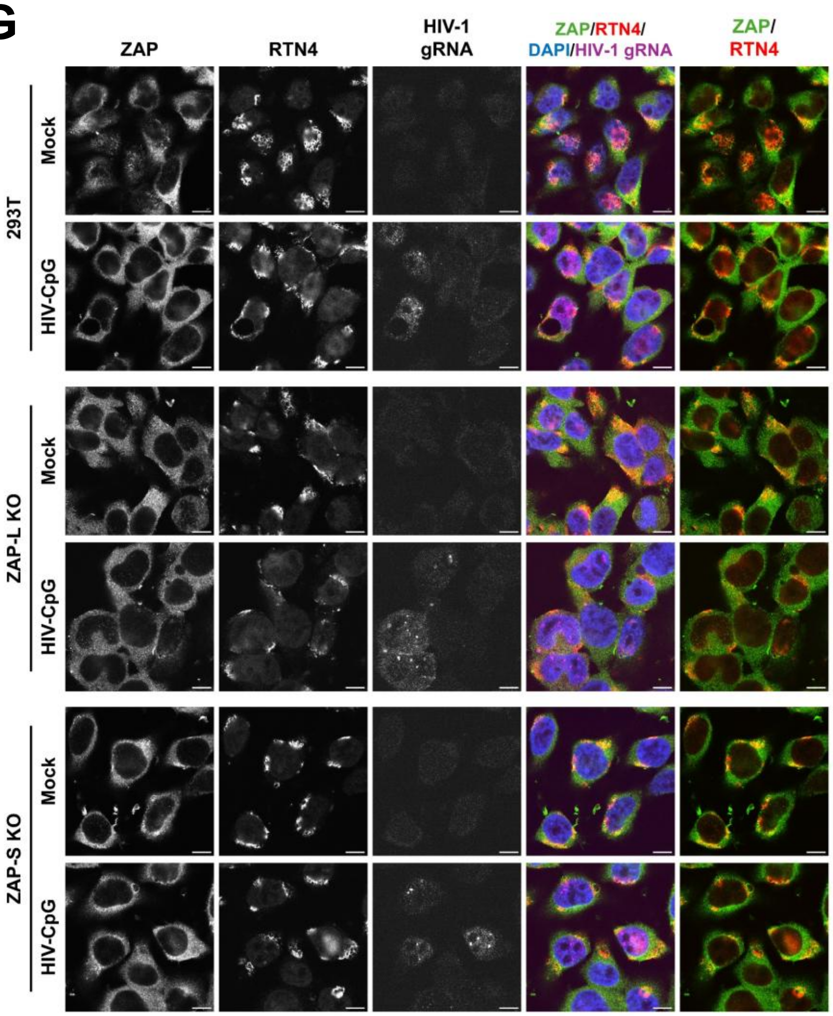


**Fig S2**

**TNFRSF10D Chr8:23137031-23137422 (-)strand**

```
         10        20        30        40        50        60        70        80        90       100
UUUUUUUCUGCUUCUUAUAUUGCAAGCUCCAUCUCUACUGGUGUGUGCAUUUAAUGACAUCUAACUACAGAUGCCGCACAGCCACAAUGCUUUGCCUUAU

        110       120       130       140       150       160       170       180       190       200
AGUUUUUUAACUUUAGAACGGGAUUAUCUUGUUAUUACCUGUAUUUUCAGUUUCGGAUAUUUUUGACUUAAUGAUGAGAUUAUCAAGACGUAGCCCUAUG

        210       220       230       240       250       260       270       280       290       300
CUAAGUCAUGAGCAUAUGGACUUACGAGGGUUCGACUUAGAGUUUUGAGCUUUAAGAUAGGAUUAUUGGGGCUUACCCCCACCUUAAUUAGAGAAACAUU

        310       320       330       340       350       360       370       380       390
UAUAUUGCUUACUACUGUAGGCUGUACAUCUCUUUUCCGAUUUUUGUAUAAUGAUGUAAACAUGGAAAAACUUUAGGAAAUGCACUUAUUA
```

**TNFRSF10D Chr8:23137558-23137753 (-)strand**

```
         10        20        30        40        50        60        70        80        90       100
AUAUGGGGCAGCUUACCAAUGGUCCUAGAACUUUGUUAACGCACUUGGAGUAAUUUUUAUGAAAUACUGCGUGUGAUAAGCAAACGGGAGAAAUUUAUAU

        110       120       130       140       150       160       170       180       190
CAGAUUCUUGGCUGCAUAGUUAUACGAUUGUGUAUUAAGGGUCGUUUUAGGCCACAUGCGGUGGCUCAUGCCUGUAAUCCCAGCACUUUGAUAGG
```

**Fig S3**

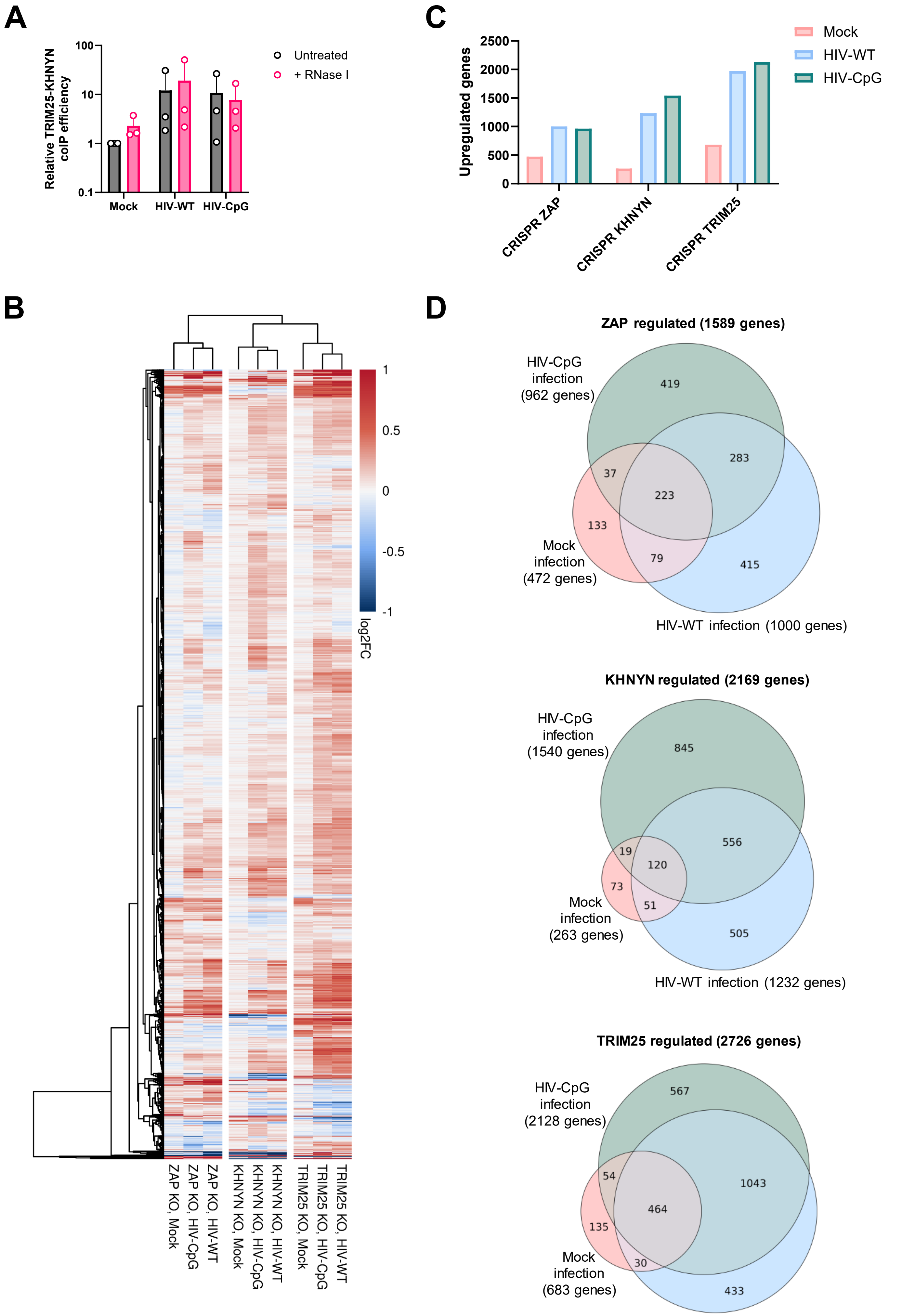
A
Relative TRIM25-KHNYN coIP efficiency
100
10
1
0.1
Untreated
+ RNase I
Mock
HIV-WT
HIV-CpG
B
1
0.5
0
-0.5
-1
log2FC
ZAP KO, Mock
ZAP KO, HIV-CpG
ZAP KO, HIV-WT
KHNYN KO, Mock
KHNYN KO, HIV-CpG
KHNYN KO, HIV-WT
TRIM25 KO, Mock
TRIM25 KO, HIV-CpG
TRIM25 KO, HIV-WT
C
Upregulated genes
2500
2000
1500
1000
500
0
Mock
HIV-WT
HIV-CpG
CRISPR ZAP
CRISPR KHNYN
CRISPR TRIM25
D
ZAP regulated (1589 genes)
HIV-CpG infection (962 genes)
419
283
37
223
133
79
415
Mock infection (472 genes)
HIV-WT infection (1000 genes)
KHNYN regulated (2169 genes)
HIV-CpG infection (1540 genes)
845
556
19
120
73
51
505
Mock infection (263 genes)
HIV-WT infection (1232 genes)
TRIM25 regulated (2726 genes)
HIV-CpG infection (2128 genes)
567
1043
54
464
135
30
433
Mock infection (683 genes)
HIV-WT infection (1970 genes)

Fig S4

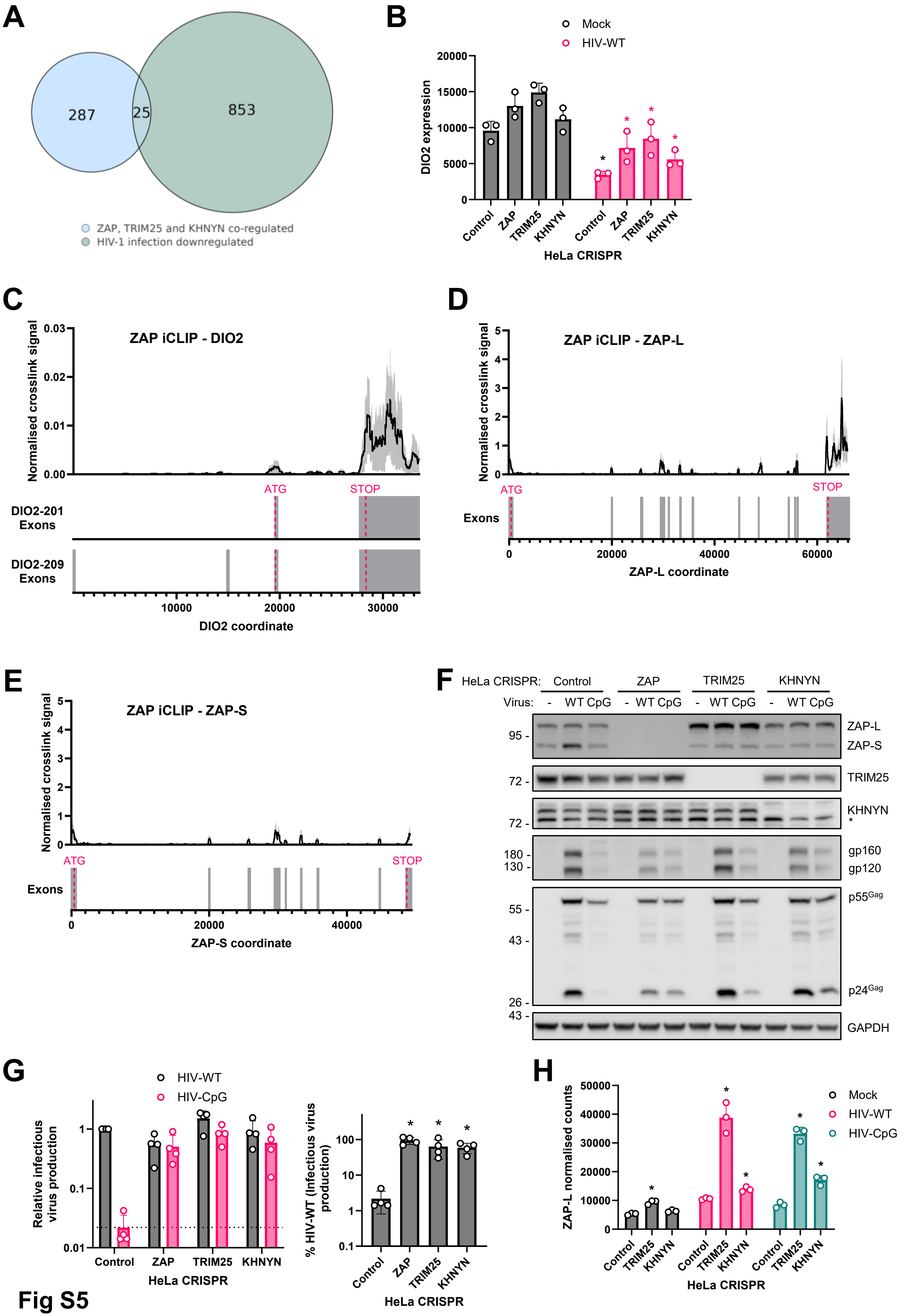
A
287
25
853
ZAP, TRIM25 and KHNYN co-regulated
HIV-1 infection downregulated
B
Mock
HIV-WT
DIO2 expression
Control
ZAP
TRIM25
KHNYN
HeLa CRISPR
C
ZAP iCLIP - DIO2
Normalised crosslink signal
ATG
STOP
DIO2-201 Exons
DIO2-209 Exons
DIO2 coordinate
D
ZAP iCLIP - ZAP-L
Exons
ZAP-L coordinate
E
ZAP iCLIP - ZAP-S
ZAP-S coordinate
F
HeLa CRISPR: Control ZAP TRIM25 KHNYN
Virus: - WT CpG
ZAP-L
ZAP-S
TRIM25
KHNYN
gp160
gp120
p55Gag
p24Gag
GAPDH
G
HIV-WT
HIV-CpG
Relative infectious virus production
% HIV-WT (Infectious virus production)
H
ZAP-L normalised counts
HIV-CpG

Fig S5

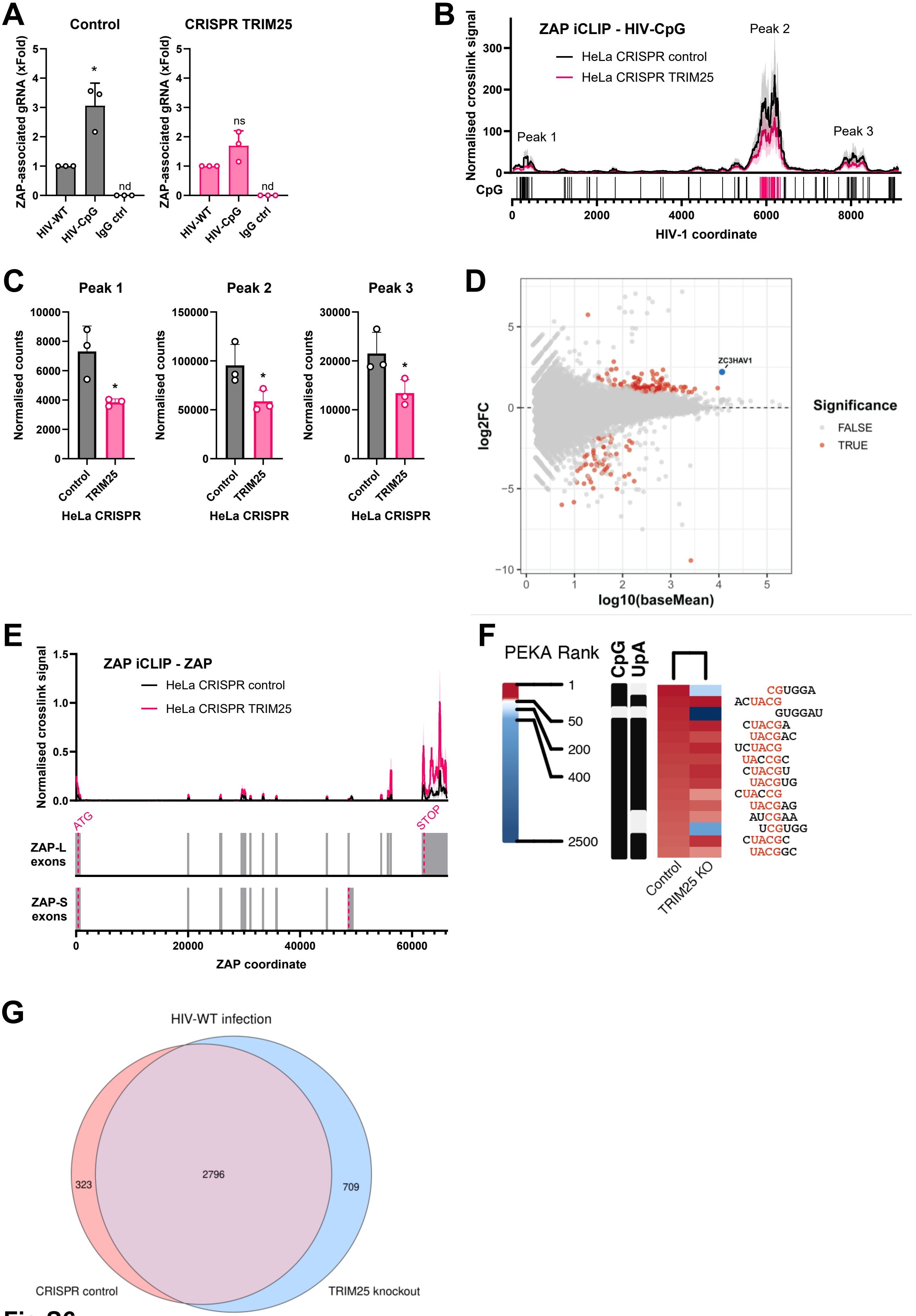
A
Control
CRISPR TRIM25
ZAP-associated gRNA (xFold)
HIV-WT
HIV-CpG
IgG ctrl
nd
ns
B
ZAP iCLIP - HIV-CpG
HeLa CRISPR control
HeLa CRISPR TRIM25
Normalised crosslink signal
Peak 1
Peak 2
Peak 3
CpG
HIV-1 coordinate
C
Peak 1
Peak 2
Peak 3
Normalised counts
Control
TRIM25
HeLa CRISPR
D
ZC3HAV1
log2FC
log10(baseMean)
Significance
FALSE
TRUE
E
ZAP iCLIP - ZAP
HeLa CRISPR control
HeLa CRISPR TRIM25
Normalised crosslink signal
ATG
STOP
ZAP-L exons
ZAP-S exons
ZAP coordinate
F
PEKA Rank
CpG
UpA
Control
TRIM25 KO
CGUGGA
ACUACG
GUGGAU
CUACGA
UACGAC
UCUACG
UACCGC
CUACGU
UACGUG
CUACCG
UACGAG
AUCGAA
UCGUGG
CUACGC
UACGGC
G
HIV-WT infection
323
2796
709
CRISPR control
TRIM25 knockout

**Fig S6**

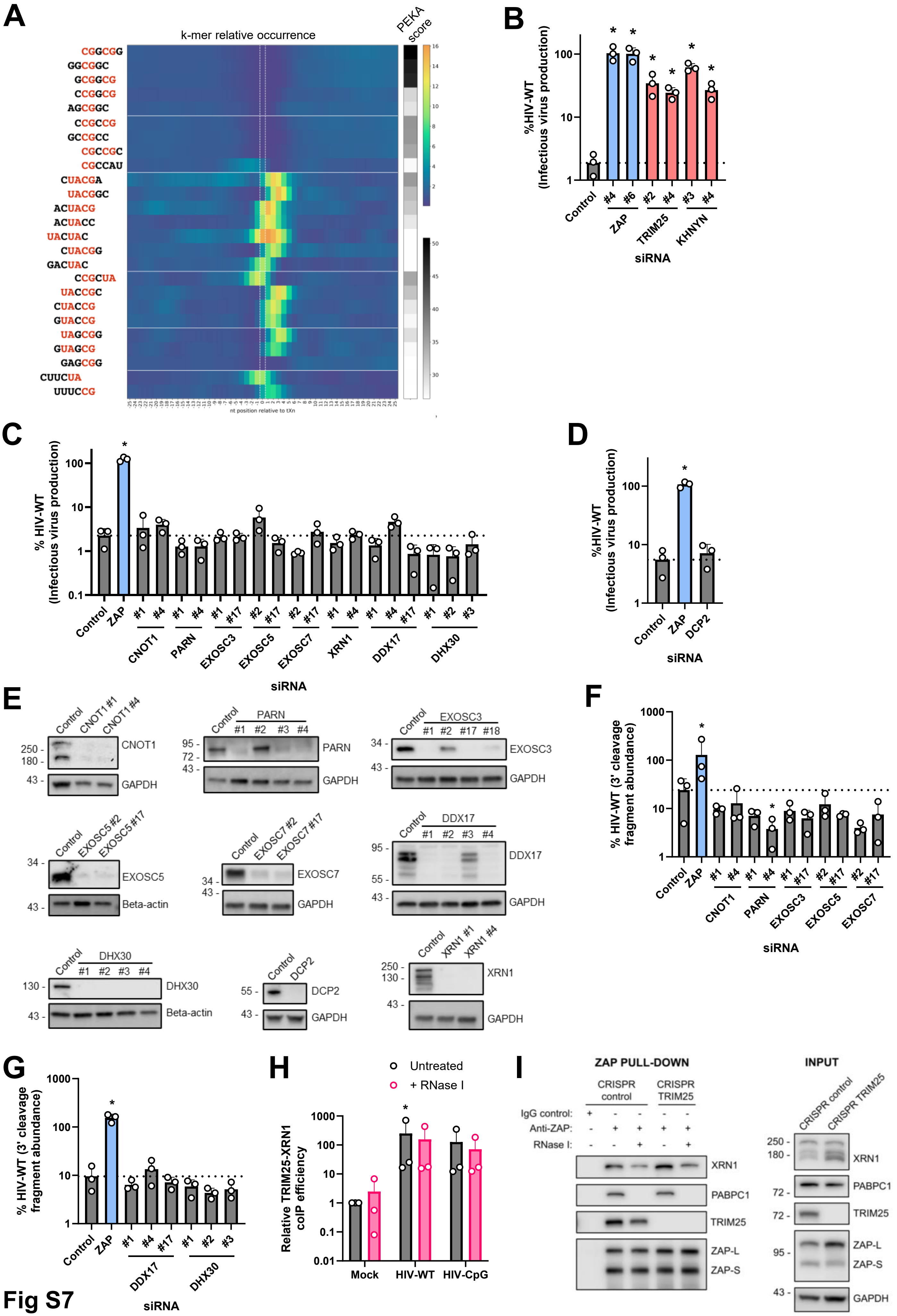
A
k-mer relative occurrence
PEKA score
CGGCGG
GGCGGC
GCGGCG
CCGGCG
AGCGGC
CCGCCG
GCCGCC
CGCCGC
CGCCAU
CUACGA
UACGGC
ACUACG
ACUACC
UACUAC
CUACGG
GACUAC
CCGCUA
UACCGC
CUACCG
GUACCG
UAGCGG
GUAGCG
GAGCGG
CUUCUA
UUUCCG
nt position relative to tXn
B
%HIV-WT (Infectious virus production)
Control
ZAP
TRIM25
KHNYN
siRNA
C
% HIV-WT (Infectious virus production)
Control
ZAP
CNOT1
PARN
EXOSC3
EXOSC5
EXOSC7
XRN1
DDX17
DHX30
siRNA
D
%HIV-WT (Infectious virus production)
Control
ZAP
DCP2
siRNA
E
CNOT1
PARN
EXOSC3
EXOSC5
EXOSC7
DDX17
DHX30
DCP2
XRN1
GAPDH
Beta-actin
F
% HIV-WT (3' cleavage fragment abundance)
siRNA
G
% HIV-WT (3' cleavage fragment abundance)
siRNA
H
Untreated
+ RNase I
Relative TRIM25-XRN1 coIP efficiency
Mock
HIV-WT
HIV-CpG
I
ZAP PULL-DOWN
CRISPR control
CRISPR TRIM25
IgG control:
Anti-ZAP:
RNase I:
INPUT
CRISPR control
CRISPR TRIM25
XRN1
PABPC1
TRIM25
ZAP-L
ZAP-S
GAPDH

Fig S7

# A

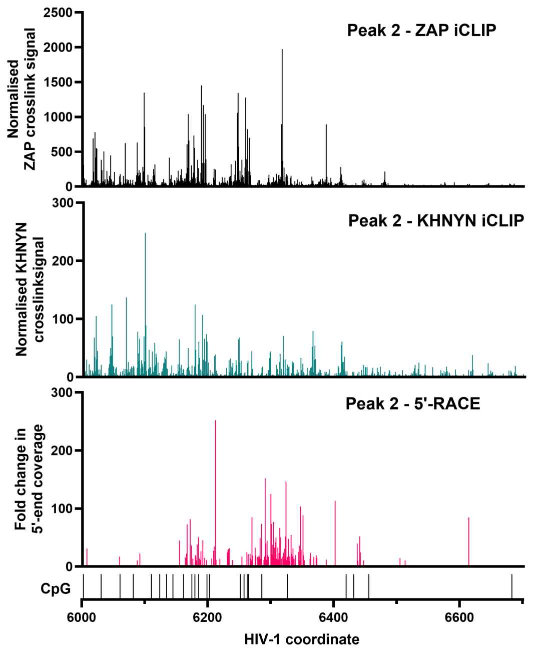


# B

6000 6010 6020 6030 6040 6050 6060 6070 6080 6090

ACCCGAA/CCCACAAGAAGUAGUAUUGGUAAACGUGACAGAAAAUUUUAACAUGUGGAAA/AACGACAUGGUAGAACAGAUGCACGAGG/AUAU/AAUCAGUUU

6100 6110 6120 6130 6140 6150 6160 6170 6180 6190

AUGGGAUCAAUCGCUAAAGCCAUGCGUAAAAUUAACGCCACUCUGCGUUAGUUU/AAAGUGCAC/GG/A/UUUGA/AG/AA/CGAU/A/C/G/AA/UA/C/G/AAU/AGU/AGU/AGC

6200 6210 6220 6230 6240 6250 6260 6270 6280 6290

GGGCG/CAU/G/AU/A/AUGGAG/AAAGGAGAGAUA/A/A/A/AACUG/CUCUUUCAAUAUCUC/GACAUCGA/UA/CGC/G/A/U/A/AGGU/G/CAG/A/A/A/G/A/A/UACG/C/A/U/U/C/U/U/UUAU/A

6300 6310 6320 6330 6340 6350 6360 6370 6380 6390

/A/A/C/U/U/G/A/U/A/U/A/G/U/A/CC/AA/U/A/G/A/U/A/A/U/A/C/G/AG/C/U/A/U/A/GG/UUGAU/A/A/G/U/UGU/A/A/CACCUCAGU/C/AUUAC/ACAG/G/CCUGUCCAAAGGUAU/CCUUUGAGCCAAU

6400 6410 6420 6430 6440 6450 6460 6470 6480 6490

U/CCCAUACAUUAUUGUGCCCCGGCUGGUUUUGCGAU/U/CUA/A/AAUGU/AAUAAUAAGACGUUCAAUGGAACAGGACCAUGUACAAAUGUCAGCACAGUACAA

6500 6510 6520 6530 6540 6550 6560 6570 6580 6590

UGUA/CACAUGGA/AUCAGGCCAGUAGUAUCAACUCAACUGCUGUUAAAUGGCAGUCUAGCAGAAGAAGAUGUAGUAAUUAGAUCUGCCAAUUUCACAGACA

6600 6610 6620 6630 6640 6650 6660 6670 6680 6690

AUGCUAAAACCAU/AAUAGUACAGCUGAACACAUCUGUAGAAAUUAAUUGUACAAGACCCAACAACAAUACAAGAAAAAGUAUCCGUAUCCAGAGGGGACC

### Fig S8

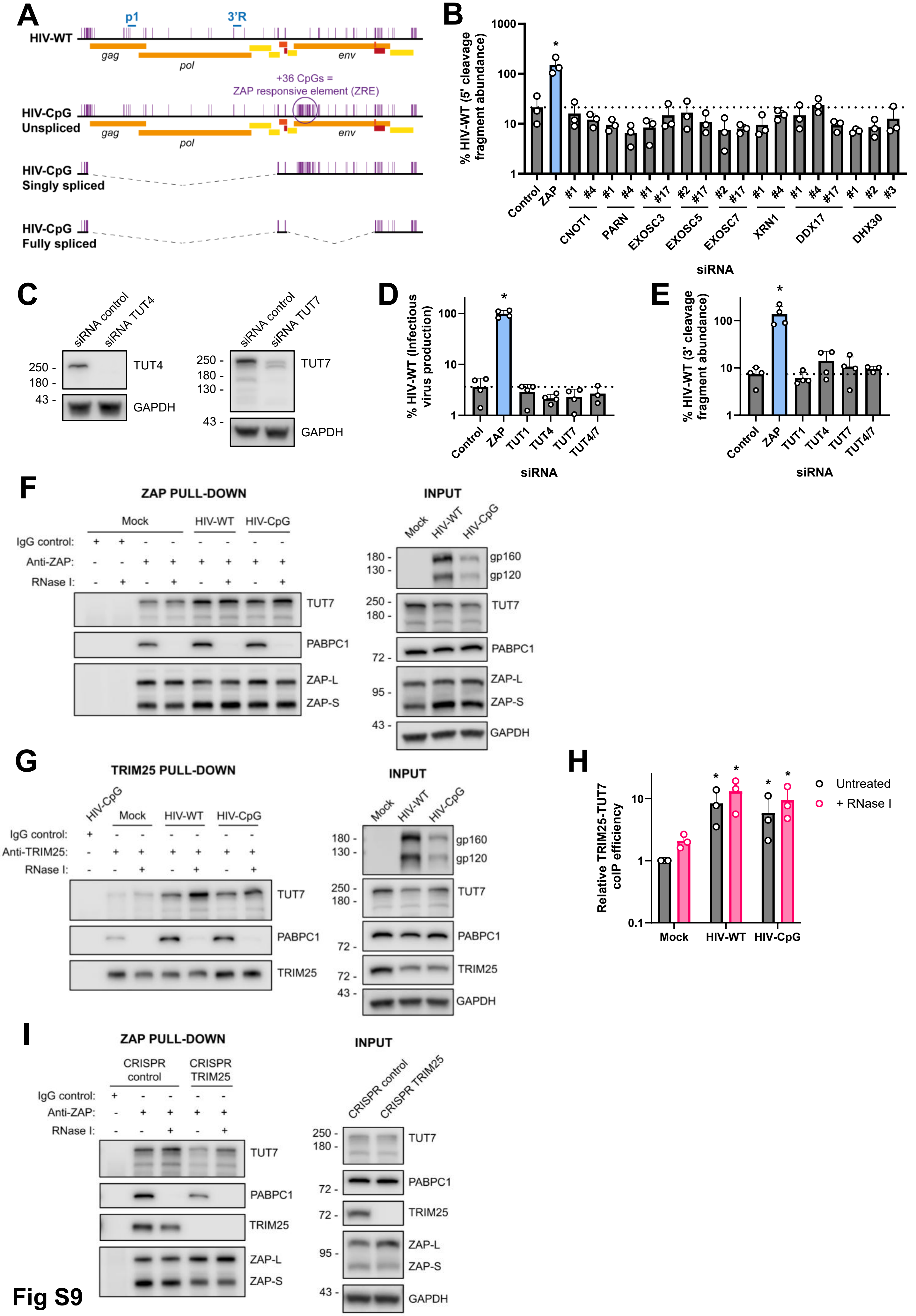
A
p1
3'R
HIV-WT
gag
pol
env
+36 CpGs =
ZAP responsive element (ZRE)
HIV-CpG
Unspliced
HIV-CpG
Singly spliced
HIV-CpG
Fully spliced
B
% HIV-WT (5' cleavage fragment abundance)
Control
ZAP
CNOT1
PARN
EXOSC3
EXOSC5
EXOSC7
XRN1
DDX17
DHX30
siRNA
C
siRNA control
siRNA TUT4
TUT4
GAPDH
siRNA TUT7
TUT7
D
% HIV-WT (Infectious virus production)
TUT1
TUT4
TUT7
TUT4/7
E
% HIV-WT (3' cleavage fragment abundance)
F
ZAP PULL-DOWN
Mock
HIV-WT
HIV-CpG
IgG control:
Anti-ZAP:
RNase I:
PABPC1
ZAP-L
ZAP-S
INPUT
gp160
gp120
G
TRIM25 PULL-DOWN
Anti-TRIM25:
TRIM25
H
Relative TRIM25-TUT7 coIP efficiency
Untreated
+ RNase I
I
CRISPR control
CRISPR TRIM25

Fig S9

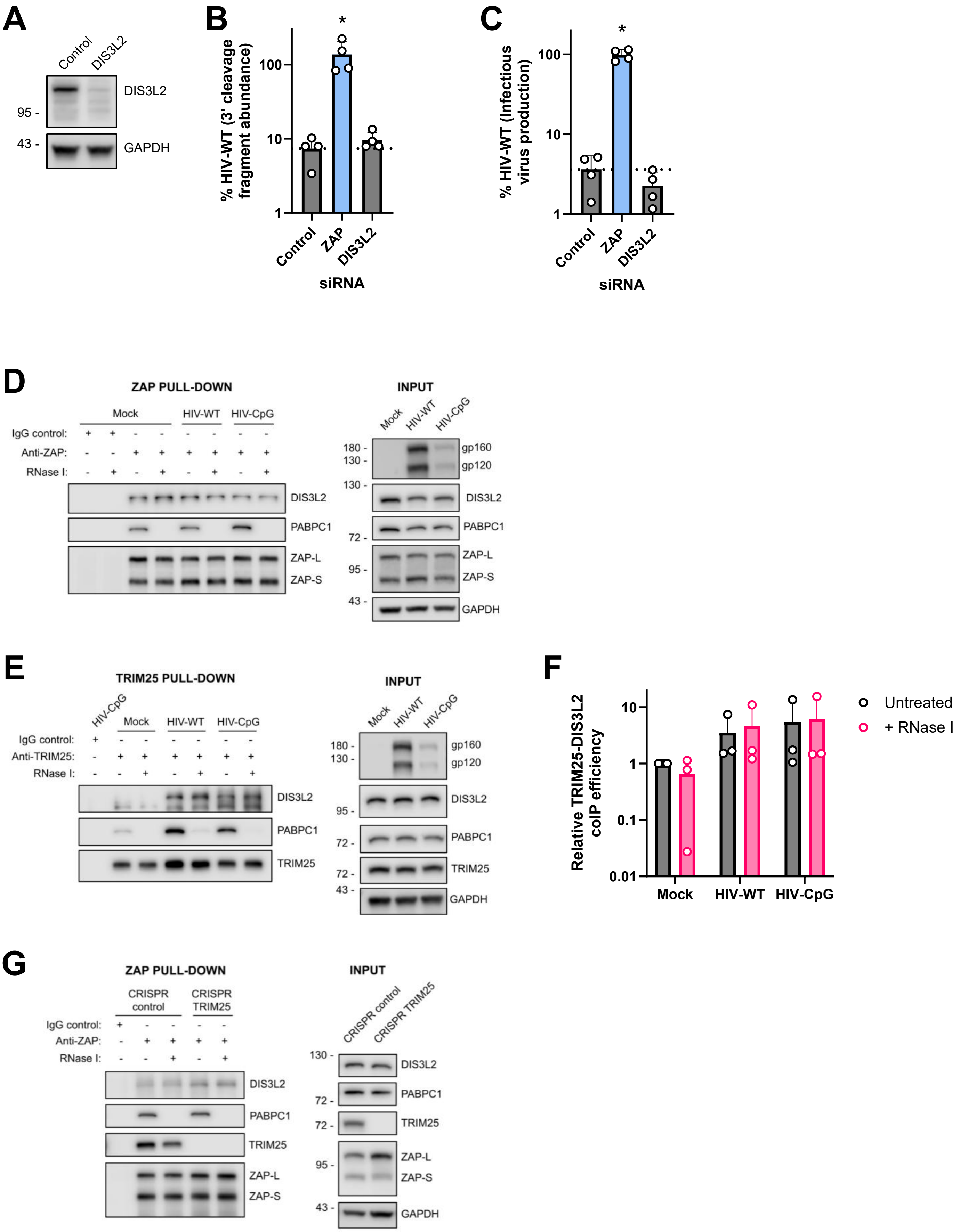
A
Control
DIS3L2
DIS3L2
95 -
43 -
GAPDH
B
% HIV-WT (3' cleavage fragment abundance)
*
100
10
1
Control
ZAP
DIS3L2
siRNA
C
% HIV-WT (Infectious virus production)
*
100
10
1
Control
ZAP
DIS3L2
siRNA
D
ZAP PULL-DOWN
Mock
HIV-WT
HIV-CpG
IgG control:
Anti-ZAP:
RNase I:
DIS3L2
PABPC1
ZAP-L
ZAP-S
INPUT
Mock
HIV-WT
HIV-CpG
180 -
130 -
gp160
gp120
130 -
DIS3L2
PABPC1
72 -
ZAP-L
95 -
ZAP-S
43 -
GAPDH
E
TRIM25 PULL-DOWN
HIV-CpG
Mock
HIV-WT
HIV-CpG
IgG control:
Anti-TRIM25:
RNase I:
DIS3L2
PABPC1
TRIM25
INPUT
Mock
HIV-WT
HIV-CpG
180 -
130 -
gp160
gp120
DIS3L2
95 -
PABPC1
72 -
TRIM25
72 -
43 -
GAPDH
F
Relative TRIM25-DIS3L2 coIP efficiency
10
1
0.1
0.01
Mock
HIV-WT
HIV-CpG
Untreated
+ RNase I
G
ZAP PULL-DOWN
CRISPR control
CRISPR TRIM25
IgG control:
Anti-ZAP:
RNase I:
DIS3L2
PABPC1
TRIM25
ZAP-L
ZAP-S
INPUT
CRISPR control
CRISPR TRIM25
130 -
DIS3L2
PABPC1
72 -
72 -
TRIM25
95 -
ZAP-L
ZAP-S
43 -
GAPDH

**Fig S10**
